# High-efficiency, transgene-free plant genome editing by viral delivery of an engineered TnpB

**DOI:** 10.64898/2025.12.05.692691

**Authors:** Ugrappa Nagalakshmi, Jorge E. Rodriguez, Thi Nguyen, Rachel F. Weissman, Brittney W. Thornton, David F. Savage, Savithramma P Dinesh-Kumar

## Abstract

Genome editing has revolutionized plant biology research. However, efficient and straightforward delivery of editing reagents remains a major challenge. Viral delivery systems can address these issues, but CRISPR-Cas nucleases are often too large for viral vectors. Recently, smaller editors like TnpBs have been identified, but wild-type TnpBs are significantly less active than commonly used Cas9 nucleases. Here, we optimized a tobacco rattle virus (TRV)-based system to deliver newly discovered, highly active engineered ISDra2 TnpB variants. Our results demonstrate that the eTnpBc variant delivered via TRV enables effective somatic editing in systemic leaves and achieves up to 90% editing efficiency at target loci, significantly higher than that of wild-type ISDra2 TnpB. Additionally, up to 89% of offspring exhibit a mutant phenotype, with editing efficiencies reaching 100%. The design principles outlined here are expected to accelerate broader adoption of eTnpBc for transformation- and transgene-free genome editing in plants.

## Main

The development of programmable RNA-guided endonucleases, such as CRISPR-Cas9, has revolutionized genome editing in plants^1^. However, the efficient delivery of gene editing components into plant cells remains a significant challenge. Current methods rely on a labor-intensive, lengthy tissue culture process involving complex transformation and regeneration steps, which can also introduce unintentional changes to the genome and epigenome. To overcome these challenges, an alternative approach uses engineered plant viral vectors to deliver gene editing components^2^. Because most plant viruses have a limited cargo capacity insufficient to accommodate CRISPR-Cas effectors, they have primarily been used to deliver guide RNAs (gRNAs) into Cas9-expressing transgenic plants^2–11^. A major limitation of this method is the need to generate an initial transgenic expressing a nuclease, such as Cas9. Although many plant viral vectors can deliver gRNA to induce somatic editing, only a few have been shown to produce heritable edits in plants^3–7,9–12^. Some engineered plant negative-strand rhabdoviruses can deliver both Cas9 and gRNA, but they face other obstacles, such as the need for tissue regeneration or pruning of infected plants^13–16^. Additionally, delivery of most rhabdoviruses is feasible only through vector transmission, which requires specialized facilities and containment^16^.

The delivery and cargo limits of mammalian and plant viral vectors have increased interest in smaller RNA-guided endonucleases, such as TnpBs (∼400 amino acids), as alternatives to commonly used CRISPR-Cas proteins (>1000 amino acids) for genome editing^17–19^. TnpBs are proposed evolutionary ancestors of Type V CRISPR-Cas systems and form a ribonucleoprotein (RNP) complex with a programmable omega (obligate mobile element-guided activity) RNA (ωRNA), also known as reRNA (right-end element RNA)^17,18^.

Among TnpBs, ISDra2 TnpB from *Deinococcus radiodurans* was one of the first to be experimentally characterized^17^. ISDra2 requires a 5’-TTGAT as a transposon-associated motif (TAM) for target recognition and cleavage, which is functionally similar to the protospacer-adjacent motif (PAM) of Cas9^17^. ISDra2 has been shown to enable genome editing in transgenic plants^20–22^. The small size of ISDra2 TnpB makes it ideal for virus-based delivery into plants, facilitating transgene-free editing. Moreover, ISYmu1 TnpB delivered via a viral vector in Arabidopsis demonstrated very low somatic (3-8%) and heritable (∼3%) editing efficiency^23^. Additionally, TRV-based delivery of another compact Cas nuclease, AsCas12f, has been shown to induce editing efficiency of ∼6%^24^. Therefore, we tested whether our recently engineered, highly active ISDra2 TnpB variants, generated through deep mutational scanning^25^ and delivered via our well-established tobacco rattle virus (TRV) system, could induce high levels of somatic and heritable editing in plants.

TRV is a bipartite, positive-sense RNA virus consisting of RNA1 and RNA2^26^. We previously engineered RNA1 (TRV1) and RNA2 (TRV2) vectors, which can be delivered to plants via *Agrobacterium tumefaciens*-mediated infiltration^27,28^ (Fig. 1a). To express RNA or protein-coding sequences (cargo), we incorporated the pea early browning virus coat protein promoter (pPEBV) into TRV2^27,29^ (Fig. 1a). We expressed wild-type (WT) TnpB using the TRV system by cloning *Nicotiana benthamiana* codon-optimized (NbCo) ISDra2 TnpB, fused to a duplicated Simian Virus 40 (SV40) nuclear localization sequence (NLS) at the C-terminus (NbCoTnpB_2xSV40nls_) into TRV2 downstream of pPEBV (Extended Data Fig. 1a). We added the mobile tRNA isoleucine (tRNA^Ile^) sequence downstream of the TnpB coding sequence because, as we previously demonstrated, adding this sequence improves virus systemic movement and enhances somatic and heritable editing efficiency when delivering gRNA into Cas9-expressing plants^3,6^. We then generated TRV2 constructs with reRNA targeting all four TAM sites within the *N. benthamiana Phytoene desaturase* (*PDS*) genes, homeologs Nbe06g25970 and Nbe05g35010^30^ (Extended Data Fig. 1a and Extended Data Fig. 2a). *PDS* is critical for carotenoid pigment biosynthesis, and disrupting *PDS* causes chlorophyll breakdown, resulting in a photobleaching phenotype. Agrobacterium carrying TRV1, TRV2-NbCoTnpB_2xSV40nls_, and TRV2 vectors with individual *PDS* guides (reRNA1 to reRNA4) was infiltrated into the leaves of 2.5-week-old *N. benthamiana* plants. After 4-5 weeks post-infiltration, we observed no photobleaching phenotype in systemic leaves with any of the *PDS*-targeting guides (Extended Data Fig. 1a). Amplicon sequencing revealed a very low indel frequency in the infiltrated (0.01 to 1.47%; Extended Data Fig. 1b and Supplementary Table 1) and systemic leaves (0.02 to 0.23%; Extended Data Fig. 1c and Supplementary Table 2). This suggests that the lack of an observable photobleaching phenotype in these plants could be due to a low editing activity of WT TnpB.

**Figure 1.**
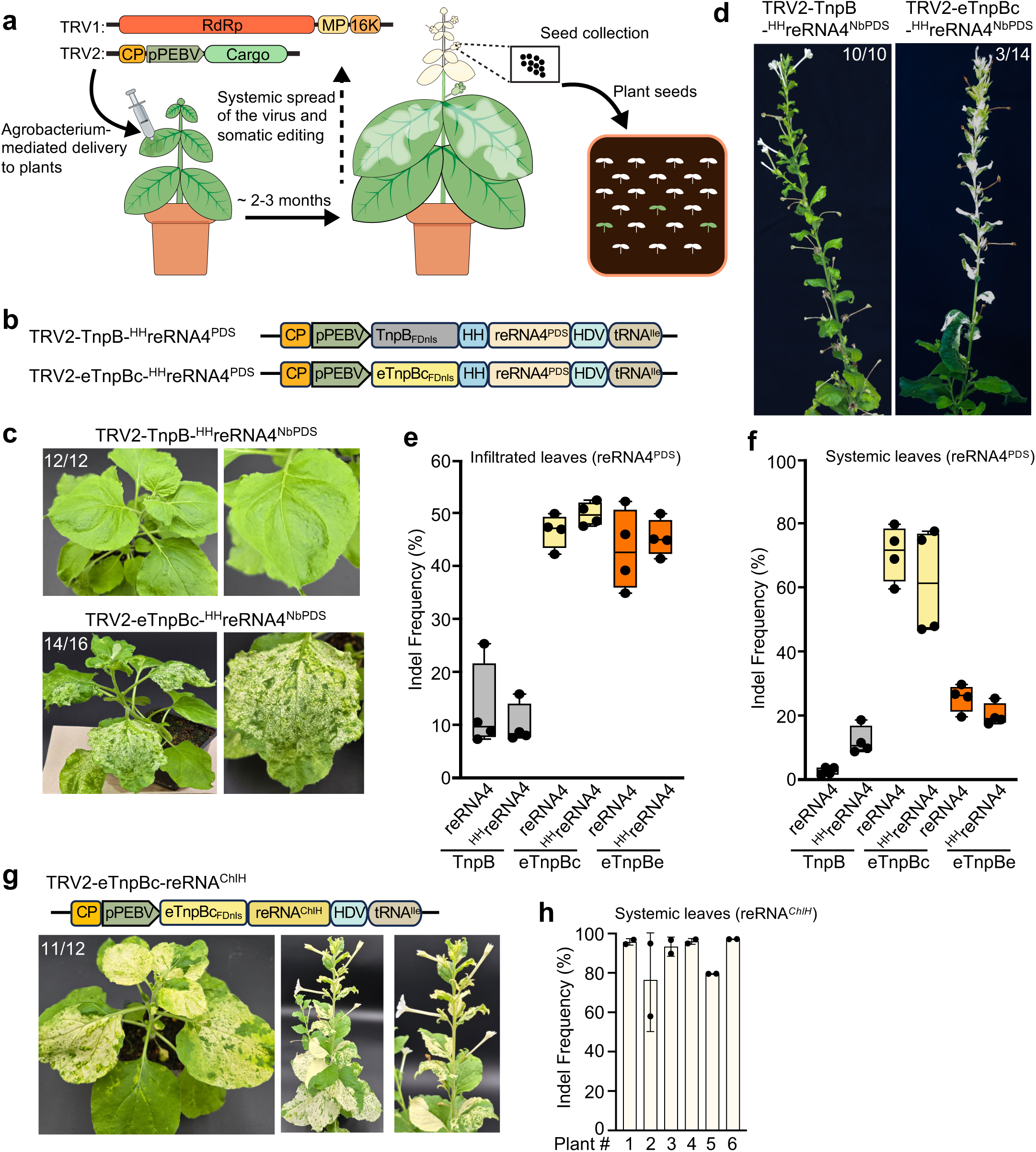
Virus delivery of the enhanced eTnpBc variant results in efficient systemic somatic editing. **a,** Overview of the viral delivery method. Agrobacterium carrying the tobacco rattle virus (TRV) vector with TnpB and reRNA (cargo) is injected into 2.5-week-old plants using a needleless syringe (left panel). The somatic editing phenotype appears as the virus spreads throughout the plants (middle panel). Pods collected from the top third of mature plants and seeds are planted, screened for mutant traits, and evaluated for editing efficiency (right panel). RdRp, RNA-dependent RNA polymerase; MP, movement protein; 16K, RNA silencing suppressor; CP, coat protein; pPEBV, pea early browning virus subgenomic promoter. Refer to methods for details. **b,** Diagram showing TRV2 vectors with wild-type TnpB and the eTnpBc variant fused to the nuclear localization sequence of Arabidopsis FD protein (FDnls), positioned downstream of pPEBV. reRNA4 targeting two *PDS* homeologs in *Nicotiana benthamiana* is flanked by self-cleaving hammerhead (HH) and hepatitis delta virus (HDV) ribozymes (^HH^reRNA4^PDS^). The tRNA isoleucine sequence (tRNA^Ile^) is located at the 3’ end to enhance virus mobility. **c,** Phenotypes of plants infected with TRV carrying wild-type TnpB (top panels) and eTnpBc (bottom panels) with ^HH^reRNA4^PDS^. Plants were photographed approximately 2.5 weeks after TRV infiltration. The white sectors on the leaves, which appear bleached, indicate the loss of *PDS* function. The inset in the top left corner of the photo indicates the number of plants showing the corresponding phenotype. n=≥3 plants; 4 independent experiments. **d,** Photos of mature plants showing complete photobleaching of the terminal stem, including leaves and flowers, in plants infiltrated with eTnpBc (right panel) compared to TnpB (left panel). The inset in the top right corner of the photo indicates the number of plants showing the corresponding phenotype. **e-f,** Indel frequency of TnpB and TnpB variants (eTnpBc and eTnpBe) with two designs of reRNA4 targeting the *PDS* locus, reRNA4 without HH ribozyme at the 5’ end (reRNA4^PDS^), and ^HH^reRNA4^PDS^ assessed in the infiltrated leaves (**e**) and systemic leaves (**f**) by amplicon sequencing as described in methods. Data is from biological replicates (n=4). The box shows the interquartile range (Q1 to Q3), the line inside indicates the median, and the whiskers extend to the most extreme points within 1.5× IQR of the quartiles. **g,** Somatic editing phenotype of *ChlH*. Phenotype of plants infiltrated with TRV expressing eTnpBc with reRNA targeting two *ChlH* homeologs in *N. benthamiana* (reRNA^ChlH^). Plants were photographed approximately 2.5 weeks (left panel) and 5 weeks (middle panel) after TRV infiltration. The right panel is an enlarged top view of the middle panel. The yellow coloration on the leaves indicates the loss of *ChlH* function. The inset in the top left corner of the photo indicates the number of plants showing the corresponding phenotype. n=4 plants; 3 independent experiments. **h,** Indel frequency at the *ChlH* locus in systemic leaves. Data are plotted as the mean and standard deviation (SD) from biological replicates (n=6); two tissue samples from each plant.

Recently, we used deep mutational scanning of ISDra2 TnpB and reRNA to identify improved variants that significantly enhance editing efficiency compared to the WT ISDra2 TnpB in yeast, human cells, *N. benthamiana* and pepper leaves, and rice calli^25^. Therefore, we tested the two most active enhanced variants, eTnpBc (N4Y/R110K/V192L/L222I) and eTnpBe (L172G/V192L/L222I/P282V/I304R), against the WT TnpB, using the TRV delivery system (Fig. 1a). For TnpB and its variants, we used coding sequences described previously^25^, which were codon-optimized for expression in both human and yeast cells, unless otherwise noted. We fused the TnpBs to the NLS derived from the Arabidopsis Flowering Locus D (FD) transcription factor, which functions better in plant cells than SV40 NLS^31^ (Extended Data Fig. 3a). We used the reRNA4 site for these experiments because it exhibits increased editing activity in *N. benthamiana* leaves when expressed using the U6 promoter^25^ and when expressed with WT TnpB using TRV2 (Extended Data Fig. 1b and Supplementary Table 1). To test whether the precise 3’ and 5’ ends of the reRNA are essential for optimal editing, we introduced a self-cleaving hepatitis delta virus (HDV) ribozyme at the 3’ end of the reRNA, either alone or together with a hammerhead (HH) ribozyme at the 5’ end (referred to as reRNA4^PDS^ and ^HH^reRNA4^PDS^, respectively; Extended Data Fig. 3a). Although infiltration of TRV with WT TnpB and enhanced variants with both reRNA4 designs, failed to induce the photobleaching phenotype in the systemic leaves of *N. benthamiana* plants (Extended Data Fig. 3b), we observed significantly higher indel frequencies in infiltrated leaves with the enhanced variants eTnpBc (14.4±3 to 18.6±1.7%) and eTnpBe (8.6±3.1 to 11.3±3.4%) compared to WT TnpB (3.7±0.7 to 3.8±1.8%) (Extended Data Fig. 3c and Supplementary Table 3). However, indel frequencies in the systemic leaves remained very low (<0.7%) for WT and TnpB variants (Extended Data Fig. 3d and Supplementary Table 4). These findings suggest that the TnpB variants are more active in infiltrated leaves but do not induce editing in systemic leaves when TnpB and reRNA are delivered via two separate TRV2 vectors.

Since the reRNA is endogenously encoded within TnpB’s mRNA at the 3’ end^17^, we decided to encode TnpB and the reRNA as a single transcript unit expressed under pPEBV in TRV2 with reRNA4^PDS^ and ^HH^reRNA4^PDS^ (Fig. 1b and Extended Data Fig. 4a). Strikingly, 88% of plants (14/16) infiltrated with Agrobacterium carrying the eTnpBc variant showed a significant photobleaching phenotype in the systemic leaves compared to no photobleaching in WT TnpB infiltrated plants (Fig. 1c and Extended Data Fig. 4b, c), indicating high-efficiency systemic somatic editing at the *PDS* locus by eTnpBc. In 3 out of 14 plants showing phenotype, we observed complete photobleaching of the terminal stem, including leaves and flowers (Fig. 1d). Although we observed a photobleaching phenotype with the eTnpBe variant (Extended Data Fig. 4d, e), this phenotype was significantly milder compared to that of the eTnpBc variant (Fig. 1c and Extended Data Fig. 4c). The photobleaching phenotype observed with eTnpBc indicates tetra-allelic somatic editing since two homeolog genes encode *PDS* in *N. benthamiana* (Extended Data Fig. 2a).

To determine whether the difference in systemic somatic editing phenotype results from variations in editing efficiency, we assessed editing efficiency in TRV-infiltrated leaves. Interestingly, both eTnpBc and eTnpBe variants with reRNA4^PDS^ and ^HH^reRNA4^PDS^ showed higher editing efficiencies of 47±3% and 50±2% and 43±8% and 45±4%, respectively, compared to WT, which had 10±4% and 13±8 % (Fig. 1e; Supplementary Table 5). These results indicate that although both eTnpBc and eTnpBe induce high somatic editing efficiencies in infiltrated leaves, only eTnpBc causes a significant systemic photobleaching phenotype. We next evaluated systemic editing efficiency and found that eTnpBc induced 71±9% for reRNA4^PDS^ and 62±17% for ^HH^reRNA4^PDS^, which are much higher than those induced by eTnpBe and WT TnpB, which induce 26±4% and 20±4% and 3±1% and 12±4%, respectively (Fig. 1f; Supplementary Table 6). Additionally, we observed that fusing eTnpBc to duplicated SV40nls (Extended Data Fig. 5a) caused a significant systemic photobleaching phenotype (Extended Data Fig. 5b) and increased editing efficiency in both infiltrated (Extended Data Fig. 5c and Supplementary Table 7) and systemic leaves (Extended Data Fig. 5d and Supplementary Table 8) compared to WT TnpB. Overall, these results indicate that eTnpBc fused to FDnls or SV40nls with reRNA4 targeting *PDS* can efficiently edit both *PDS* genes, leading to a strong systemic photobleaching phenotype.

As our high-activity variants were not explicitly codon-optimized for plants, we next generated a series of TRV2 constructs with WT TnpB and variants optimized for either *N. benthamiana* (Nb) or *N. tabacum* (Nt) (Extended Data Fig. 6a and Extended Data Fig. 7a). Both Nb and Nt versions of eTnpBc with reRNA4^PDS^ caused a very mild photobleaching phenotype in the systemic leaves of some plants (Extended Data Fig. 6a, b, and Extended Data Fig. 7a). Additionally, the Nb version of WT TnpB and the eTnpBe variant did not induce a photobleaching phenotype in systemic leaves (Extended Data Fig. 6a, b, and Extended Data Fig. 7a). Assessment of editing efficiency in infiltrated leaves showed 55±8% and 56±6% editing for NbCoeTnpBc with reRNA4^PDS^ and ^HH^reRNA4^PDS,^ respectively (Extended Data Fig. 6c and Supplementary Table 9). Meanwhile, editing efficiencies for the Nb versions of eTnpBe and WT TnpB with reRNA4^PDS^ and ^HH^reRNA4^PDS^ are 29±18%, 20±8%, 7±8%, and 7±8%, respectively (Extended Data Fig. 6c and Supplementary Table 9). The NtCoeTnpBc showed editing efficiencies of 7±2% and 15±3% with reRNA4^PDS^ and ^HH^reRNA4^PDS^ (Extended Data Fig. 7b and Supplementary Table 10). In contrast, the editing efficiency of the NT version of eTnpBc with reRNA2^PDS^ and reRNA3^PDS^ was very low, ranging from 0% to 3% (Extended Data Fig. 7b and Supplementary Table 10). Despite the NbCoeTnpBc showing significant editing in the infiltrated leaves, it failed to induce a strong systemic photobleaching phenotype (Extended Data Fig. 6b). Overall, these results demonstrate that our original eTnpBc sequence^25^, rather than one optimized for the plant host, is more effective at inducing a photobleaching phenotype in both infiltrated and systemic leaves.

Because eTnpBc combined with both designs of reRNA4^PDS^ is highly effective for inducing somatic editing at the *PDS* locus, we tested its activity at additional sites by targeting two *ChlH* homeologs, *ChlH1*-Nbe07g00760 and *ChlH2*-Nbe08g21240^30^ (Extended Data Fig. 2b). These encode the H subunit of magnesium chelatase, an enzyme involved in chlorophyll biosynthesis^32^. Plants infiltrated with reRNA^ChlH^ showed a typical yellow phenotype, indicating very high systemic somatic editing (Fig. 1g). Amplicon sequencing revealed an indel frequency of 90±12% in the systemic leaves with the phenotype (Fig. 1h and Supplementary Table 11). Since two homeologs encode *ChlH* in *N. benthamiana*, the yellow phenotype suggests tetra-allelic somatic editing. These results, along with tetra-allelic *PDS* editing data, demonstrate that eTnpBc can achieve nearly quantitative (i.e. 100%) gene editing in plants.

Since WT ISYmu1 TnpB has recently been shown to cause low-efficiency somatic and heritable editing when delivered via TRV in Arabidopsis^23^, we tested whether ISYmu1 with reRNA4^PDS^ induces a photobleaching phenotype in systemic leaves of *N. benthamiana*. WT ISYmu1 codon optimized for *Arabidopsis thaliana* as reported in^23^ and *N. tabacum*, and co-optimized for human and yeast^25^ fail to produce the *PDS* photobleaching phenotype (Extended Data Fig. 8a). Sequencing of the amplicons from infiltrated leaves shows an editing efficiency of 33±2% and 24±3% for Nt and Hs/Sc versions, respectively (Extended Data Fig. 8b and Supplementary Table 12). We previously demonstrated that ISYmu1 variants (H4Y/V305R and L167G/V305R) informed by the ISDra2 deep mutational scanning dataset showed increased editing activity in yeast cells^25^. Consequently, we expressed these variants along with reRNA4^PDS^ using TRV2. Significant photobleaching was observed in the systemic leaves of all plants infiltrated with the ISYmu1^H4Y/V305R^ and ISYmu1^L167G/V305R^ variants (Extended Data Fig. 8c, d). These results indicate that, as in yeast, ISYmu1 variants are highly active in *N. benthamiana* when delivered via TRV, outperforming WT ISYmu1. Further studies on these variants could shed light on their potential for plant genome editing.

Given the high somatic editing activity of eTnpB variants, we next examined whether heritable, germline editing occurred. Seeds were collected from pods showing photobleaching or from the top third of parent plants infiltrated with either WT TnpB or eTnpB variants. Most seedlings from photobleached pods were completely white (89±9% for reRNA4^PDS^ and 89±8% for ^HH^reRNA4^PDS^) (Fig. 2b, Extended Data Fig. 9b, and complete statistics in Supplementary Table 13). Additionally, 52±14% for reRNA4^PDS^ and 53±20% for ^HH^reRNA4^PDS^ seedlings appeared white when seeds from pods collected from the top third (including a mix of non-photobleached and photobleached pods) of the parent plants infected with eTnpBc were planted (Extended Data Fig. 9b and Supplementary Table 13). None of the progeny seedlings from seeds of parent plants infiltrated with WT TnpB (Fig. 2a, Extended Data Fig. 9a, and Supplementary Table 13) or the eTnpBe variant (Extended Data Fig. 9c, d, and Supplementary Table 13) appeared white.

**Figure 2.**
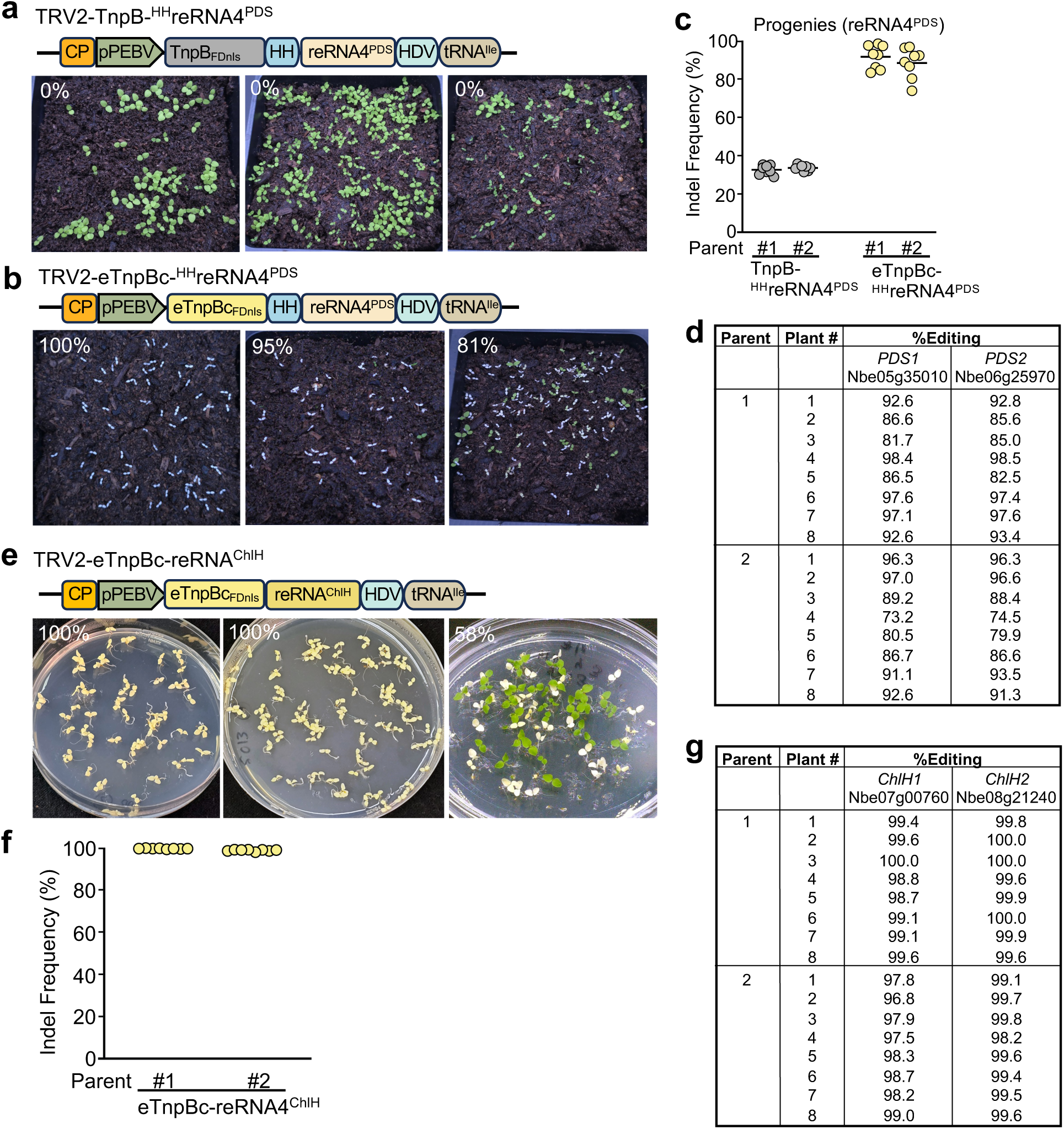
Efficient heritable editing of *PDS* and *ChlH* using the eTnpBc variant. **a-b,** The phenotype of M1 progeny from seeds collected from pods with a photobleaching phenotype or from the top third of plants infected with TnpB (**a**) or eTnpBc (**b**) using ^HH^reRNA4^PDS^. Progeny from three parents are shown. White seedlings indicate tetra-allelic editing and loss of *PDS* function. The inset in the top-left corner of the photo shows the percentage of plants exhibiting the white seedling phenotype. **c,** Indel frequency of eight selected green seedlings from a (see Supplementary Table 11) and white seedlings from b (see Supplementary Table 10) across two parent plants. **d,** Editing frequency of two *PDS* genes, *PDS1* and *PDS2,* of white seedlings from c (see Supplementary Table xx). **e,** M1 progeny phenotype from seeds collected from parent plants infected with TRV expressing eTnpBc-reRNA^ChlH^. Yellow seedlings indicate tetra-allelic editing and loss of *ChlH* function. The insert in the top-left corner of the photo shows the percentage of plants displaying the phenotype. **f,** Indel frequency of eight yellow seedlings from two parents in d (see Supplementary Table x). **g,** Editing frequency of two *ChlH* genes, ChlH1 and *ChlH2*, of yellow seedlings form f (see Supplementary Table xx).

Since *N. benthamiana* is an allotetraploid, the white seedling phenotype indicates editing in both *PDS* homeologs. Assessing the overall editing efficiency of 6-8 randomly selected white seedlings from two representative parent plants infected with eTnpBc showed 90±9% and 96±2% editing frequencies for reRNA4^PDS^ and 88±8% and 92±6% for ^HH^reRNA4^PDS^ (Fig. 2c, Extended Data Fig. 9f, and Supplementary Table 14). Furthermore, the evaluation of editing efficiency for each *PDS* homeolog revealed that both *PDS* genes, *PDS1*-Nbe05g35010 and *PDS2*-Nbe06g25970, exhibited high editing frequencies (Fig. 2d and Supplementary Table 15). In contrast, the editing frequency in green seedlings from parent plants infected with TnpB or eTnpBe ranged from 2±1% to 34±1% (Fig. 2c, Extended Data Fig. 9e, and Supplementary Table 16) and from 16±7% to 24±14% (Extended Data Fig. 9g, h, and Supplementary Table 17), respectively. We also observed heritable *PDS* phenotype and editing with eTnpBc fused to SV40nls (Extended Data Fig. 10 and Supplementary Table 18). These results demonstrate that delivering the eTnpBc variant with reRNA4^PDS^ via TRV achieves highly efficient, tetra-allelic, heritable editing in *N. benthamiana* plants.

We then evaluated heritable germline editing for *ChlH.* We found that 100% of the seedlings from seeds collected from pods showing a yellow phenotype appeared completely yellow (Fig. 2e). Additionally, 58% of the seedlings from pooled seeds collected from the top third of the plant with the *ChlH* phenotype appeared yellow (Fig. 2e). Overall indel frequency in these yellow seedlings ranged from 99-100% indicating very high editing efficiency at *ChlH* locus (Fig. 2f and Supplementary Table 19). Furthermore, editing efficiency at each *ChlH* homeolog ranged from 98-100% (Fig. 2g and Supplementary Table 20). These results indicate that TnpBc also induces very high tetra-allelic, heritable editing at the *ChlH* locus.

Our findings demonstrated that eTnpBc, co-optimized for both human and yeast, fused to FDnls or 2xSV40nls, and combined with reRNA containing the HDV ribozyme at the 3’ end, achieves high editing efficiency. To explore whether the human-yeast co-optimized sequence or variant sequences might be better suited for viral delivery, we performed qPCR to measure viral and TnpB transcript levels. We observed no significant difference across constructs in either infiltrated or systemic leaves (Supplementary Fig. 1). These results imply that differences in editing efficiency could be due to variations in protein expression levels or stability. Additionally, the lack of phenotype in some constructs could result from the loss of parts or the entire insert from the viral vector as the virus spreads systemically through the upper parts of the plant.

Since TRV-expressing eTnpBc leads to high heritable edits in the *PDS* and *ChlH* genes, we tested whether progeny seedlings carry the virus. No virus was detected in the progenies of seedlings edited at the *PDS* or *ChlH* locus (Supplementary Fig. 2). These findings are consistent with previously published results indicating that TRV is not transmitted through the germline to the next generation ^3,6,23^.

To evaluate off-target activity in progeny seedlings displaying the *PDS* and *ChlH* phenotypes, we identified seven genomic off-target sites for reRNA4^PDS^ and six off-target sites for reRNA^ChlH^ using Cas-OFFinder^33^ (Supplementary Figs. 3-5). DNA from five randomly selected seedlings exhibiting the *PDS* photobleaching phenotype and five with the *ChlH* yellow phenotype was used to PCR amplify regions covering the off-target sites (Supplementary Figs. 4-5). Sanger sequence analysis of these amplicons did not reveal any indels at the *PDS* (Supplementary Table 21) or *ChlH* (Supplementary Table 22) off-target sites. These findings indicate that eTnpBc maintains on-target specificity in *N. benthamiana*.

Transformation and regeneration remain major bottlenecks in plant biotechnology. Here, we demonstrate that TRV-mediated delivery of engineered small genome editing effectors results in extremely high, heritable editing efficiencies at targeted loci. These efficiencies in both somatic and heritable editing are comparable to previously reported results using TRV-delivered gRNA in plants already expressing Cas9^3^, and they significantly exceed the efficiencies observed with WT TnpB and AsCas12f proteins^23,24^. Unlike *PDS* and *ChlH*, most genes do not show an obvious visual phenotype when edited. Thus, the high yields observed here—over 50% in pods collected from the top third of the parental plants—point to a future genome editing approach that does not require lengthy transformation, regeneration, or genotyping. In such cases, seeds can be phenotyped directly, greatly accelerating plant breeding. TRV’s broad host range includes key crops such as tomato, pepper, and potato; the principles and findings here should aid in developing transgene-free viral systems for these plants. Lastly, precise edits within alleles of biotechnological interest—such as those achieved through base or prime editing—are likely needed instead of indels. Improving viral delivery systems and engineering more advanced editing effectors will be crucial to generate the necessary edits and germplasm libraries for future precision plant breeding.

## Methods

### Generation of TRV vectors with TnpBs and reRNAs

The vectors TRV1 (pYL192; Addgene #148968) and TRV2-pPEBV-MCS-tRNA^Ileu^ (SPDK3888, Addgene #149276) are described in^3,27^. All DNA fragments used for vector construction in this study were synthesized by Twist Biosciences. reRNA sequences are listed in Supplementary Table 23. Primers used to generate constructs are listed in Supplementary Table 24. Various TRV2 derivatives produced and used in this study are listed in Supplementary Table 25 (see details below). All plasmids created were confirmed by sequencing the insert region.

The following scheme was used to generate TRV2 with TnpB and TnpB variants optimized for a balance of human and yeast codons, with reRNA4^PDS^ as a single-unit vector. To create 4927 (TRV2-TnpB-reRNA4^PDS^), the synthesized TnpB_FDnls_ fragment and reRNA-reRNA4^PDS^-HDV fragment were PCR amplified using primers 104639+104640, and 103925+104230, respectively, followed by digestion with XbaI-KpnI and KpnI-SacI, then cloned into the XbaI-SacI digested SPDK3888. To produce 4928 (TRV2-TnpB-^HH^reRNA4^PDS^), the PCR-amplified HH-reRNA-reRNA4^PDS^-HDV fragment, using primers 104200+104230, was digested with KpnI-SacI and ligated into the XbaI-SacI digested SPDK3888, along with the TnpB_FDnls_ fragment from above. To generate 4784 (TRV2-eTnpBc-reRNA4^PDS^), the amplified eTnpBc_FDnls_ fragment with primers 104365+104366 was digested with XbaI-KpnI, combined with the reRNA-reRNA4^PDS^-HDV fragment from above, then ligated into the XbaI-SacI digested SPDK3888. Vector 4787 (TRV2-eTnpBc-^HH^reRNA4^PDS^) was created by ligating the eTnpBc_FDnls_ fragment and the HH-reRNA-reRNA4^PDS^-HDV fragment into SPDK3888, which was cut with XbaI-SacI.

To generate TRV2 with *Nicotiana benthamiana* codon-optimized TnpB (NbCoTnpB) fused to the duplicated SV40 NLS (4654, TRV2-NbCoTnpB_2xSV40nls_), the synthesized NbCoTnpB_2xSV40nls_ fragment was amplified using 103919+103920 primers, digested with XbaI-NruI, and ligated into SPDK3888 digested with XbaI-Eco35KI. To construct TRV2 with different reRNAs targeting *PDS*, 4655 (TRV2-reRNA1^PDS^), 4657 (TRV2-reRNA2^PDS^), 4742 (TRV2-reRNA3^PDS^), and 4743 (TRV2-reRNA4^PDS^), the respective synthesized reRNA fragments were amplified using 103921+103922, 103921+103923, 103921+104172, and 103921+104173 primer pairs, then digested with XbaI-SacI, and ligated into SPDK3888 digested with the same enzymes.

TnpB and TnpB variants fused to FDnls and reRNA4^PDS,^ as two separate TRV2 vectors, were constructed following this scheme. Fragments for 4984 (TRV2-TnpB_FDnls_), 4949 (TRV2-eTnpBc_FDnls_), and 4985 (TRV2-eTnpBe_FDnls_) were amplified using primer pairs 104639+104640, 104534+104640, and 104639+104640, then digested with XbaI and KpnI, and ligated into SPDK3888 cut with the same enzymes. To generate 4950 (TRV2-reRNA4^PDS^) and 4951 (TRV2-HHreRNA4^PDS^), fragments amplified with primers SP104452+104230 and SP104205+104230 were digested with XbaI and SacI and then ligated into SPDK3888 cut with those enzymes.

The following scheme was used to generate NbCoTnpB and TnpB variants with FDnls and reRNA4 flanked by HDV at the 3’ end or flanked by HH at the 5’ end and HDV at the 3’ end as a single transcript in the TRV2 vector. To produce 4947 (TRV2-NbCoTnpB-reRNA4^PDS^) and 4948 (TRV2-NbCoTnpB-^HH^reRNA4^PDS^), NbCoTnpB was amplified using a synthesized fragment as a template with primer pairs 104729+104730 (fragment I). reRNA4PDS-HDV (fragment II) and HH-reRNA4PDS-HDV (fragment III) were amplified from plasmids 4784 and 4787, respectively, using primer pairs 104717+101572. Fragments I and II, and fragments I and III, were cloned into SPDK3888, cut with XbaI and SacI, using NEBuilder HiFi DNA Assembly. To generate 4981 (TRV2-NbCoeTnpBc-reRNA4^PDS^) and 4982 (TRV2-NbCoeTnpBc-^HH^reRNA4^PDS^), NbCoeTnpBc was PCR amplified with a synthesized fragment using primer pairs 104718+104640, digested with XbaI and KpnI, and ligated into plasmids 4784 and 4787, respectively, cut with the same enzymes. To create 4945 (TRV2-NbCoeTnpBe-reRNA4^PDS^) and 4946 (TRV2-NbCoeTnpBe-^HH^reRNA4^PDS^), NbCoeTnpBe was amplified using primers 104729+104730 (fragment I). reRNA4^PDS^ was amplified with primers 104717+101572 (fragment II), and ^HH^reRNA4^PDS^ with the same primers (fragment III). Fragments I and II, and fragments I and III, were cloned into SPDK3888, cut with XbaI and SacI, using NEBuilder HiFi DNA Assembly.

To generate a *Nicotiana tabacum* codon-optimized eTnpBc (NbCoeTnpBc) variant fused to FDnls and different PDS guide RNAs as a single TRV2 vector, the following method was employed. NbCoeTnpBc was amplified using primers 104367+104368 and digested with XbaI and KpnI. The fragments reRNA2^PDS^-HDV, reRNA3^PDS^-HDV, and reRNA4^PDS^-HDV were amplified with primers 104230+103925 and digested with KpnI and SacI. The HH-reRNA2PDS-HDV, HH-reRNA3^PDS^-HDV, and HH-reRNA4^PDS^-HDV fragments were amplified with primers 104230+104200 and digested with KpnI and SacI. Fragments NbCoeTnpBc and reRNA2^PDS^-HDV, reRNA3^PDS^-HDV, or reRNA4^PDS^-HDV were ligated into SPDK3888, digested with XbaI and SacI, resulting in vectors 4788 (TRV2-NtCoeTnpBc-reRNA2^PDS^), 4789 (TRV2-NtCoeTnpBc-reRNA3^PDS^), and 4790 (TRV2-NtCoeTnpBc-reRNA4^PDS^). Fragments NbCoeTnpBc and HH-reRNA2^PDS^-HDV, HH-reRNA3^PDS^-HDV, or HH-reRNA4^PDS^-HDV were ligated into SPDK3888, digested with XbaI and SacI, producing vectors 4791 (TRV2-NtCoeTnpBc-^HH^reRNA2^PDS^), 4792 (TRV2-NtCoeTnpBc-^HH^reRNA3^PDS^), and 4793 (TRV2-NtCoeTnpBc-^HH^reRNA4^PDS^).

TnpB constructs with SV40nls were assembled as described. TnpB and eTnpBc were amplified from synthesized fragments using primers 104639+104641 and 104534+104641, respectively, then digested with XbaI and KpnI. The reRNA4^PDS^-HDV and HH-reRNA4^PDS^-HDV fragments were amplified with primers 103925+104230 and 104200+104230, respectively, and digested with KpnI and SacI. The resulting fragments were ligated into SPDK3888, cut with XbaI and SacI, resulting in vectors 4931 (TRV2-TnpB_2xSV40nls_-reRNA4^PDS^), 4932 (TRV2-TnpB_2xSV40nls_-^HH^reRNA4^PDS^), 4929 (TRV2-eTnpBc_2xSV40nls_-reRNA4^PDS^), and 4930 (TRV2-eTnpBc_2xSV40nls_-^HH^reRNA4^PDS^).

TRV2 with ISYmu1 constructs were created by amplifying the SV40nls-NtCoISYmu1-NLS fragment and the HsCoISYmu1-2xSV40nls using primers 104180+104178 and 104930+104931, respectively, then digesting with XbaI and KpnI. The reRNA4^PDS^-HDV fragment was amplified and digested with KpnI and SacI. The two fragments were ligated into 4916, digested with XbaI and SacI, resulting in vectors 4756 (TRV2-NtcoISYmu1-reRNA4^PDS^) and 5034 (TRV2-HscoISYmu1-reRNA4^PDS^). To clone AtCoISYmu1, as described in^23^, the PCR product was amplified using primers 105196+ 100703 with Addgene #236574 as template, digested with XbaI and KpnI, and cloned into 5034 cut with the same enzyme, resulting in vector 5166 (TRV2-AtCoISYmu1-reRNA4^PDS^). To clone ISYmu1 variants H4Y/V305R and L167G/V305R, primers 104930+104931 and SP104932+104931 with plasmid template described in^25^ were used to amplify PCR product, digested with XbaI and KpnI, and cloned into 4756 vector cut with the same enzymes resulting in vectors 5035 (TRV2-ISYmu1^L167G/V305R^-reRNA4^PDS^) and 5036 (TRV2-ISYmu1^H4Y/V305R^-reRNA4^PDS^).

### Plant Growth Conditions and Agrobacterium-Mediated TRV Delivery into Plants

*Nicotiana benthamiana* plants were grown in a growth chamber set at 26°C with a 12-hour light and 12-hour dark cycle, a light intensity of 100 μE m-2 sec-1, and 50% humidity.

TRV1 and various TRV2 derivative vectors listed in Supplementary Table 15 were transformed into Agrobacterium tumefaciens strain GV2260 competent cells and grown on lysogeny broth (LB) agar medium supplemented with KSRC (kanamycin at 50 μg/ml, streptomycin at 50 μg/ml, rifampicin at 25 μg/ml, and carbenicillin at 50 μg/ml) for three days at 28°C. Two colonies per construct were picked and streaked onto a fresh KSRC LB agar plate, then grown overnight at 28°C and confirmed by colony PCR. Colonies were inoculated into 2 mL of LB liquid culture with KSRC, grown in a shaker incubator overnight at 28°C, made into glycerol stocks, and stored at - 80°C. To conduct each experiment, Agrobacterium containing TRV1 and the necessary TRV2 derivative constructs were streaked on LB agar plates with KSRC and grown overnight at 28°C. About 5-6 mL of LB liquid media with KSRC was inoculated with each TRV2 construct and incubated in a shaker at 28°C for 14-16 hours. For each TRV2 derivative construct to be infiltrated, 6 mL of TRV1 was inoculated. Agrobacterium cells were collected by centrifugation at 3500×g for 20 minutes, resuspended in agroinfiltration medium containing 10 mM MgCl_2_, 10 mM 2-(N-morpholine)-ethanesulfonic acid (MES), and 250 μM 3′,5′-dimethoxy-4′-hydroxyacetophenone (acetosyringone) in mQ H2O) to a OD_600_ of 1.0, and incubated with gentle shaking for at least 3 hours at room temperature. TRV1 and TRV2 derivatives were mixed at a 1:1 ratio and infiltrated onto three diagonally opposite leaves of 2.5-3-week-old *N. benthamiana* plants using a 1 mL needleless syringe. Infiltrated plants were either kept on the growth light carts in the laboratory or in a growth chamber set at 24°C with a 12-hour light/12-hour dark photoperiod, a light intensity of 100 μE m-2 sec-1, and 50% humidity. Somatic editing phenotypes were monitored throughout the plant’s lifecycle. Generally, initial signs of somatic editing were visible two weeks after TRV infiltration.

### Assessment of somatic editing efficiency in TRV-infiltrated leaves and upper systemic leaves

To evaluate editing frequency in the TRV-infiltrated leaves, four days post-infiltration, leaf samples approximately 8-10 mm in diameter were collected from the infiltrated areas. These samples were placed in 35 μl of dilution buffer provided with the PHIRE Plant Direct PCR Kit (Thermo Fisher Scientific) in an Eppendorf tube and crushed with a 1 ml pipette tip by pressing it against the tube walls. The tubes were centrifuged for 5 minutes at 14,000 rpm in an Eppendorf centrifuge. To amplify the amplicon surrounding the target region, 2 μl of leaf lysate was used in a PCR reaction with barcoded primers. The PCR conditions included an initial denaturation at 98°C for 5 min, 39 cycles of 98°C for 10 s, 55°C for *PDS*, and 60°C for *ChlH* for 10 s, followed by 72°C for 20 s, with a final incubation at 72°C for 5 min. The barcoded primers used for amplicon generation are listed in Supplementary Table 16. The amplicons were purified using the Zymo PCR Clean-up kit and eluted in 25 μl of water. The concentration of the purified amplicons was measured with a Qubit 4 Fluorometer (Thermo Fisher Scientific). Amplicons were pooled at equal molar ratios and sent for Amplicon-EZ next-generation sequencing (NGS) (Genewiz/Azenta). Sequencing files received from Genewiz/Azenta were demultiplexed, and the resulting paired-end .fastq files were used to determine indels and their frequencies using CRISPResso2^34^, following the parameters described in^17^, which ignore substitutions and insertions to prevent false positives.

To evaluate editing frequency in the systemic leaves, leaf samples approximately 8-10 mm in diameter were collected from the upper systemic leaves at 3-5 weeks after infiltration of TRV and processed as described to assess the editing frequency in the infiltrated leaves.

### Assessment of heritable germline editing efficiency and mutation rates

To evaluate heritable editing efficiency, pods displaying a photobleaching phenotype were collected as they formed. Additionally, pods from the top 3^rd^ of the plants were harvested approximately three months after TRV infiltration. About 50-300 seeds were planted per pot in soil, and the trays with pots were covered with plastic domes before being placed in a growth chamber set at 24°C day/night conditions with a 12-hour light/12-hour dark photoperiod and a light intensity of 100 μE m-2 sec-1, with 50% humidity. After about 10-14 days, the phenotypes (green and white) were recorded to estimate the percentage of progeny seedlings exhibiting the white photobleaching phenotype. To determine mutation frequencies, approximately 6-8 green and white seedlings were individually collected and processed to generate amplicons flanking the target region as described above using barcoded primers (Supplementary Table 26). The amplicons were then sent for Amplicon-EZ NGS (Azenta/Genewiz). Demultiplexed paired-end .fastq files were used to identify indel types and their frequencies with CRISPResso2^34^, following the parameters described in^17^, which ignore substitutions and insertions to avoid false positives.

### Quantitative real-time PCR analysis

Leaf samples were collected from infiltrated tissue 5 days after infiltration with various TRV constructs, and systemic tissue was collected 3 weeks post-infiltration. Total RNA was extracted with TRIzol reagent (Thermo Fisher Scientific) and treated with RNase-free DNase I (New England Biolabs). First-strand cDNA from TRV, HsCoTnpBs, NbCoTnpBs, and PP2A was synthesized using SuperScript III reverse transcriptase (Thermo Fisher Scientific) with primers 105432, 105444, 105449, and 105425, respectively. Real-time qPCR was conducted on a Bio-Rad CFX Opus 96 with iTaq Universal SYBR Green Supermix (Bio-Rad), using 2.5 mM of primers targeting the TRV CP region (105433+105434), HsCoTnpBs (105443+105444), NbCoTnpBs (105448+105449), and PP2A (105426+105425), with 5% diluted cDNA. Expression levels were calculated via the ΔΔcT method and normalized to PP2A.

### RT-PCR

Total RNA was extracted from progeny seedlings with the *PDS* photobleaching phenotype and seedlings with the ChlH yellow phenotype using TRIzol reagent (Thermo Fisher Scientific). Following extraction, the RNA was treated with RNase-free DNase I (New England Biolabs). First-strand cDNA synthesis for TRV and PP2A was performed using SuperScript III reverse transcriptase (Life Technologies) with primers 105432, which target the TRV2 CP region, and 105425 for PP2A. PCR was carried out with 2 μl of cDNA in a 50 μl reaction containing 20 pmol of TRV-specific primers (105433+105434) and PP2A-specific primers (105426+105425), along with Taq polymerase. The PCR protocol involved an initial denaturation at 95 °C for 2 minutes, followed by 25 cycles of 95 °C for 15 seconds, 55 °C for 15 seconds, and 72 °C for 30 seconds, ending with a final extension at 72 °C for 5 minutes. About 5 μL of the PCR products was then analyzed by 1% agarose gel electrophoresis.

### Off-target analysis

To assess the target specificity of editing by the eTnpBc variant, CRISPR RGEN Tools (Cas-OFFinder, http://www.rgenome.net/cas-offinder/) was used to identify potential genomic off-target sites with the “TTGAT” TAM and 4-6 mismatches in the target sequences related to reRNA4^PDS^ and reRNA^ChlH^ ^33^. Amplicon sequences around the PAM and off-target sites were amplified from five seedlings with a white *PDS* phenotype and five with a yellow *ChlH* phenotype, using the PHIRE Plant Direct PCR Kit (Thermo Fisher Scientific), as previously described, with primers listed in Supplementary Table 27. The amplicons were purified with the Zymo PCR Clean-up Kit, eluted in 25 μL of water, and sequenced via Genewiz/Azenta Sanger sequencing service.

### Reporting summary

Further information on research design is available in the Nature Research Reporting Summary linked to the paper.

## Data availability

All data are available in the paper or the supplementary material. Additionally, all amplicon-EZ sequencing files will be accessible on the NCBI Sequence Read Archive at the time of publication.

## Acknowledgments

We thank Jinqiao Hu, Chloe Jang, and Trevan Murakami from the University of California, Davis, for assistance with some genotyping. We also thank Luke Oltrogge and Cynthia Terrace (University of California, Berkeley, CA) for their help with data analysis and sequencing. Additionally, we appreciate Jung-Un Park from the University of California, Berkeley, for critically reviewing the manuscript. This research was supported by the National Science Foundation (NSF) grant IOS-2303522 (to SPD-K), NSF grant DGE 2146752 (to BWT and RFW), and the Innovative Genomics Institute (IGI; to SPD-K and DFS). Any opinions, findings, and conclusions or recommendations expressed in this material are those of the author(s) and do not necessarily reflect the views of the NSF. DFS is an Investigator at the Howard Hughes Medical Institute (HHMI).

## Author Contributions

UN, JER, DFS, and SPD-K conceptualized the project. SPD-K and DFS supervised the study. UN designed and conducted the experiments with assistance from TN and JER. UN analyzed the plant data, and JER analyzed the amplicon sequencing data and prepared the graphs. RFW and BWT provided materials before publication and participated in discussing the results. UN, JER, DFS, and SPD-K prepared the figures and tables. UN, JER, DFS, and SPD-K wrote the manuscript with input from all authors. All authors reviewed and approved the final version.

## Competing Interests

A patent application has been filed by The Regents of the University of California, with UN, JER, BWT, RFW, DFS, and SPD-K listed as inventors. DFS is a co-founder and member of the scientific advisory board at Scribe Therapeutics.

