## Supplementary File for "High-efficiency, transgene-free plant genome editing by viral delivery of an engineered TnpB"

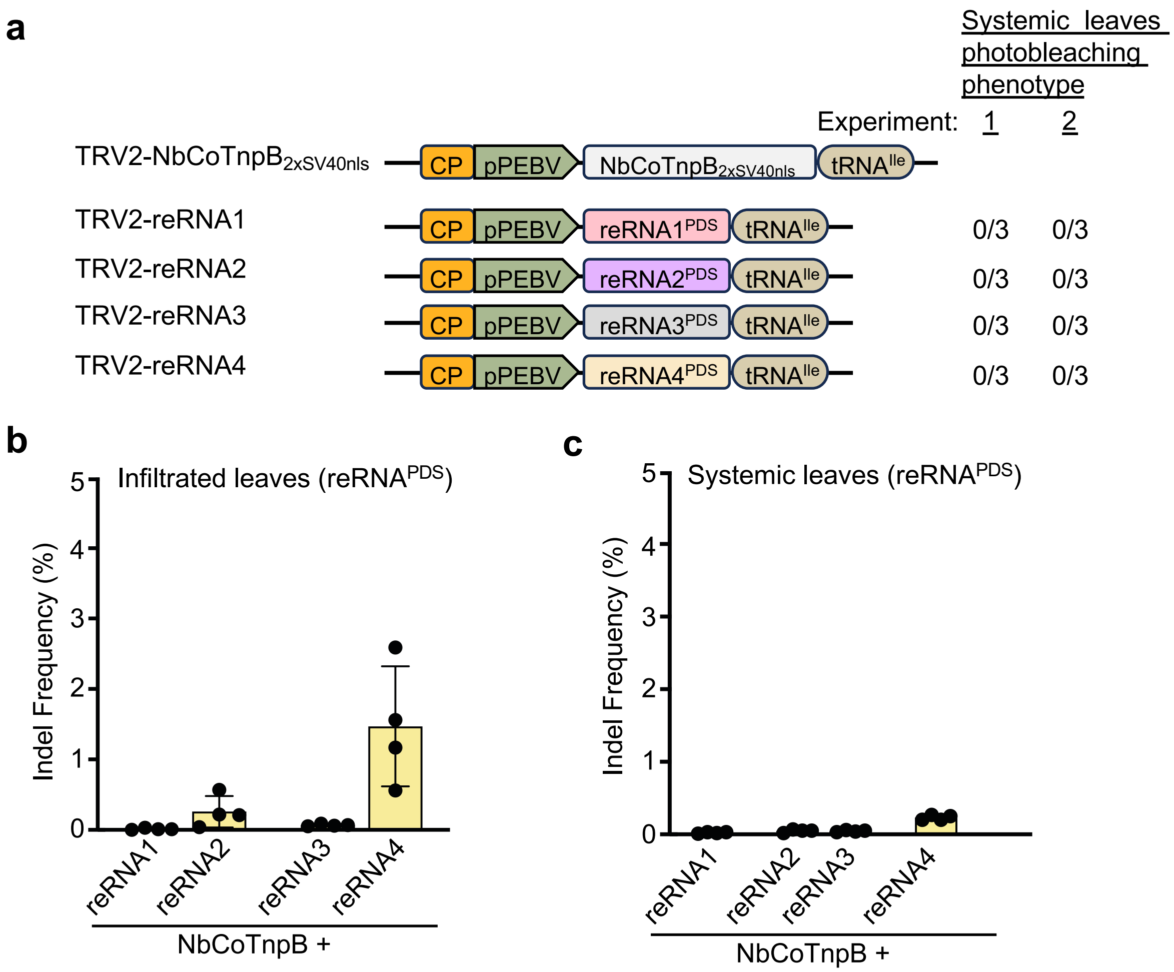

**Extended Data Figure 1. Delivery of TnpB and reRNA from two separate TRV2 vectors does not induce the somatic *PDS* photobleaching phenotype. a.** Agrobacterium-mediated infiltration of TRV2 carrying *Nicotiana benthamiana* codon-optimized TnpB fused with the nuclear localization sequence from SV40 (NbCoTnpB$_{2xSV40nls}$) and various reRNAs (reRNA1-4) targeting the *PDS* genes did not produce a photobleaching phenotype in the systemic leaves. n=3 plants; two independent experiments. **b-c,** Indel frequency at the *PDS* locus in infiltrated (b) and systemic leaves (c). Data are plotted as the mean and standard deviation (SD) from biological replicates (n=4); two tissue samples from each plant.

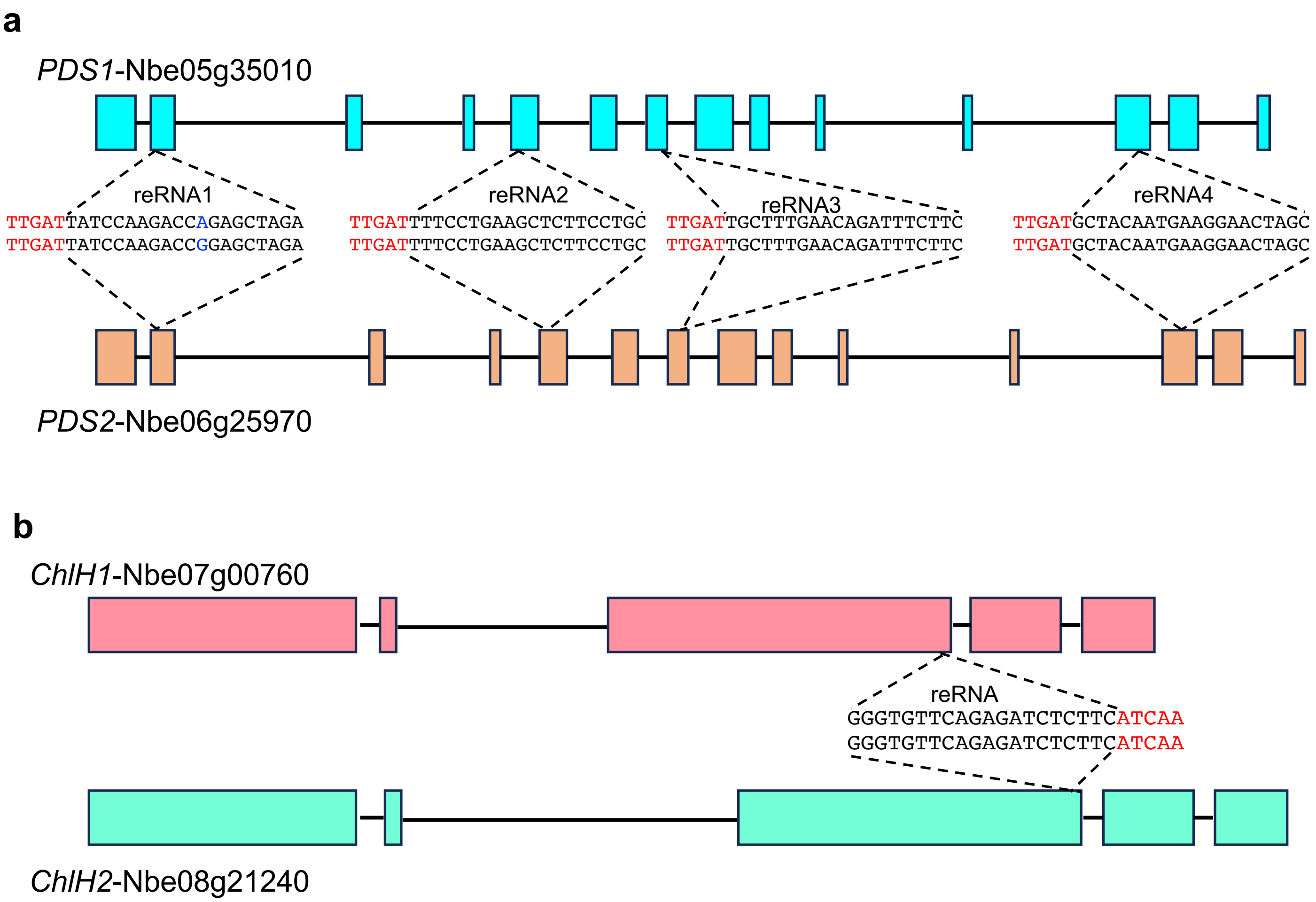

**Extended Data Figure 2. Diagram of *N. benthamiana PDS* (a) and *ChlH* (b) homeologs.**
Exons are shown as colored boxes, while introns are represented as lines. Four reRNA sequences targeting *PDS* (**a**) and one reRNA targeting *ChlH* (**b**) are indicated. Red letters in the reRNA sequence indicate the transposon-associated motif (TAM).

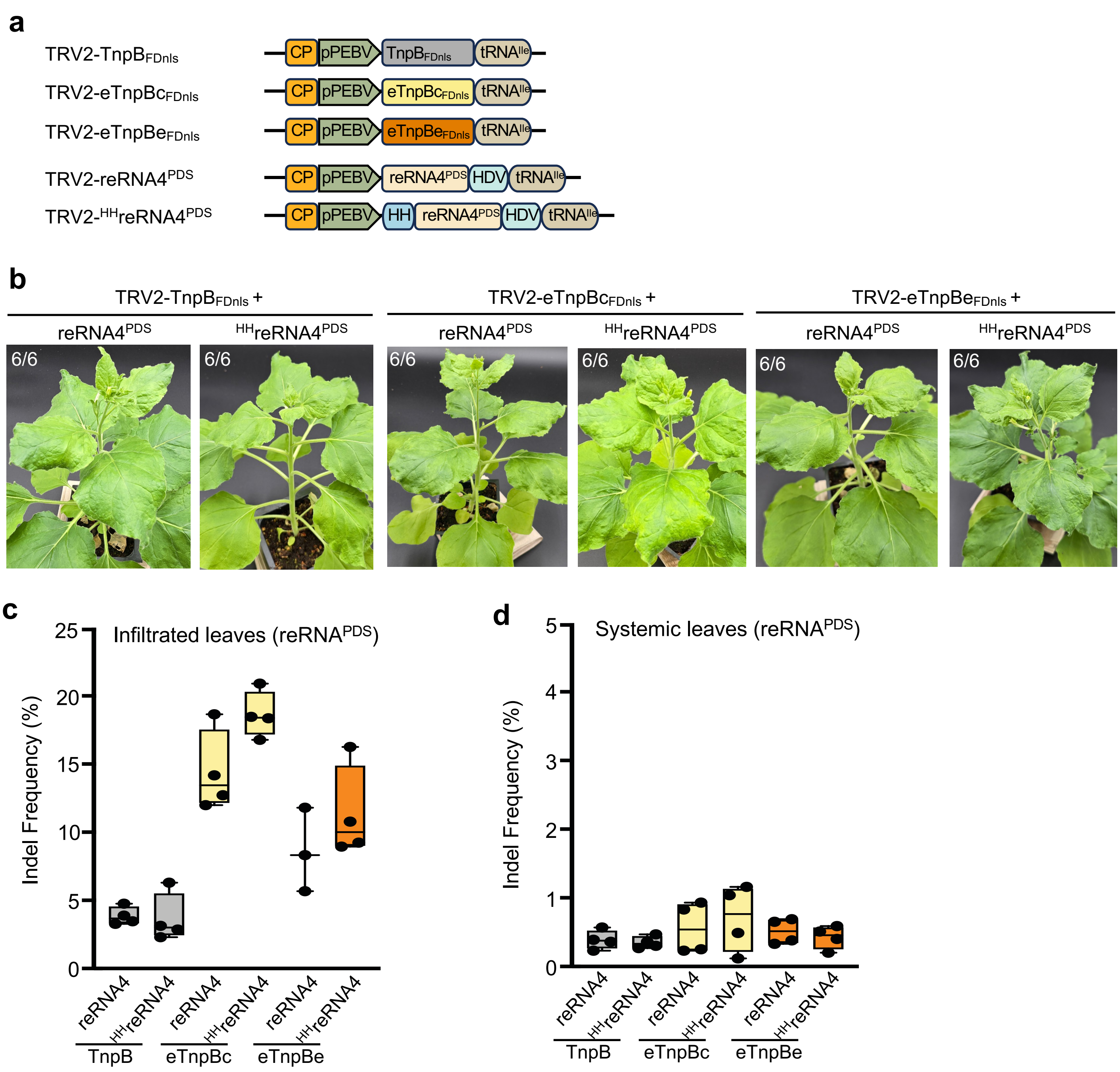

**Extended Data Figure 3. Delivery of eTnpB and reRNA from two separate TRV2 vectors does not induce the somatic *PDS* photobleaching phenotype. a,** Diagram showing TRV2 vectors with wild-type TnpB and two variants, eTnpBc and eTnpBe, fused to the nuclear localization sequence from Arabidopsis FD protein, along with two reRNA4 designs—one without a self-cleaving hammerhead (HH) ribozyme at the 5' end of reRNA and one with HH ribozyme. Both designs include a self-cleaving hepatitis delta virus (HDV) ribozyme at the 3' end of the reRNA. **b,** Plants infiltrated with TRV2 containing TnpB, eTnpBc, and eTnpBe, with two reRNA4 designs, were photographed approximately 5 weeks after TRV infiltration. The inset in the top-left corner of the photo indicates the number of plants exhibiting the corresponding phenotype. n=3 plants; 2 independent experiments. **c-d,** Indel frequency at the *PDS* locus in infiltrated (**b**) and systemic leaves (**c**). Data is from biological replicates (n=4). The box shows the interquartile range (Q1 to Q3), the line inside indicates the median, and the whiskers extend to the most extreme points within 1.5× IQR of the quartiles.

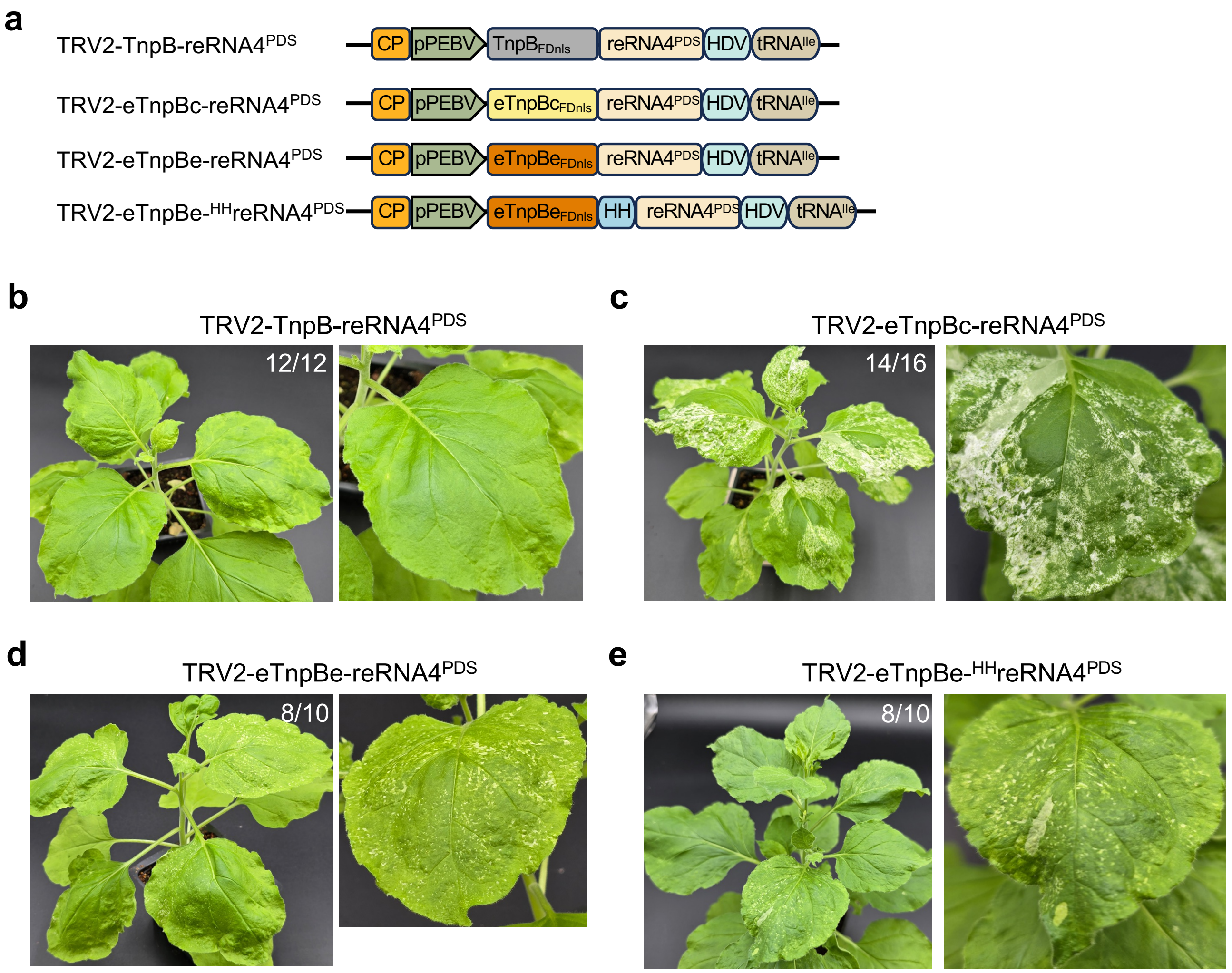

**Extended Data Figure 4. Virus delivery of the enhanced variant eTnpBc results in an efficient systemic somatic editing phenotype, but this is not observed with the eTnpBe variant.**
**a,** Diagram showing TRV2 vectors with TnpB, eTnpBc, and eTnpBe fused to FDnls with reRNA4 targeting *PDS* genes, featuring an HDV self-cleaving ribozyme at the 3' end of the reRNA (reRNA4$^{PDS}$). Additionally, a diagram with eTnpBe$_{FDnls}$, which has a self-cleaving HH ribozyme at the 5' end and an HDV ribozyme at the 3' end of the reRNA ($^{HH}$reRNA4$^{PDS}$), is shown. **b-e,** Phenotypes of plants infected with TRV expressing TnpB with reRNA4$^{PDS}$ (**b**), eTnpBc with reRNA4$^{PDS}$ (**c**), eTnpBe with reRNA4$^{PDS}$ (**d**), and eTnpBe with $^{HH}$reRNA4$^{PDS}$ (**e**). Plants were photographed 2.5 weeks after TRV infiltration. The white sectors on the leaves, which appear bleached, indicate the loss of *PDS* function. In **b-e**, the inset in the top right corner of the photo indicates the number of plants showing the corresponding phenotype. n≥3 plants; 4 independent experiments for b and c. n≥2 plants; 4 independent experiments for d and e.

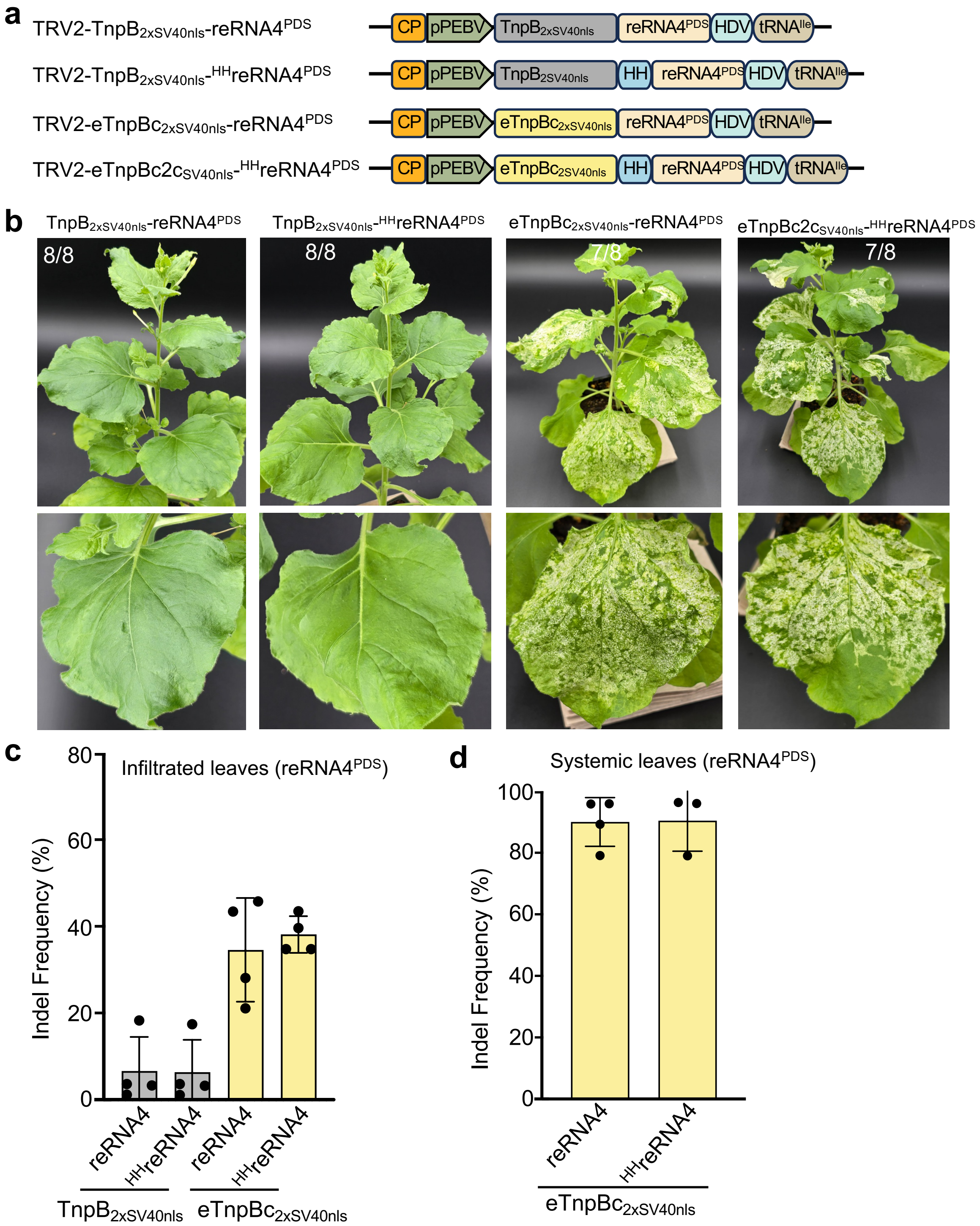

**Extended Data Figure 5. Virus delivery of eTnpBc fused to SV40nls enables efficient systemic somatic editing. a,** Diagram showing TRV2 vectors with TnpB and eTnpBc fused to duplicated SV40 nuclear localization sequence (2xSV40nls) and two designs of reRNA4: one with HDV at the 3' end of reRNA ($reRNA4^{PDS}$) and the other with HH at the 5' and HDV at the 3' end of reRNA ($^{HH}reRNA4^{PDS}$). **b,** Phenotypes of plants infiltrated with TRV expressing TnpB and eTnpBc fused to SV40nls with two guides targeting *PDS* genes. Plants were photographed 3.5 weeks after TRV infiltration. The white, bleached sectors on the leaves indicate loss of *PDS* function. The inset in the top left corner of the photo indicates the number of plants showing the corresponding phenotype. n=2 plants; 4 independent experiments. **c-d,** Indel frequency at the *PDS* locus in both infiltrated (**c**) and systemic (**d**) leaves. Data are shown as the mean and SD from biological replicates (n=4).

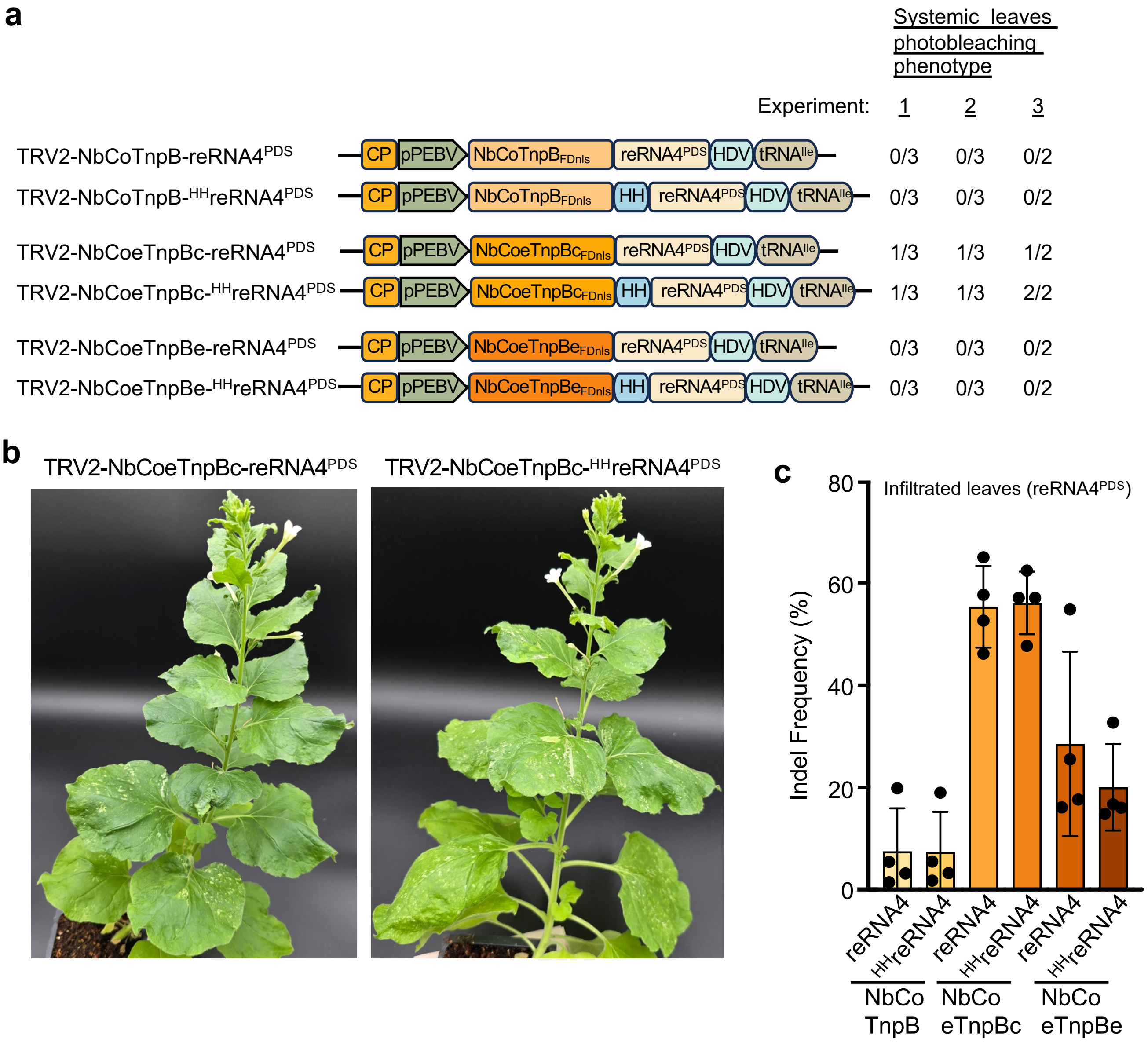

**Extended Data Figure 6. Virus delivery of NbCoeTnpBc causes a mild somatic *PDS* photobleaching phenotype in some plants. a,** Number of plants showing a somatic photobleaching phenotype after infiltration with TRV2 carrying *Nicotiana benthamiana* codon-optimized TnpB (NbCoTnpB), NbCoeTnpBc, or NbCoeTnpBe, along with two reRNA4 designs targeting the *PDS* genes, reRNA4$^{PDS}$ and $^{HH}$reRNA4$^{PDS}$. n≥2 plants; three independent experiments. **b,** Plants infiltrated with NbCoeTnpBc and two reRNA4 designs from the experiment in a displaying a mild photobleaching phenotype, were photographed five weeks after TRV infiltration. **c,** Indel frequency in the infiltrated leaves at the *PDS* locus for two reRNA4 designs with the NbCo version of TnpB and TnpB variants. Data are presented as the mean and SD from biological replicates (n=3).

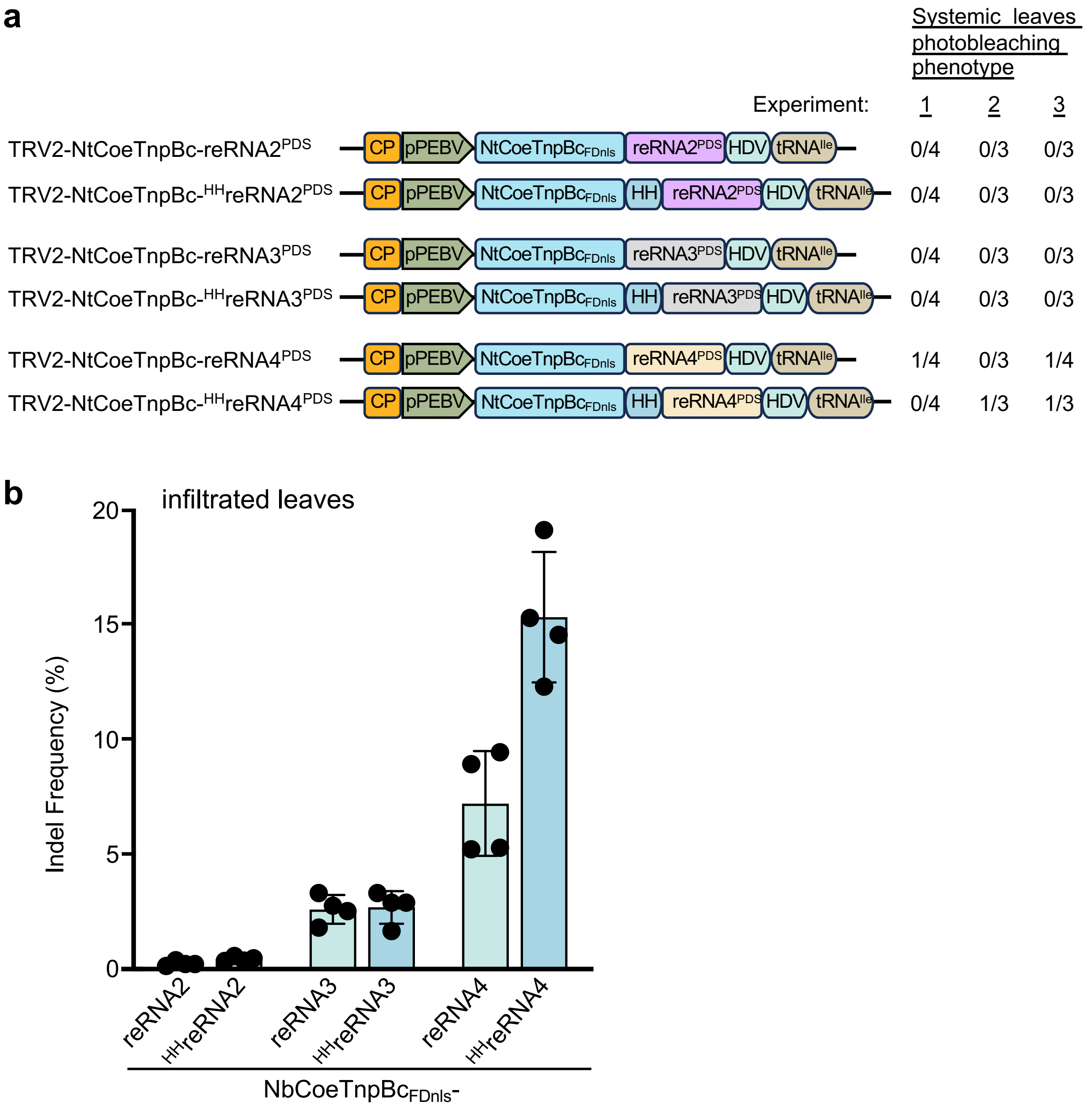

**Extended Data Figure 7. Virus delivery of NtCoeTnpBc causes a mild somatic photobleaching phenotype in some plants. a,** Number of plants showing a somatic photobleaching phenotype after infiltration with TRV2 carrying *Nicotiana tabacum* codon-optimized eTnpBc variant (NbCoeTnpBc) along with two designs of reRNA2, reRNA3, and reRNA4 targeting *PDS* genes. n≥3 plants; three independent experiments. **b,** Indel frequency at the *PDS* locus in the infiltrated leaves for the indicated reRNAs with NbCoeTnpBc. Data represent the mean and SD from biological replicates (n=3).

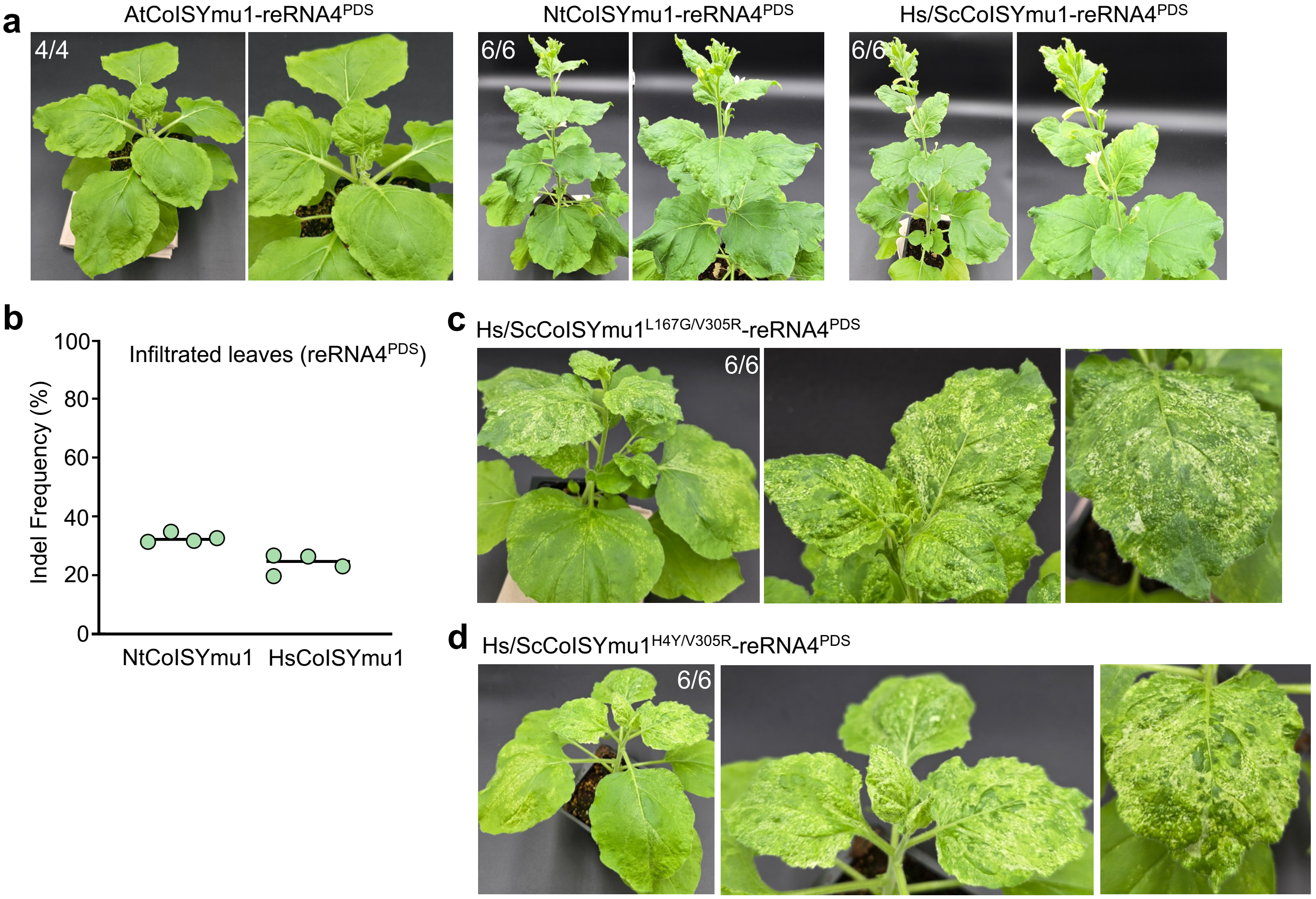

**Extended Data Figure 8. Virus delivery of ISYmu1 TnpB mutants causes systemic somatic editing.** **a,** Phenotype of plants infiltrated with TRV expressing Arabidopsis codon-optimized ISYmu1 (AtCoISYmu1), *N. tabacum* codon-optimized ISYmu1 (NtCoISYmu1), and human/yeast codon-optimized ISYmu1 (Hs/ScCoISYmu1) with reRNA4 targeting *PDS*, including an HDV ribozyme at the 3' end of the reRNA. n=2 or 3 plants; experiments repeated twice. **b,** Indel frequency at the *PDS* locus in plants infiltrated with NtCoISYmu1 and Hs/ScCoISYmu1 (see Supplementary Table 7). **c-d,** Phenotype of plants infiltrated with TRV expressing Hs/ScCoISYmu1$^{L167G/V305R}$ (**c**) and Hs/ScCoISYmu1$^{H4Y/V305R}$ (**d**) mutants with reRNA4 targeting *PDS*. Plants were photographed approximately 2.5 weeks after infiltration (left panel). The white sectors on the leaves, which appear bleached, indicate the loss of *PDS* function. n=3 plants; experiments repeated twice. The inset in the top-right corner of the photo indicates the number of plants exhibiting the corresponding phenotype.

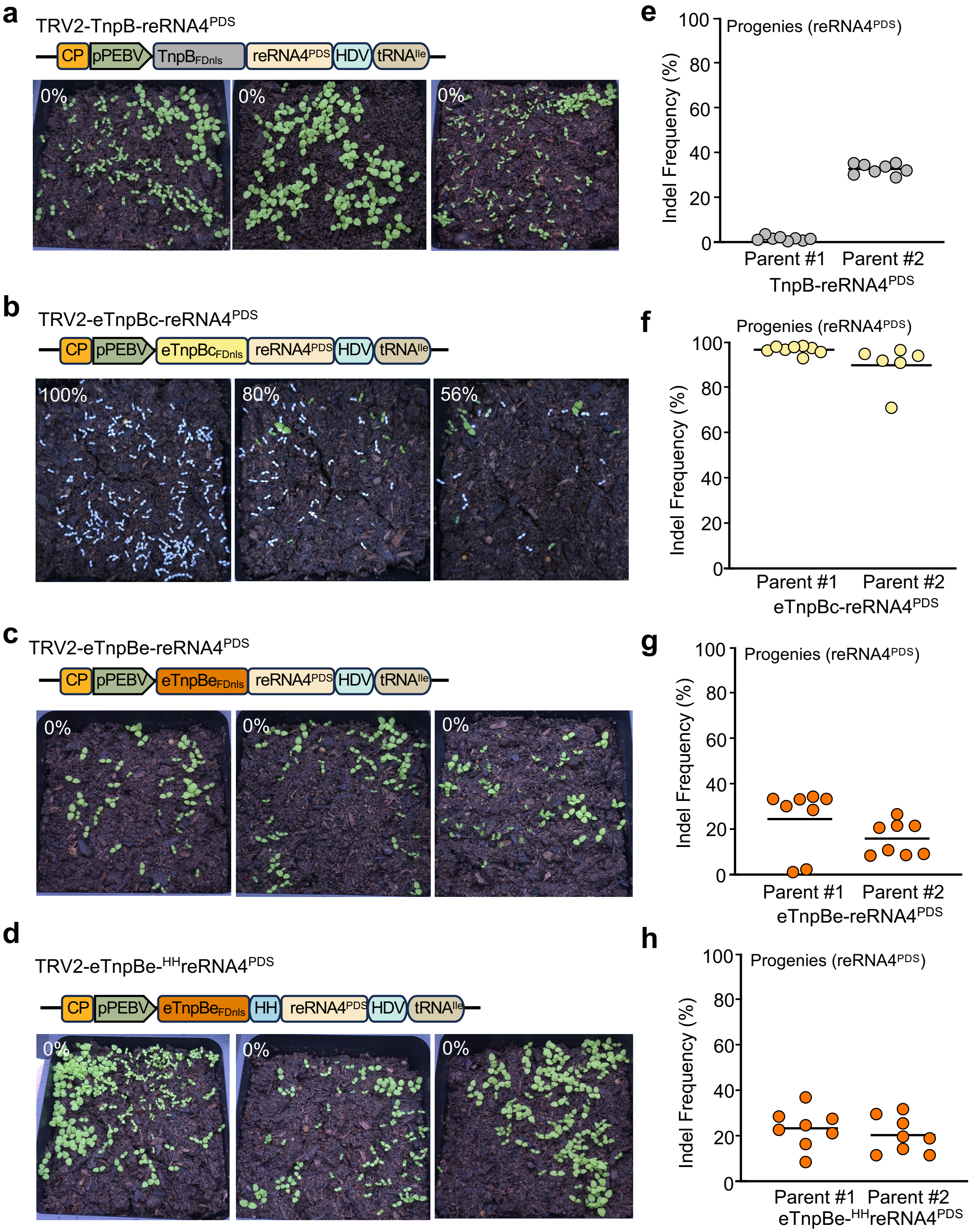

**Extended Data Figure 9. Efficient, heritable editing of *PDS* using the eTnpBc variant, but not with wild-type TnpB or the eTnpBe variant. a-d,** The phenotype of M1 progenies from seeds collected from pods showing a photobleaching phenotype or from the top third of infected plants, with TnpB-reRNA4$^{PDS}$ (**a**), eTnpBc-reRNA4$^{PDS}$ (**b**), eTnpBe-reRNA4$^{PDS}$ (**c**), and eTnpBe-$^{HH}$reRNA4$^{PDS}$ (**d**). Progenies from three parent plants are shown. White seedlings indicate tetra-allelic editing and loss of *PDS* function. The inset in the top left corner of the photo shows the percentage of plants displaying the white seedling phenotype. **e-h,** Indel frequency at the *PDS* locus in 6-8 selected green seedlings from a (**e**) (see Supplementary Table 11), c (**g**) and d (**h**) (see Supplementary Table 12), and white seedlings from b (**f**) (see Supplementary Table 10) across two parent plants.

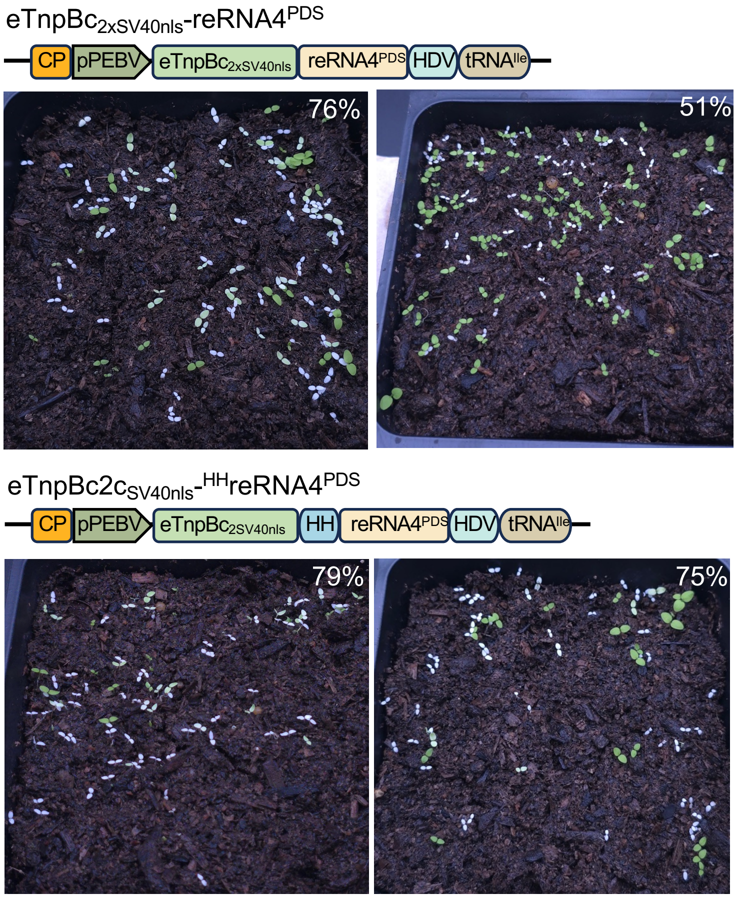

**Extended Data Figure 10. Efficient heritable editing of *PDS* using the eTnpBc variant fused to SV40nls.** The phenotype of M1 progenies from seeds collected from the top third of plants infiltrated with eTnpBc$_{2xSV40nls,}$ along with reRNA4$^{PDS}$ (top panels) and $^{HH}$reRNA4$^{PDS}$ (bottom panels). Progenies from two parent plants are shown. White seedlings indicate tetra-allelic editing and the loss of *PDS* function. The inset in the top right corner of the photo shows the percentage of plants displaying the white seedling phenotype.

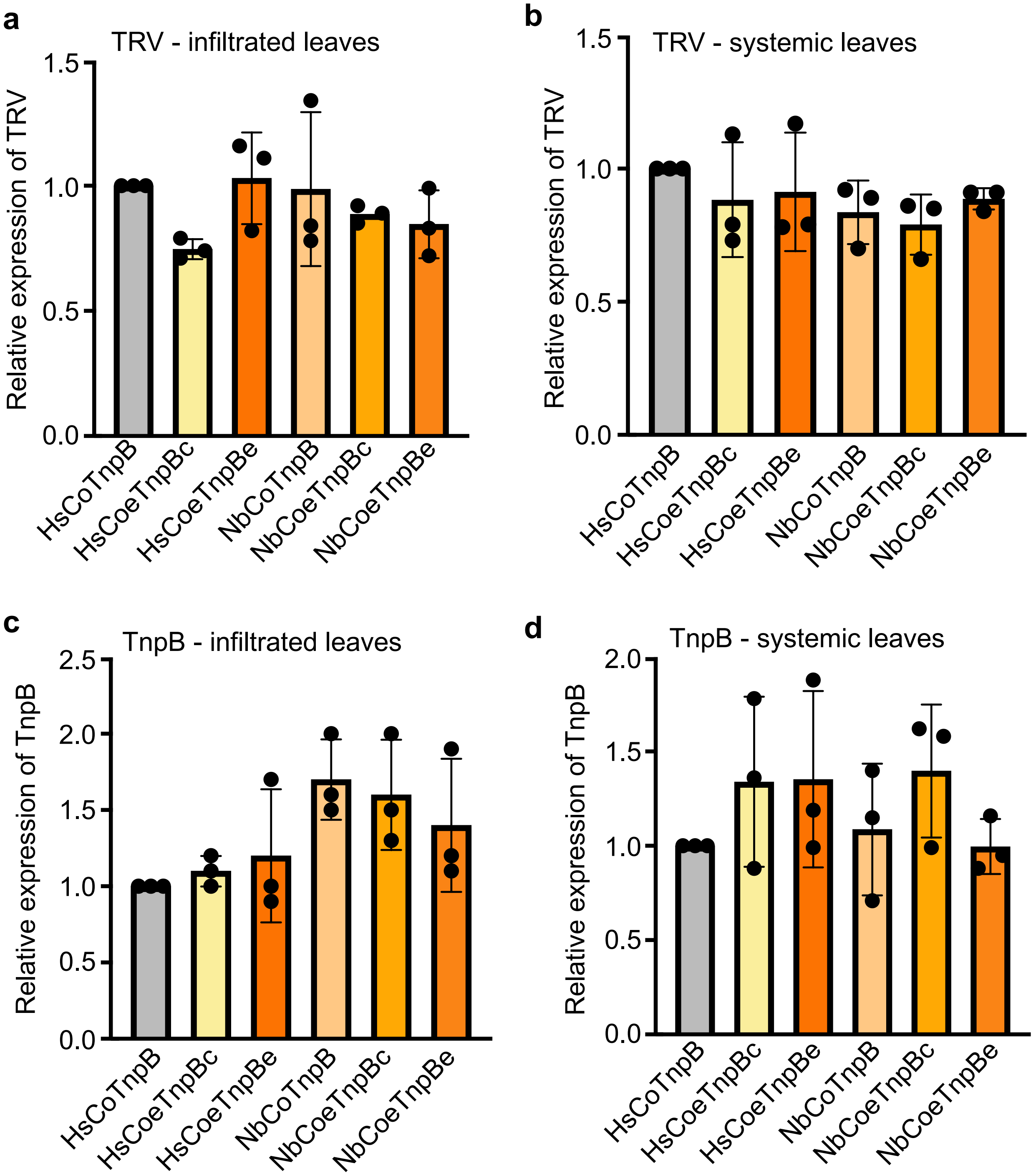

**Supplementary Figure. 1. Transcript expression of TRV, TnpB, and TnpB variants.** **a-d,** cDNA from infiltrated (**a, c**) and systemic (**b, d**) leaf tissues with indicated TnpB and TnpB variants was analyzed by qPCR using virus-specific (**a, b**) or TnpB-specific (**c, d**) primers. Values are shown as relative expression compared to HsCoTnpB, with means ± SD (n=3). Data were analyzed using the ΔΔCt method and normalized to PP2A. No significant differences were detected by ANOVA with post-hoc Tukey HSD test, $\alpha = 0.05$.

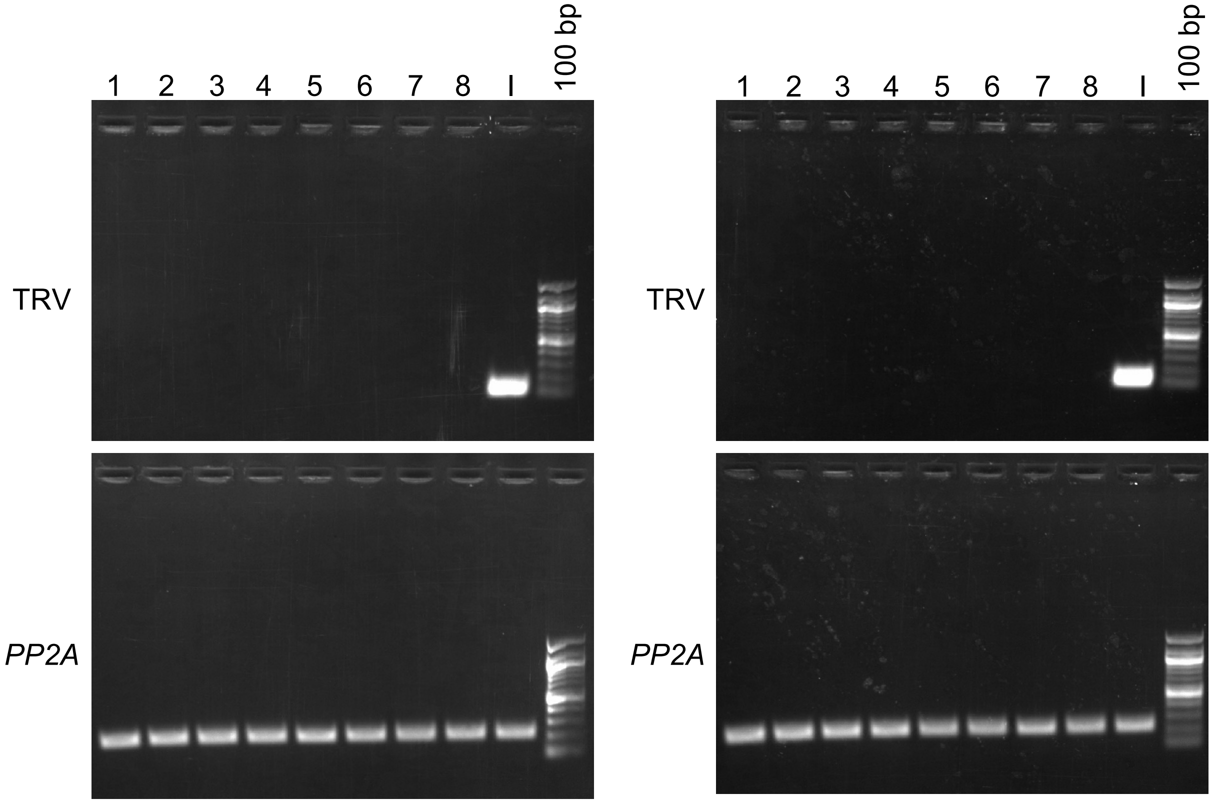

**Supplementary Figure 2. Edited progenies are virus-free.** Tissue from eight (1-8) *PDS* edited white seedlings (left) or *ChlH* edited yellow seedlings (right) and TRV infected plant (I) were used for RT-PCR using primers that amplify the coat protein region of TRV (top panels) and the endogenous PP2A gene (bottom panels). 100 bp, NEB 100 base pair ladder.

| | |
|---|---|
| PDS4 | TTGATGCTACAATGAAGGAACTAGC |
| PDS4.1off | TTGATGCTTCAATAAAGGAAATGTC |
| PDS4.2off | TTGATGCAACTGTAAAGGCACTAGC |
| PDS4.3off | TTGATGCTACACTGATGAAACTTCC |
| PDS4.4off | TTGATGCTAGTATGAGAGAACCAGC |
| PDS4.5off | TTGATGCTGCAATTAAAGAAGTAAC |
| PDS4.6off | TTGATGCTACTCTGTAGGAATTATC |
| PDS4.7off | TTGATGTTGCAATGACGAAACTAAC |

| | |
|---|---|
| ChlH | TTGATGAAGAGATCTCTGAACACCC |
| ChlH.1off | TTGATATTGAGATCTTTGATCACCC |
| ChlH.2off | TTGATGAAGACAGATCTGAAAACCA |
| ChlH.3off | TTGATTAAAAGAGATCTGAAAACCC |
| ChlH.4off | TTGATGAAGAGATCTCTGTATATTT |
| ChlH.5off | TTGATGAAGAGATTTCTGTTTAGCC |
| ChlH.6off | TTGATGAAGAGTTCGCAAAACTGCC |

**Supplementary Figure 3. Analyzed off-target sites.** Off-target sites corresponding to reRNA4$^{PDS}$ (top) and reRNA$^{ChlH}$ (bottom) were identified using Cas-OFFinder. Underlined letters indicate the TAM sequence, and red letters indicate mismatches relative to the target site used.

**PDS4.1off**

CAGGCAATGTAGACCTGTCATCAGTTGCTTGATCAACCAGAATAGAGTTCTCTAAACTGTACTGCTGGTCGA
TCAACATATGTTCCTGCCTTTTTGTATTGTGCTGAGCTTTCACAGCAGACAACTTTGACATTTCCTTTATTG
AAGCATCAAAATGAATTGGTATTGTTGACTCCTCTACTGGTTCAGCAGATACAAAAGTTGGATCTCTATTAA
CATTGAAAGAAACAGAATCCTCTGTACCATTTCTTTGATCAGTTAATGTACTGTTTTGATTCTTACAATTAT
GTACCAACAACTGCTGGTC

**PDS4.2off**

GAGGATAATTGAGTTCCTGAAGGGCCAATCGGGTTGATCATGGAATGTGATAACCCAAAGAAACCAATATTT
ACTTTCACTACGATCGTTGTATAGATTCAGCCCGCTGGGGATGCTGAGGCAAAGATGTCTATGGCACTTGAG
TTTGATGCAACTGTAAAGGCACTAGCGCCAATTGAGGCCGAGTTCGTGTCTCCAGCAAATGCGCCTGCACCA
CTTGAAGTTACAATTTTACCCTCAAAGCAGATGCTC

**PDS4.3off**

GACACTTTCCCTTTACCTCTACTTTGAAGACTTCTTCCTCCTGCCTTTGCAGGCTAGTATTTACTCTAAAGA
TCTGTACAAGGGATTGCATCCCCTTGATGCTACACTGATGAAACTTCCAAATTGTGTCCAATCAATTAACTG
TGTCTCAAACTCCAAACAAATTAGGTTCGGAAAAATCCTATCCCCAAGCTTTCATATTTAGCGGGTGTACTG
TTTTTCCTAAAACGTGTTGATTGGACCATAAGGGGATTTCTACCAAGGATTTGGTTAGGGAATAAGATTACA
TTTGCGGTGCT

**PDS4.4off**

GGACCAAACTCTTGTTGGTAGTTGAATTCCTACATGGAGCGGCTTTGTTGTACATCATGAATCTTTCTGTTT
ACACACCTCTCCACATCCATCACCAGGATTTCGAAAGTAATGAAACCTTACTATTGGGTTAACCTTAAACAT
CAGTTGATGCTAGTATGAGAGAACCAGCAGAGAGGGGGCGAGCAATCTTTTCCATCAGGGTTGCCACAGTAA
AATATACCGGAGAGGGGCAGCTCCTAACAAAAGCACCATGAGACGGGGCTTTTCTCCCAAACATCTCTTTCT
GCACGAGTGGCTATCACGCCTAACCTTTGAAATTG

**PDS4.5off**

GCAGTTCTATCGGACAAGAGAAAGAGATCACTATATGACACAGAGTTGCTTCACCTCATGGGAGATGGATGA
TATATGAGTTTCTTTCTTCTTCTGTCTTTCCCATTTCAGCTGTAACTTATGCTAGCATATGACTAAAATGTG
CTTTTGATGCTGCAATTAAAGAAGTAACTGTATATTATTGTTATTCCCATGTTAATAGGAATTCTATGAGTT
TATGCAAGAAATGATAACAATGATGAAAGCTGAGAAATCTAGGATATTATGTTCCTTCAGTTTCTGTTATTT
ATGAACGTGAACAATATATGTCTTTTTGATGCCAT

**PDS4.6off**

AATAATTTCGCACGCTTGACTCACGAGAAATTGCTAATTGTTGACCATATTCTATACATCAAGCACCAGAGA
TAATTCCTACAGAGTAGCATCAATTGCTTTTGTCAATATTTTCACCAAGAGTAATGAATCTGATTCCACTAT
CACTCTGTTATAACCAATCACCACGCATGTTCACAAAAGTTATTATCTAGATCAGCTCACTGGTATATTCAT
GT

**PDS4.7off**

GGAAGTATTTCAAGAGATGCCAAGTTTTGCTAAGTATTTGAAATATTTGATCACCAAGAAGAATACCATAAA
AAATGAAGTGGTGAATGTTACGAACCGAGTTAGTTTCGTCATTGCAACATCAACCGTTCAAAAGAAAGCTGA
CCCGAGAGAATTTACCATTCCTTGTACCATCGCGGTACATGATTTTACAAGAGCCCTATGTAATAATGGAGC
TAGAATCGACTTGA

**Supplementary Figure 4.** Sequences of potential off-target sites for the reRNA4$^{PDS}$ target. The reRNA sequence region is underlined, and the TAM sequence appears in red.

**ChlH.1off**

GCCATAACAAAGAACATAGCCATAGAATGTGCATTCATCTTGACATCTTCAAGCATATTTTGACACTACTAT
ATCAATTAGTGGTTATAAATCTAACATCATGTCCGACAACGGGTGATCAAAGATCTCAATATCAATTGCAGT
GACAGGATCTCTACGAACGGGATTAATTTTTTTGTTTATAGCTAATGAGAATTGAACTCATGATCTTATGT
AGATTTTGACAACCCTTAATCACTAAATTGCA

**ChlH.2off**

CTCTCCACATAGTAAGGCATTAGGTTTCAATAAAAAAACTATGTGTGTGAAAGAAAATATTCATATTATTTT
TTATGAATCTAACATTTGGCCGAGAAAGGTCTGCAGGATGAAGATTATGACATAGGATTTACTGGTGAGGGA
GATACAAAGTAATTTGATGAAGACAGATCTGAAAACCACACGAAAATTAAAAATGGTCATGAGGAACCAGAA
GAAGAAAGAACTACTCTAATTACTATCAGTATAATGAAGTGACACCTACTAACCTGTTCCTTTGGGTCACTC
CTCGAGTGAATCTTCTCCGGGTATACAGATCAAAC

**ChlH.3off**

CAGCATCTTCAACAAACAAATGTCGGGGGGAAGAAAATTAGTCATTTTAGAAAAATTATAGTTCCTTTTACT
ACTCTATTTCCTTTTACTTTTCTGATTATTCTGTCTATGTTCCTCGTTTGCTCTTTGTAAGTACGAATATTT
CTATATCTTCTGTTGATTCGTATTTACCATAAGTTAGGGTTTTCAGATCTCTTTTAATCAAACCAAAAATGG
ATTCGATTCTCTAATTTCAAGTATTATTATATCCATATTTGTCCTCTATCGTACTACTGCCCGATTTTTCTT
TTATAGTTTTTGTCAATATTTATCTAGTCCTAAAACTAGGATTTTCAGACCTCTTTTTCGAACAAAAATTGG
GTTGATTGTCTGATTTTGAATAATATTTGTGTCTCTATATATCTTGTGGTATTGTG

**ChlH.4off**

CCTGTTTGGTCAGAGGTATATCGAGGATCCTGGGGTTTGAAGTTGCATATTGCTTCAGCAGCTGGTGACCTA
CTTCAGGTCCTCTATTTCTCGATAATTTCTGTTTTTGACACATAGGACTTGCTGATCATTTTTTATCAACTT
GAATTGTAACTTACCAAAATATACAGAGATCTCTTCATCAAAAAAGAAAGAATATTTACAAAGACATAATAA
TTACTGAGACTCGCTGCTTAAGCATCAACAACGAAAAGCATACCCTATGTGACCTTACTTGTCTAGC

**ChlH.5off**

AGGACATCATAAGCAGTTGTCATAAGGTCCTTCGAACCATGGGTATTGTAGAAACCAATGGGGTCGACTTCG
CCTTTTTTCGCCTAGCAGGATCCGCCAAGACTTGATGAAGAGATTTCTGTTTAGCCAGGCCAGTAGGGTCGC
CATCATTAACCTGGGATCAGTTTTTTGAGCTATTTCTGGGGAAGTTTCTCCCAGTTACTCAGAGAGATGTCC
TTCGCAGGCAGTTTGAGCGCCTCCAGCAGGGCTCTATGTCAGTTATGCAGTATGAGACTCGATTTATTGATC
TCGCACGTCAGTA

**ChlH.6off**

GCCAATCCTTCAATGCTTCTATCTTTTGGGGCTCCACTTGAATCCCCCTACCGGAAATAACATGTCCCAGAA
AAATGAAGGACTGCGACCAAAATTCACATTTTGAGAATTAGAATACAACTGGTGTTGATGAAGAGTTCGCAA
AACTGCCCTGAGGTGATCGGCATGGTCCTCCCTGCTTCGCGAATAAACAAGGATGTCGTCAATAAAATTAAC
CATAAAGGAGTCTAGAAATAGCTTGAACCCGATTCATAAGATCCATGAAAGCTTTCTGGGCGTTGATTATCC
CAAAAGATATGACCAAGAATTCGAAGTAACCATAGCAGGTTGTGA

**Supplementary Figure 5.** Sequences of potential off-target sites for the reRNA4$^{ChlH}$ target. The reRNA sequence region is underlined, and the TAM sequence appears in red.

**Supplementary Table 1.** *PDS* editing frequencies in infiltrated tissue of plants infiltrated with NbCoTnpB and reRNAs from two separate TRV2 vectors

| Construct | Plant # | Overall editing frequency | Average editing frequency | Standard deviation |
|---|---|---|---|---|
| NbCoTnpB + reRNA1$^{PDS}$ | 1 | 0.01 | 0.01 | 0.01 |
| | 2 | 0.01 | | |
| | 3 | 0.03 | | |
| | 4 | 0.00 | | |
| NbCoTnpB + reRNA2$^{PDS}$ | 1 | 0.04 | 0.26 | 0.22 |
| | 2 | 0.21 | | |
| | 3 | 0.22 | | |
| | 4 | 0.57 | | |
| NbCoTnpB + reRNA3$^{PDS}$ | 1 | 0.06 | 0.07 | 0.02 |
| | 2 | 0.09 | | |
| | 3 | 0.05 | | |
| | 4 | 0.07 | | |
| NbCoTnpB + reRNA4$^{PDS}$ | 1 | 2.59 | 1.47 | 0.85 |
| | 2 | 1.56 | | |
| | 3 | 1.17 | | |
| | 4 | 0.56 | | |

**Supplementary Table 2.** *PDS* editing frequencies in systemic tissue of plants infiltrated with NbCoTnpB and reRNAs from two separate TRV2 vectors

| Construct | Plant # | Overall editing frequency | Average editing frequency | Standard deviation |
|---|---|---|---|---|
| NbCoTnpB + reRNA1$^{PDS}$ | 1 | 0.01 | 0.02 | 0.01 |
| | 2 | 0.03 | | |
| | 3 | 0.03 | | |
| | 4 | 0.02 | | |
| NbCoTnpB + reRNA2$^{PDS}$ | 1 | 0.02 | 0.05 | 0.02 |
| | 2 | 0.07 | | |
| | 3 | 0.05 | | |
| | 4 | 0.05 | | |
| NbCoTnpB + reRNA3$^{PDS}$ | 1 | 0.04 | 0.05 | 0.01 |
| | 2 | 0.05 | | |
| | 3 | 0.06 | | |
| | 4 | 0.03 | | |
| NbCoTnpB + reRNA4$^{PDS}$ | 1 | 0.27 | 0.23 | 0.04 |
| | 2 | 0.20 | | |
| | 3 | 0.20 | | |
| | 4 | 0.25 | | |

**Supplementary Table 3.** *PDS* editing frequencies in infiltrated tissue of plants infiltrated with TnpB and TnpB variants with two designs of reRNA4$^{PDS}$ from two separate TRV2 vectors

| Constructs | Plant # | Overall editing frequency | Average editing frequency | Standard deviation |
|---|---|---|---|---|
| TnpB + reRNA4$^{PDS}$ | 1 | 4.76 | 3.8 | 0.7 |
| | 2 | 3.28 | | |
| | 3 | 3.89 | | |
| | 4 | 3.46 | | |
| TnpB + $^{HH}$reRNA4$^{PDS}$ | 1 | 3.13 | 3.7 | 1.8 |
| | 2 | 6.31 | | |
| | 3 | 2.87 | | |
| | 4 | 2.31 | | |
| eTnpBc + reRNA4$^{PDS}$ | 1 | 18.65 | 14.4 | 3.0 |
| | 2 | 14.18 | | |
| | 3 | 11.98 | | |
| | 4 | 12.71 | | |
| eTnpBc + $^{HH}$reRNA4$^{PDS}$ | 1 | 16.78 | 18.6 | 1.7 |
| | 2 | 20.91 | | |
| | 3 | 18.46 | | |
| | 4 | 18.36 | | |
| eTnpBe + reRNA4$^{PDS}$ | 1 | 11.8 | 8.6 | 3.1 |
| | 2 | 5.67 | | |
| | 3 | 8.32 | | |
| eTnpBe + $^{HH}$reRNA4$^{PDS}$ | 1 | 10.79 | 11.3 | 3.4 |
| | 2 | 8.94 | | |
| | 3 | 9.24 | | |
| | 4 | 16.25 | | |

**Supplementary Table 4.** *PDS* editing frequencies in systemic tissue of plants infiltrated with TnpB and TnpB variants with two designs of reRNA4$^{PDS}$ from two separate TRV2 vectors

| Constructs | Plant # | Overall editing frequency | Average editing frequency | Standard deviation |
|---|---|---|---|---|
| TnpB + reRNA4$^{PDS}$ | 1 | 0.23 | 0.4 | 0.1 |
| | 2 | 0.57 | | |
| | 3 | 0.39 | | |
| | 4 | 0.36 | | |
| TnpB + $^{HH}$reRNA4$^{PDS}$ | 1 | 0.30 | 0.4 | 0.1 |
| | 2 | 0.47 | | |
| | 3 | 0.36 | | |
| | 4 | 0.27 | | |
| eTnpBc + reRNA4$^{PDS}$ | 1 | 0.23 | 0.6 | 0.4 |
| | 2 | 0.83 | | |
| | 3 | 0.93 | | |
| | 4 | 0.25 | | |
| eTnpBc + $^{HH}$reRNA4$^{PDS}$ | 1 | 0.12 | 0.7 | 0.5 |
| | 2 | 1.16 | | |
| | 3 | 1.04 | | |
| | 4 | 0.49 | | |
| eTnpBe + reRNA4$^{PDS}$ | 1 | 0.69 | 0.6 | 0.2 |
| | 2 | 0.65 | | |
| | 3 | 0.38 | | |
| | | 0.33 | | |
| eTnpBe + $^{HH}$reRNA4$^{PDS}$ | 1 | 0.20 | 0.4 | 0.2 |
| | 2 | 0.59 | | |
| | 3 | 0.40 | | |
| | 4 | 0.51 | | |

**Supplementary Table 5.** Somatic editing efficiency in infiltrated leaves at the *PDS* locus targeted by TnpB and TnpB variants with reRNA4$^{PDS}$

| Construct | Plant # | 4 most common indels | n_deleted | Reads | % Edited reads | Overall editing frequency | Average editing frequency | Standard deviation |
|---|---|---|---|---|---|---|---|---|
| | | **TTGAT**<u>GCTACAATGAAGGAACTAGC</u>GAAGCTTTTCCCT* | Wild-type | | | | | |
| TnpB-reRNA4$^{PDS}$ | 1 | TTGATGCTACAATGAAGGA----GCGAAGCTTTTCCAT | 4.0 | 103 | 8.0 | 7.6 | 10 | 4 |
|  |  | TTGATGCTACAATGAAGGAACTG----AGCTTTTCCCT | 4.0 | 102 | 7.9 |  |  |  |
| | | TTGATGCTACAATGAAGGAACTAGC-AAGCTTTTCCCT | 1.0 | 100 | 7.8 | | | |
|  |  | TTGATGCTACAATGAAGGAA--------GCTTTTCCCT | 8.0 | 80 | 6.2 |  |  |  |
| | 2 | TTGATGCTACAATGAAGGAACTAGC-AAGCTTTTCCCT | 1.0 | 193 | 12.8 | 8.0 | | |
|  |  | TTGATGCTACAATGAAGGAA--------GCT-----CT | 13.0 | 162 | 10.8 |  |  |  |
|  |  | ---------------------------AGCTTTTCCCT | 86.0 | 160 | 10.6 |  |  |  |
|  |  | TTGATGCTACAATGAAGGAAC----GAAGCTTTTCCCT | 4.0 | 137 | 9.1 |  |  |  |
|  | 3 | TTGATGCTACAATGAAGGA----GCGAAGCTTTTCCCT | 4.0 | 139 | 13.3 | 8.7 |  |  |
|  |  | TTGATGCTACAATGAAGGAACAA----AGCTTTTCCCT | 4.0 | 99 | 9.5 |  |  |  |
| | | TTGATGCTACAATGAAGGAACTAGC-AAGCTTTTCCCT | 1.0 | 69 | 6.6 | | | |
|  |  | TTGATGCTACAATGAAGAAA--------GCTTTTCCCT | 8.0 | 60 | 5.7 |  |  |  |
|  | 4 | TTGATGCTACAATGAAGGAAC-------GCTTTTCCCT | 7.0 | 619 | 16.1 | 15.8 |  |  |
|  |  | TTGATGCTACA-----------------GCTTTTCCCT | 17.0 | 592 | 15.4 |  |  |  |
| | | TTGATGCTACAATGAAGGAA---GCAAAGCTTTTCCCT | 3.0 | 585 | 15.3 | | | |
|  |  | TTGATGCTACAATGAAGGAAC----GAAGCTTCTCCCT | 4.0 | 233 | 6.1 |  |  |  |
| TnpB-$^{HH}$reRNA4$^{PDS}$ | 1 | ---------------------------AGCTTTTCCCT | 86.0 | 462 | 19.1 | 7.3 | 13 | 8 |
| | | TTGATGCTACAATGAAGGAACTAGC-AAGCTTTTCCCT | 1.0 | 399 | 16.5 | | | |
|  |  | TTGATGCTACAATGAAGGAA--------GCT-----CT | 13.0 | 226 | 9.4 |  |  |  |
|  |  | TTGATGCTACAATGAAGGAAC----GAAGCTTTTCCCT | 4.0 | 174 | 7.2 |  |  |  |
|  | 2 | TTGATGCTACAATGGAGGAACTA----AGCTTTTCCCT | 4.0 | 103 | 9.7 | 25.3 |  |  |
|  |  | TTGATGCTACAATGAAGGA----GCGAAGCTTTTCCCT | 4.0 | 88 | 8.3 |  |  |  |
| | | TTGATGCTACAATGAAGGAACTAGC-AAGCTTTTCCAT | 1.0 | 81 | 7.6 | | | |
|  |  | TTGATGCTACAATGAAGGAAC----GAAGCTTTTCCCT | 4.0 | 75 | 7.0 |  |  |  |
|  | 3 | ---------------------------AGCTTTTCCCT | 86.0 | 142 | 12.2 | 10.5 |  |  |
| | | TTGATGCTACAATGAAGGAACTAGC-AAGCTTTTCCCT | 1.0 | 112 | 9.6 | | | |
|  |  | TTGATGCTACAATGAAGGAAC----GAAGCTTTTCCCT | 4.0 | 106 | 9.1 |  |  |  |
|  |  | TTGATGCTACAATGAAGGAA--------GCT-----CT | 13.0 | 99 | 8.5 |  |  |  |
|  | 4 | TTGATGCTACAATGAAGGA----GCAAAGCTTTTCCCT | 4.0 | 136 | 11.4 | 8.8 |  |  |
| | | TTGATGCTACAATGAAGGAACTAGC-AAGCTTTTCCCT | 1.0 | 118 | 9.9 | | | |
|  |  | TTGATGCTACAATGAAGGAACTA----AGCTTTTCCCT | 4.0 | 95 | 8.0 |  |  |  |
|  |  | TTGATGCTACAATGAAGGAAC----GAAGCTTTTCCCC | 4.0 | 79 | 6.6 |  |  |  |
| eTnpBc-reRNA4$^{PDS}$ | 1 | TTGATGCTACAATGAAGGA----GCGAAGCTTTTCCCT | 4.0 | 1600 | 17.3 | 49.9 | 47 | 3 |
|  |  | TTGATGCTACAATGAAG------------CTTTTCCCT | 12.0 | 441 | 4.8 |  |  |  |
|  |  | TTGATGCTACAATGAAGCAA-------AGCTTTTCCCT | 7.0 | 381 | 4.1 |  |  |  |
|  |  | TTGATGCTACAATGAAGGAACTG----AGCTTTTCCCT | 4.0 | 227 | 2.5 |  |  |  |
|  | 2 | TTGATGCTACAATGAAGGA----GCGAAGCTTTTCCCT | 4.0 | 1364 | 15.5 | 47.3 |  |  |
|  |  | TTGATGCTACAATGAAG------------CTTTTCCCT | 12.0 | 459 | 5.2 |  |  |  |
|  |  | TTGATGCTACAATGAAGCAA-------AGCTTTTCCCT | 7.0 | 372 | 4.2 |  |  |  |
|  |  | TTGATGCTATAATGAA-----------AGCTTTTCCCT | 11.0 | 272 | 3.1 |  |  |  |
|  | 3 | TTGATGCTACAATGAAGGA----GCGAAGCTTTTCCCT | 4.0 | 1166 | 13.4 | 46.9 |  |  |
|  |  | TTGATGCTACAATGAAGGAA-------AGCTTTTCCCT | 7.0 | 341 | 3.9 |  |  |  |
|  |  | TTGATGCTACAATGAAG------------CTTTTCCCT | 12.0 | 307 | 3.5 |  |  |  |
|  |  | TTGATGCTACAATGAACGAAC----GAATCTTTTCCCT | 4.0 | 269 | 3.1 |  |  |  |
|  | 4 | TTGATGCTACAATGAAGGA----GCGAAGCTTTTCCCT | 4.0 | 1524 | 19.7 | 42.2 |  |  |
|  |  | TTGATGCTACAATGAAG------------CTTTCCCCT | 12.0 | 417 | 5.4 |  |  |  |
|  |  | TTGATGCTACAATGAAGCAA-------AGCTTTTCCCT | 7.0 | 277 | 3.6 |  |  |  |
|  |  | TTGATGCTACAATGAAGGAA--------GCTTTTCCCT | 8.0 | 228 | 2.9 |  |  |  |
| eTnpBc-$^{HH}$reRNA4$^{PDS}$ | 1 | TTGATGCTACAATGAAGGA----GCGAAGCTTTTCCCT | 4.0 | 943 | 10.4 | 50.8 | 50 | 2 |
|  |  | TTGATGCTACAATGAAGGAAC----GAAGCTTTTCCCT | 4.0 | 498 | 5.5 |  |  |  |
|  |  | TTGATGCTACAATGAAGCAA-------AGCTTTTCCCT | 7.0 | 363 | 4.0 |  |  |  |
|  |  | TTGATGCTACAATGAAG------------CTTTTCCCT | 12.0 | 351 | 3.9 |  |  |  |
|  | 2 | TTGATGCTACAATGAAGGA----GCGAAGCTTTTCCCT | 4.0 | 1190 | 13.3 | 47.5 |  |  |
|  |  | TTGATGCTACAATGAAG------------CTTTTCCCT | 12.0 | 385 | 4.3 |  |  |  |
|  |  | TTGATGCTACAATGAAGGAC-------AGCTTTTCCCT | 7.0 | 350 | 3.9 |  |  |  |
|  |  | TTGATGCTACAATGAAGGAAC----GAAGCTTTTCCCT | 4.0 | 348 | 3.9 |  |  |  |
|  | 3 | TTGATGCTACAATGAAGGA----GCGAAGCTTTTCCCT | 4.0 | 940 | 10.9 | 48.5 |  |  |
|  |  | TTGATGCTACAATGAAGGAAC----GAAGCTTTTCCCT | 4.0 | 562 | 6.5 |  |  |  |
|  |  | TTGATGCGACAATGAAG------------CTTTTCCCT | 12.0 | 401 | 4.7 |  |  |  |
| | | TTGATGCTACAATGAAGGAACTAGC-AAGCTTTTCCCT | 1.0 | 293 | 3.4 | | | |

| | | | | | | | | |
|---|---|---|---|---|---|---|---|---|
|  | 4 | TTGATGCTACAATGAAGGA----GCGAAGCTTTTCCCT | 4.0 | 949 | 9.6 | 52.4 |  |  |
|  |  | TTGATGCTACAATGAAGGAA--------GCT-----CT | 13.0 | 662 | 6.7 |  |  |  |
|  |  | TTGATGCTACGATGAAGGAAC----GAAGCTTTCCCCT | 4.0 | 649 | 6.6 |  |  |  |
|  |  | TTGATGCTACAATGAAG------------CTTTTCCCT | 12.0 | 474 | 4.8 |  |  |  |
| eTnpBe-reRNA4$^{PDS}$ | 1 | TTGATGCTACGATGAAGGAA-------AGCTTTTCCCT | 7.0 | 435 | 6.9 | 39.2 | 43 | 8 |
|  |  | TTGATGCTACAATGAAGGA----GCGAAGCTTTTCCCT | 4.0 | 386 | 6.1 |  |  |  |
|  |  | TTGATGCTACAATGAAG------------CTTTTCCCT | 12.0 | 378 | 6.0 |  |  |  |
|  |  | TTGATGCTACAATGAGGGAA--------GCTTTTCCCT | 8.0 | 264 | 4.2 |  |  |  |
|  | 2 | TTGATGCTACAATGAAGTAA-------AGCTTTTCCCT | 7.0 | 415 | 6.5 | 34.9 |  |  |
|  |  | TTGATGCTACAATGAAG------------CTTTTCCCT | 12.0 | 351 | 5.5 |  |  |  |
|  |  | TTGATGCTACAATGAAGCAA--------GCTTTTCCCT | 8.0 | 342 | 5.4 |  |  |  |
|  |  | TTGATGCTACATTGAAGGA----GCGAAGCTTTTCCCT | 4.0 | 342 | 5.4 |  |  |  |
|  | 3 | TTGATGCTACAATGAAGGA----GCGAAGCTTTTCCCT | 4.0 | 635 | 8.8 | 46.0 |  |  |
|  |  | TTGATGCTACGATGAAGGAA-------AGCTTTTCCCT | 7.0 | 482 | 6.7 |  |  |  |
|  |  | TTGATGCTACAATGAAG------------CTTTTCCCT | 12.0 | 448 | 6.2 |  |  |  |
|  |  | TTGATGCTACAATGAAGGAA--------GCTTGTCCCT | 8.0 | 334 | 4.6 |  |  |  |
|  | 4 | TTGATGCTACAATGAAGGA----GCGAAGCTTTTCCCT | 4.0 | 650 | 7.8 | 52.2 |  |  |
|  |  | TTGATGCTACAATGAAGGAA-------AGCTTTTCCCT | 7.0 | 621 | 7.4 |  |  |  |
|  |  | TTGATGCTACAATGAAG------------CTTTTCCCT | 12.0 | 391 | 4.7 |  |  |  |
|  |  | TTGATGCTACAATGAAGGAA--------GCTTTTCCCT | 8.0 | 297 | 3.6 |  |  |  |
| eTnpBe-$^{HH}$reRNA4$^{PDS}$ | 1 | TTGATGCTACAATGAAGGA----GCGAAGCTTTTCCCT | 4.0 | 612 | 9.6 | 41.4 | 45 | 4 |
|  |  | TTGATGCTACAATGAAG------------CTTTTCCCT | 12.0 | 426 | 6.7 |  |  |  |
|  |  | TTGATGCTACAATGAAGGAA-------AGCTTCTCCCT | 7.0 | 384 | 6.0 |  |  |  |
|  |  | TTGATGCTACAATGAAGGAA--------GCTTTTCCCT | 8.0 | 368 | 5.8 |  |  |  |
|  | 2 | TTGATGCTACAATGAAG------------CTTTTCCCT | 12.0 | 512 | 6.6 | 49.9 |  |  |
|  |  | TTGATGCTACAATGAAGGA----GCGAAGCTTTTCCCT | 4.0 | 460 | 5.9 |  |  |  |
|  |  | TTGATGCTACAATGAAGGAA-------AGCTTTTCCCT | 7.0 | 365 | 4.7 |  |  |  |
|  |  | TTGATGCTACAATGAAGGAG--------GCTTTTCCCT | 8.0 | 262 | 3.4 |  |  |  |
|  | 3 | TTGATGCTACAATGAAGGA----GCGAAGCTTTTCCCT | 4.0 | 454 | 6.5 | 45.1 |  |  |
|  |  | TTGATGCTACAATGAAGGAA-------AGCTTTTCCCT | 7.0 | 428 | 6.1 |  |  |  |
|  |  | TTGATGCTACAATGAAG------------CTTTTCCCT | 12.0 | 379 | 5.4 |  |  |  |
|  |  | TTGATGCTACAATGAAGGAAC----GAAGCTTTTCCCT | 4.0 | 283 | 4.0 |  |  |  |
|  | 4 | TTGATGCTACAATGAAGGA----GCGAAGCTTTTCCCT | 4.0 | 738 | 9.3 | 44.9 |  |  |
|  |  | TTGATGCTACAATGAAGGAA-------AGCTTTTCCCT | 7.0 | 618 | 7.8 |  |  |  |
|  |  | TTGATGCTGCAATGAAG------------CTTTTCCCT | 12.0 | 418 | 5.3 |  |  |  |
|  |  | TTGATGCTGCAATGAAGGAA--------GCTTTTCCCT | 8.0 | 385 | 4.9 |  |  |  |

*Bold red letters indicate TAM, and underlined letters denote reRNA target sequences

**Supplementary Table 6.** Efficiency of somatic editing at the *PDS* locus in systemic leaves targeted by TnpB and TnpB variants with reRNA4$^{PDS}$

| Construct | Plant # | Most common indels | n_deleted | Reads | % Edited reads | Overall editing frequency | Average editing frequency | Standard deviation |
|---|---|---|---|---|---|---|---|---|
| | | TTGATGCTACAATGAAGGAACTAGCGAAGCTTTTCCCT* | Wild-type | | | | | |
| TnpB-reRNA4$^{PDS}$ | 1 | TTGATGGTACAATGAAGGA----GCGAAGCTTTTCCCT | 4 | 33 | 7.5 | 3.7 | 3 | 1 |
|  |  | TTGATGCTACAATGAAG------------CTTTTCCCT | 12 | 32 | 7.3 |  |  |  |
|  |  | TTGATGCTACAATGAAGGAA-------AGCTTTTCCCT | 7 | 27 | 6.1 |  |  |  |
|  |  | TTGATGCTACAATGATGGAAC----GAAGCTTTTCCCT | 4 | 23 | 5.2 |  |  |  |
|  | 2 | TTGATGCTACAA-------------------------- | 59 | 14 | 9.3 | 1.4 |  |  |
|  |  | TTGATGCTACAATGAAG------------CTTTTCCCT | 12 | 11 | 7.3 |  |  |  |
| | | TTGATGCTACAATGAAGGAACTAGCAAAGCTTT-CCCT | 1 | 7 | 4.7 | | | |
|  |  | TTGATG--------------------AAGCTTTTCCCT | 20 | 7 | 4.7 |  |  |  |
|  | 3 | TTGATGCTACAATGAAGGA----GCGAAGCTTTTCCCT | 4 | 29 | 6.6 | 3.8 |  |  |
|  |  | TTGATGCTACAATGAAGGAA--------GCTTTTCCCT | 8 | 29 | 6.6 |  |  |  |
|  |  | TTGATGGTACAATGAAG------------CTTTTCCCT | 12 | 26 | 5.9 |  |  |  |
|  |  | TTGATGCTACAATGAAGGAACTA----AGCTTTTCCCT | 4 | 19 | 4.3 |  |  |  |
|  | 4 | TTGATGCTACAATGAAGGAA-------AGCTTTTCCCT | 7 | 18 | 11.5 | 2.0 |  |  |
|  |  | TTGATGCTACAATGAAGGAAC----GAAGCTTTTCCCT | 4 | 11 | 7.1 |  |  |  |
|  |  | TTGATGCCACACTGAAG------------CTTTTCCCT | 12 | 11 | 7.1 |  |  |  |
|  |  | TTGATGCTA------------------AGCTTTTCCCT | 18 | 7 | 4.5 |  |  |  |
| TnpB-$^{HH}$reRNA4$^{PDS}$ | 1 | TTGATGCTACAATGAAG------------CTTTTCCCT | 12 | 423 | 9.4 | 18.6 | 12 | 4 |
|  |  | TTGATGCTACAATGAAGGAA--------GCTTTTCCCT | 8 | 319 | 7.1 |  |  |  |
|  |  | TTGATGCTACAATGAAGGAA-------AGCTTTTCCCT | 7 | 252 | 5.6 |  |  |  |
|  |  | TTGATGCTACAATGAAGGAAC----GAAGCTTTTCCCT | 4 | 246 | 5.5 |  |  |  |
|  | 2 | TTGATGCTACAATGAAG------------CTTTTCCCT | 12 | 303 | 12.1 | 11.5 |  |  |
|  |  | TTGATGCTACAATGA-------------GCTTTTCCCT | 13 | 163 | 6.5 |  |  |  |
|  |  | TTGATGCTACAATGAAGGA----GCAAAGCTTTTCCCT | 4 | 152 | 6.0 |  |  |  |
|  |  | TTGATGCTACAAAGAAGGAAC----GAAGCTTCTCCCT | 4 | 146 | 5.8 |  |  |  |
|  | 3 | TTGATGCTACAATGAAG------------CTTTTCCCT | 12 | 152 | 10.0 | 9.8 |  |  |
|  |  | TTGATGCTACAATGAAGGAAC----GAAGCTTTTCCCT | 4 | 108 | 7.1 |  |  |  |
|  |  | TTGATGCTACAATGA-------------GCTTTTCCCT | 13 | 97 | 6.4 |  |  |  |
|  |  | TTGATGCTACAATGAAGGAA--------GCTTTTCCCT | 8 | 85 | 5.6 |  |  |  |
|  | 4 | TTGATGCTACAATGAAG------------CTTTTCCCT | 12 | 220 | 12.5 | 8.8 |  |  |
|  |  | TTGATGCTACAATGAAGGAAC----GAAGCTTTTCCCT | 4 | 180 | 10.2 |  |  |  |
|  |  | TTGATGCTACAATTA-------------GCTTTTCCCT | 13 | 161 | 9.1 |  |  |  |
|  |  | TTGATGCCACAATGAAGGA----GCGAAGCTTTTCCCT | 4 | 134 | 7.6 |  |  |  |
| eTnpBc-reRNA4$^{PDS}$ | 1 | TTGATGCTACAATGAAGGAA-------AGCTTTTCCCT | 7 | 519 | 9.4 | 68.9 | 71 | 9 |
|  |  | TTGATGCTACACTGAAGGAAC------AGCTTTTCCCT | 6 | 482 | 8.7 |  |  |  |
|  |  | TTGATGCTACAATGAAGGAA--------GCTTTTCCCT | 8 | 410 | 7.4 |  |  |  |
|  |  | TTGATG-----------------------CTTTTCCCT | 26 | 382 | 6.9 |  |  |  |
|  | 2 | TTGATGCTACAATGAAG------------CTTTTCCCT | 12 | 392 | 7.6 | 74.4 |  |  |
|  |  | TTGATGCTACAATGAAGGAA-------AGCTTTTCCTT | 7 | 385 | 7.4 |  |  |  |
|  |  | TTGATGCTACAATGAAGGA----GCGAAGCTTTTCCCT | 4 | 368 | 7.1 |  |  |  |
|  |  | TTGATGCTATAATGAAGGAAC-----AAGCTTTTCCCT | 5 | 201 | 3.9 |  |  |  |
|  | 3 | TTGATGCCACAATGAAG------------CTTTTCCCT | 12 | 523 | 7.9 | 79.7 |  |  |
|  |  | TTGATGCTACAATGA----------------------- | 25 | 493 | 7.4 |  |  |  |
|  |  | TTGATGCTACAATGAAGGAGC-----AAGCTTTTCCCT | 5 | 388 | 5.8 |  |  |  |
|  |  | -------------------------------------- | 55 | 303 | 4.6 |  |  |  |
|  | 4 | TTGATGCTACAATGAAGGAAC------AGCTTTTCCCT | 6 | 656 | 16.6 | 59.6 |  |  |
|  |  | TTGATGCTACAATGAAGCAA-------AGCTTTTCCTT | 7 | 600 | 15.2 |  |  |  |
|  |  | TTGATGCTACAATGAAGGA-------------TTCCCT | 13 | 600 | 15.2 |  |  |  |
|  |  | TTGATGCTACAATGAGGGA---------GCTTTTCCCT | 9 | 471 | 11.9 |  |  |  |
| eTnpBc-$^{HH}$reRNA4$^{PDS}$ | 1 | TTGATGCTACAATGAAGCAA-------AGCTTTTCCCT | 7 | 361 | 16.2 | 47.9 | 62 | 17 |
|  |  | TTGATGCAACAATGAAGGAAC------AGCTTTTCCCT | 6 | 286 | 12.8 |  |  |  |
|  |  | TTGATGCTACAATGAAGGA-------------TTCCCT | 13 | 283 | 12.7 |  |  |  |
|  |  | ------------------------CAAAGCTTTTCCCT | 38 | 233 | 10.5 |  |  |  |
|  | 2 | -----------------------------CTTTTCCCT | 26 | 1182 | 14.9 | 77.5 |  |  |
|  |  | TTGATGCTACAATGAAGGAAC------AGCTTTTCCCT | 6 | 1099 | 13.8 |  |  |  |
|  |  | TTGATGCTACAATGAAGGAA-------AGCTTTTCCCT | 7 | 1051 | 13.2 |  |  |  |
|  |  | TTGATGCTACAATGAAGGAACGAGC-----TTTTCCCT | 5 | 1026 | 12.9 |  |  |  |
|  | 3 | TTGATGCTACAATGAAGGAACAAGC------TTTCCCT | 6 | 563 | 16.4 | 46.9 |  |  |
|  |  | TTGATGCTACAATGAAGGAA-------AGCTTTTCCGT | 7 | 557 | 16.2 |  |  |  |
|  |  | TTGATGCTACAATGAAG-----AGC----------CCT | 15 | 488 | 14.2 |  |  |  |
|  |  | TTGATGCTACAATGAAGGAAC-----AAGCTTTTCCCT | 5 | 439 | 12.8 |  |  |  |

| | | | | | | | | |
|---|---|---|---|---|---|---|---|---|
|  | 4 | TTGATGCTACAATGAAGGAA-------AGCTTTTCCCT | 7 | 767 | 15.3 | 74.8 |  |  |
|  |  | ------------------------CAAAGCTTTTCCCT | 38 | 553 | 11.0 |  |  |  |
|  |  | TTGATGCTGCAATGAAGGAA------------------ | 21 | 445 | 8.9 |  |  |  |
|  |  | TTGATGCTACAATGAAGG--------------TTCAC- | 14 | 438 | 8.7 |  |  |  |
| eTnpBe-reRNA4^PDS | 1 | TTGATGCTACAATGAAGGA----GCGAAGCTTTTCCCT | 4 | 59 | 9.2 | 29.7 | 26 | 4 |
|  |  | TTGATGCTACAATGAAGGAA-------AGCTTTTCCCT | 7 | 46 | 7.2 |  |  |  |
|  |  | TTGATGCTACAATGAAG------------CTTTTCCCT | 12 | 32 | 5.0 |  |  |  |
|  |  | -------------------------------------- | 65 | 27 | 4.2 |  |  |  |
|  | 2 | TTGATGCTACAATGAAGGAA-------AGCTTTTCCCT | 7 | 36 | 8.3 | 19.6 |  |  |
|  |  | TTGATGCTACAATGAAGGAA--------GCTTTTCCCT | 8 | 31 | 7.2 |  |  |  |
|  |  | TTGATGCTACAATGAAG------------CTTTTCCCT | 12 | 25 | 5.8 |  |  |  |
|  |  | TTGATGCTACAATGAAGGA----GCAAAGCTTTTCCCT | 4 | 22 | 5.1 |  |  |  |
|  | 3 | TTGATGCTACAATGAAGGAA-------AGCTTTTCCCT | 7 | 80 | 12.6 | 26.7 |  |  |
|  |  | TTGATGCTACAATGAAGGAG--------GCTTTTCCCT | 8 | 54 | 8.5 |  |  |  |
|  |  | TTGATGCTACAATGAAGGA----GCAAAGCTTTTCCCT | 4 | 29 | 4.6 |  |  |  |
|  |  | TTGATGCTACAATGAAG------------CTTTTCCCT | 12 | 27 | 4.3 |  |  |  |
|  | 4 | TTGATGCTACAATGAAGGAA--------GCTTTTCCCT | 8 | 69 | 11.2 | 25.9 |  |  |
|  |  | TTGATGCTACAATGAAGGA----GCGAAGCTTTTCCCT | 4 | 62 | 10.0 |  |  |  |
|  |  | TTGATGCTACAATGAAGGAA-------AGCTTTTCCCT | 7 | 55 | 8.9 |  |  |  |
|  |  | TTGATGCTACAATGAAGGAACTA----AGCTTTTCCCT | 4 | 26 | 4.2 |  |  |  |
| eTnpBe-^HHreRNA4^PDS | 1 | TTGATGCTACAATGAAG------------CTTTTCCCT | 12 | 40 | 9.5 | 17.4 | 20 | 4 |
|  |  | TTGATGCTACAATGAAGGAA--------GCTTTTCCCC | 8 | 35 | 8.3 |  |  |  |
|  |  | TTGATGCTACAATGAAGGAA-------AGCTTTCCCCT | 7 | 30 | 7.1 |  |  |  |
|  |  | TTGATGCTACACTGAAGGA----GCAAAGCTTTTCCCT | 4 | 26 | 6.1 |  |  |  |
|  | 2 | TTGATGCTACAATGAAGGA----GCGAAGCTTTTCCCT | 4 | 60 | 11.1 | 19.3 |  |  |
|  |  | TTGATGCTACAATGAAGCAA-------AGCTTTTCCCT | 7 | 28 | 5.2 |  |  |  |
|  |  | TTGATGCTACAATGAAG------------CTTTTCCCT | 12 | 27 | 5.0 |  |  |  |
|  |  | TTGATGCTACAATGAAGGAAC----GAAGCTTTTCCCT | 4 | 27 | 5.0 |  |  |  |
|  | 3 | TTGATGCTACAATGAAGGAA--------GCTTTTCCCT | 8 | 97 | 9.2 | 25.4 |  |  |
|  |  | TTGATGCTACAATGAAGGA---------GCTTTTCCCT | 9 | 75 | 7.1 |  |  |  |
|  |  | TTGATGCTACAATGAAGGAA-------AGCTTTTCCCT | 7 | 73 | 6.9 |  |  |  |
|  |  | TTGATGCTACAATGAAGGA----GCGAAGCTTTTCCCT | 4 | 71 | 6.7 |  |  |  |
|  | 4 | TTGATGCTACAATGAAGGA----GCGAAGCTTTTCCCT | 4 | 42 | 8.7 | 18.8 |  |  |
|  |  | TTGATGCTACAATGAAGGAA-------AGCTTTTCCCT | 7 | 38 | 7.9 |  |  |  |
|  |  | TTGATGCTACAATGAAGGAA--------GCTTTTCACT | 8 | 34 | 7.0 |  |  |  |
|  |  | TTGATGCTACAATGAAG------------CTTTTCCCT | 12 | 31 | 6.4 |  |  |  |

*Bold red letters indicate TAM, and underlined letters denote reRNA target sequences

**Supplementary Table 7.** Efficiency of somatic editing in infiltrated leaves at the *PDS* locus targeted by TnpB and eTnpBc fused to 2xSV40nls with reRNA4$^{PDS}$

| Construct | Plant # | Most common indels | n_deleted | Reads | % Edited reads | Overall editing frequency | Average editing frequency | Standard deviation |
|---|---|---|---|---|---|---|---|---|
| | | **TTGAT**GCTACAATGAAGGAACTAGCGAAGCTTTTCCCT* | Wild-type | | | | | |
| TnpB$_{2xSV40nls}$-reRNA4$^{PDS}$ | 1 | TTGATGCTACAATGAAGGAA-------AGCTTTTCCCT | 7 | 76 | 7.3 | 3.3 | 7 | 8 |
|  |  | TTGATGCTACAATGAAGGA----GCAAAGCTTTTCCCT | 4 | 73 | 7.0 |  |  |  |
|  |  | TTGATGCTACAATGAAG------------CTTTTCCCT | 12 | 58 | 5.5 |  |  |  |
|  |  | TTGATGCTACAATGAA-----------AGCTTTTCCCT | 11 | 37 | 3.5 |  |  |  |
|  | 2 | TTGATGCTACAATGAAG------------CTTTTCCCT | 12 | 38 | 7.2 | 1.2 |  |  |
|  |  | TTGATGCTACAATGAAGGAA--------GCTTTTCCCT | 8 | 37 | 7.0 |  |  |  |
| | | TTGATGCTACAATGAAGGAACTAGCAAAGCTTT-CCCT | 1 | 34 | 6.5 | | | |
|  |  | -------------------------------------- | 74 | 27 | 5.1 |  |  |  |
|  | 3 | TTGATGCTACAATGAAGGA----GCGAAGCTTTTCCCT | 4 | 171 | 10.9 | 3.6 |  |  |
|  |  | TTGATGCCACAATGAAG------------CTTTTCCCT | 12 | 79 | 5.1 |  |  |  |
|  |  | TTGATGCTACAATGAAGCAA--------GCTTTTCCCT | 8 | 68 | 4.4 |  |  |  |
|  |  | TTGATGCTACAATGAA-----------AGCTTTTCCCT | 11 | 60 | 3.8 |  |  |  |
|  | 4 | TTGATGCTACAATGAAGGAA--------GCTTTTCCCT | 8 | 628 | 10.4 | 18.3 |  |  |
|  |  | TTGATGCTACAATGAAG------------CTTTTCCCT | 12 | 414 | 6.8 |  |  |  |
|  |  | TTGATGCTACAATGAAGGAA-------AGCTTTTCCCT | 7 | 372 | 6.1 |  |  |  |
|  |  | TTGATGCTACAATGAAGGA----GCGAAGCTTTTCCCT | 4 | 305 | 5.0 |  |  |  |
| TnpB$_{2xSV40nls}$-$^{HH}$reRNA4$^{PDS}$ | 1 | TTGATGCTACAATGAAGGA----GCGAAGCTTTTCCCT | 4 | 86 | 9.0 | 3.3 | 6 | 7 |
|  |  | TTGATGCTACAATGAAG------------CTTTTCCCT | 12 | 62 | 6.5 |  |  |  |
| | | TTGATGCTACAATGAAGGAACTAGCGAAGCTTT-CCCT | 1 | 41 | 4.3 | | | |
|  |  | TTGATGCTACAATGAAGCAA-------AGCTTTTCCCT | 7 | 37 | 3.9 |  |  |  |
| | 2 | TTGATGCTACAATGAAGGAACTAGCAAAGCTTT-CCCT | 1 | 35 | 7.2 | 1.1 | | |
|  |  | TTGATGCTACAATGAAGCAA-------AGCTTTTCCCT | 7 | 26 | 5.3 |  |  |  |
|  |  | -------------------------------------- | 74 | 20 | 4.1 |  |  |  |
|  |  | TTGATGCTACAA-------------------------- | 59 | 20 | 4.1 |  |  |  |
|  | 3 | TTGATGCTACAATGAAGGA----GCGAAGCTTTTCCCT | 4 | 149 | 10.1 | 3.6 |  |  |
|  |  | TTGATGCTACAATGAAGGAA--------GCTTTTCCCT | 8 | 88 | 6.0 |  |  |  |
|  |  | TTGATGCTACAATGTAG------------CTTTTCCCT | 12 | 73 | 4.9 |  |  |  |
|  |  | TTGATGCTACAATGAAGGAAC----GAAGCTTTTCCCT | 4 | 59 | 4.0 |  |  |  |
|  | 4 | TTGATGCTACAATGAAGGAA--------GCTTTTCCCT | 8 | 433 | 8.4 | 17.5 |  |  |
|  |  | TTGATGCTACAATGAAGGAA-------AGCTTTTCCCT | 7 | 351 | 6.8 |  |  |  |
|  |  | TTGATGCTACAATGAAG------------CTTTTCCCT | 12 | 333 | 6.5 |  |  |  |
|  |  | TTGATGCTACAATGAAGGA----GCGAAGCTTTTCCCT | 4 | 299 | 5.8 |  |  |  |
| eTnpBc$_{2xSV40nls}$-reRNA4$^{PDS}$ | 1 | TTGATGCTACAATGAAG------------CTTTTCCCT | 12 | 871 | 16.2 | 28.1 | 35 | 12 |
|  |  | TTGATGATACAATGA-------------GCTTTTCCCC | 13 | 640 | 11.9 |  |  |  |
|  |  | TTGATGCTACAATGAAGGAAC----GAAGCTTTTCCCT | 4 | 622 | 11.6 |  |  |  |
|  |  | TTGATGCTACAATGAAGGAA--------GCTTCTCCCT | 8 | 454 | 8.4 |  |  |  |
|  | 2 | TTGATGAATGAAG------------CTTTTCCCTTTCG | 12 | 373 | 13.2 | 21.1 |  |  |
|  |  | TTGATGAATGAAGGA----GCGAAGCTTTTCCCTTTCG | 4 | 306 | 10.8 |  |  |  |
|  |  | TTGATGAATGAAGGAAC----GAAGCTTTTCCCTTTCG | 4 | 259 | 9.1 |  |  |  |
|  |  | TTGATGAATGA-------------GCTTTTCCCTTTCA | 13 | 215 | 7.6 |  |  |  |
|  | 3 | TTGATGCTACAATGAAG------------CTTTTCCCT | 12 | 1586 | 19.8 | 45.8 |  |  |
|  |  | TTGATGCTACAATGA-------------GCTTTTCCCT | 13 | 1277 | 16.0 |  |  |  |
|  |  | TTGATGCTTCAATGAAGGAAC----GAAGCTTTTCCCT | 4 | 1065 | 13.3 |  |  |  |
|  |  | TTGATGCTACAATGAAGGAA--------GCTTTTCCCT | 8 | 676 | 8.5 |  |  |  |
|  | 4 | TTGATGCTACAATGAAG------------CTTTTCCCT | 12 | 851 | 19.9 | 43.4 |  |  |
|  |  | TTGATGCTACAATGAAGGAAC----GAAGCTTTTCCCT | 4 | 572 | 13.4 |  |  |  |
|  |  | TTGATGCTACAATGA-------------GCTTTTCCTT | 13 | 571 | 13.3 |  |  |  |
|  |  | TTGATGCTACAATGAAGGAA--------GCTTTTCCCT | 8 | 365 | 8.5 |  |  |  |
| eTnpBc$_{2xSV40nls}$-$^{HH}$reRNA4$^{PDS}$ | 1 | TTGATGCTACAATGAAG------------CTTTTCCCT | 12 | 965 | 10.4 | 39.6 | 38 | 4 |
|  |  | TTGATGCTACAATGAAGGA----GCGAAGCTTTTCCCT | 4 | 653 | 7.0 |  |  |  |
|  |  | TTGATGCTACAATGAAGGAA-------AGCTTTTCCCT | 7 | 555 | 6.0 |  |  |  |
|  |  | TTGATGCCACAATGAAGGAA--------GCTTTTCCCT | 8 | 514 | 5.5 |  |  |  |
|  | 2 | TTGATGCTACAATGAAG------------CTTTTCCCT | 12 | 1307 | 14.4 | 34.8 |  |  |
|  |  | TTGATGCTACAATGAAGGAA-------AGCTTTTCCCT | 7 | 582 | 6.4 |  |  |  |
|  |  | TTGATGCTACAATGAAGGAAC----GAAGCTTTTCCCT | 4 | 575 | 6.3 |  |  |  |
|  |  | TTGATGCTACAATGA-------------GCTTTTCCCT | 13 | 546 | 6.0 |  |  |  |
| | 3 | TTGATGCTACAATGAAGGAACTAGC-AAGCTTTTCCCT | 1 | 2936 | 31.6 | 43.5 | | |
|  |  | TTGATGCTACAATGACG------------CTTTCCCCT | 12 | 750 | 8.1 |  |  |  |
|  |  | TTGATGCTACAATGA-------------GCTTTTCCCT | 13 | 455 | 4.9 |  |  |  |
|  |  | TTGATGCTACAATGAATGAA-------AGCTTTTCCCT | 7 | 426 | 4.6 |  |  |  |
|  | 4 | TTGATGCTACAATGAAG------------CTTTTCCCT | 12 | 1491 | 15.0 | 34.8 |  |  |
|  |  | TTGATGCTACAATGAAGGA----GCGAAGCTTTTCCCT | 4 | 620 | 6.2 |  |  |  |
|  |  | TTGATGCTACAATGAAGGAAC----GAAGCTTTTCCCT | 4 | 612 | 6.1 |  |  |  |
|  |  | TTGATGCTACAATGGAGGAA-------AGCTTTTCCCT | 7 | 588 | 5.9 |  |  |  |

*Bold red letters indicate TAM, and underlined letters denote reRNA target sequences

**Supplementary Table 8.** Efficiency of somatic editing at the *PDS* locus in systemic leaves targeted by eTnpBc$_{2xSV40nls}$ with reRNA4$^{PDS}$

| Construct | Plant # | Most common indels | n_deleted | Reads | % Edited reads | Overall editing frequency | Average editing frequency | Standard deviation |
|---|---|---|---|---|---|---|---|---|
| | | **TTGAT**GCTACAATGAAGGAACTAGCGAAGCTTTTCCCT* | Wild-type | | | | | |
| eTnpBc$_{2xSV40nls}$-reRNA4$^{PDS}$ | 1 | TTGATGCTACAATGAAG------------CTTTTCCCT | 12 | 1595 | 9.0 | 79.16 | 90 | 8 |
|  |  | TTGATGCTATAATGAAGGAA-------AGCTTTTCCCT | 7 | 1431 | 8.1 |  |  |  |
|  |  | TTGATGCTACAATGAAAGG-----------------CT | 17 | 829 | 4.7 |  |  |  |
|  |  | TTGATGCTACAATGAAGGA----GCAAAGCTTATCCCT | 4 | 821 | 4.6 |  |  |  |
|  | 2 | TTGATGCTACAAT------------------------- | 26 | 3666 | 14.8 | 95.92 |  |  |
|  |  | TTGATGCTACAATGAAGGAA-------AGCTTTTCCCT | 7 | 3535 | 14.3 |  |  |  |
|  |  | TTGATGCTACAATGAAGGAAC----GAAGCTTTTCCCT | 4 | 2467 | 10.0 |  |  |  |
|  |  | TTGATGCTACAATGAATGA------------------- | 19 | 1700 | 6.9 |  |  |  |
|  | 3 | TTGATGCTACAAT------------------------- | 26 | 4313 | 14.9 | 95.88 |  |  |
|  |  | TTGATGCTACAATGAAGGAA-------AGCTTTTCCCT | 7 | 3182 | 11.0 |  |  |  |
|  |  | TTGATGCTACAATGAAGGAAC----GAAGCTTTTCCCT | 4 | 2702 | 9.3 |  |  |  |
|  |  | TTGATGTTACAATGAAGGA------------------- | 19 | 1870 | 6.4 |  |  |  |
|  | 4 | TTGATGCTACAATGAAGGA----GGGAAGCTTTTCCCT | 4 | 2261 | 6.8 | 89.33 |  |  |
|  |  | TTGATGCTACAATGA--------GCAAAGCTTTTCCCT | 8 | 1833 | 5.5 |  |  |  |
|  |  | TTGATGCTACAATGAAGCAA-------AGCTTTTCCCT | 7 | 1363 | 4.1 |  |  |  |
|  |  | -------------------------------------- | 48 | 1127 | 3.4 |  |  |  |
| eTnpBc$_{2xSV40nls}$-$^{HH}$reRNA4$^{PDS}$ | 1 | TTGATGCTACAATGAAG------------CTTTTCCCT | 12 | 2309 | 9.2 | 79 | 90 | 10 |
|  |  | TTGATGCTACAATGGAGGAA-------AGCTTTTCCCT | 7 | 2135 | 8.5 |  |  |  |
|  |  | TTGATGCTACAATGAAGGA-----------------CT | 17 | 1167 | 4.7 |  |  |  |
|  |  | TTGATGCTACAATGAAGGA----GCGAAGCTTTTCCCT | 4 | 1142 | 4.6 |  |  |  |
|  | 2 | TTGATGCTACAAT------------------------- | 26 | 5709 | 16.3 | 96 |  |  |
|  |  | TTGATGCTACAATGAAGGAA-------AGCTTTTCCCT | 7 | 4963 | 14.2 |  |  |  |
|  |  | TTGATGCTACAATGAAGGAAC----GAAGCTTTTCCCT | 4 | 3654 | 10.4 |  |  |  |
|  |  | TTGATGCTACAATGTAGGA------------------- | 19 | 2587 | 7.4 |  |  |  |
|  | 3 | TTGATGCTACAAT------------------------- | 26 | 7237 | 14.4 | 96.37 |  |  |
|  |  | TTGATGCTACAATGAATGAA-------AGCTTTTCCCT | 7 | 5568 | 11.1 |  |  |  |
|  |  | TTGATGCTACATTGAAGGAAC----GAAGCTTTTCCCT | 4 | 5043 | 10.0 |  |  |  |
|  |  | TTGATGCTACAATGAAGGA------------------- | 19 | 3171 | 6.3 |  |  |  |

*Bold red letters indicate TAM, and underlined letters denote reRNA target sequences

**Supplementary Table 9.** Efficiency of somatic editing at the *PDS* locus in infiltrated leaves targeted by *N. benthamiana* codon optimized (NbCo) TnpB and TnpB variants with reRNA4$^{PDS}$

| Construct | Plant # | Most common indels | n_deleted | Reads | % Edited reads | Overall editing frequency | Average editing frequency | Standard deviation |
|---|---|---|---|---|---|---|---|---|
| | | **TTGAT**GCTACAATGAAGGAACTAGCGAAGCTTTTCCCT* | Wild-type | | | | | |
| NbCoTnpB-reRNA4$^{PDS}$ | 1 | TTGATGCTACAATGAAGGAACTAGCAAAGCTTT-CCCT | 1 | 180 | 7.7 | 5.4 | 7 | 8 |
|  |  | TTGATGCTACAATTAAGGA----GCGAAGCTTTTCCCT | 4 | 146 | 6.3 |  |  |  |
|  |  | TTGATGCTACAATGGAGGAT--------GCTTTTCCCT | 8 | 112 | 4.8 |  |  |  |
|  |  | -------------------------------------- | 45 | 107 | 4.6 |  |  |  |
| | 2 | TTGATGCTACAATGAAGGAACTAGCAAAGCTTT-CCCT | 1 | 75 | 18.7 | 1.5 | | |
|  |  | TTGATGCTACAATGAAGGAA-------AGCTTTTCCCT | 7 | 38 | 9.5 |  |  |  |
|  |  | TTGATGCTACAATGAAGGA----GCGAAGCTTTTCCCT | 4 | 24 | 6.0 |  |  |  |
|  |  | TTGATGCTACAATGAAGGAA--------GCTTTTCCCT | 8 | 15 | 3.7 |  |  |  |
| | 3 | TTGATGCTACAATGAAGGAACTAGCAAAGCTTT-CCCT | 1 | 63 | 8.0 | 3.2 | | |
|  |  | TTGATGCTACAATGAAGGAA-------AGCTTTCCCCT | 7 | 57 | 7.2 |  |  |  |
|  |  | TTGATGCTACAATGAA-----------AGCTTTTCCCT | 11 | 48 | 6.1 |  |  |  |
|  |  | TTGATGCCACAATGAAG------------CTTTTCCCT | 12 | 47 | 5.9 |  |  |  |
|  | 4 | TTGATGCTACAATGAAGGAA--------GCTTTTCCCT | 8 | 298 | 6.9 | 19.8 |  |  |
|  |  | TTGATGCTACAATGAAGGA----GCGAAGCTTTTCCCT | 4 | 265 | 6.1 |  |  |  |
|  |  | TTGATGCTACAATGAAG------------CTTTTCCCT | 12 | 226 | 5.2 |  |  |  |
|  |  | TTGATGCTACAATGAAGGAA-------AGCTTTTCCCT | 7 | 222 | 5.1 |  |  |  |
| NbCoTnpB-$^{HH}$reRNA4$^{PDS}$ | 1 | TTGATGCTACAATGAAGGAACTAGCAAAGCTTT-CCCT | 1 | 218 | 8.7 | 5.5 | 7 | 8 |
|  |  | TTGATGCTACAATGAAGGA----GCGAAGCTTTTCCCT | 4 | 211 | 8.4 |  |  |  |
|  |  | -------------------------------------- | 48 | 133 | 5.3 |  |  |  |
|  |  | TTGATGCTACAATGAAG------------CTTTTCCCT | 12 | 111 | 4.4 |  |  |  |
| | 2 | TTGATGCTACAATGAAGGAACTAGCAAAGCTTT-CCCT | 1 | 95 | 18.8 | 1.8 | | |
|  |  | TTGATGCTACAATGAAGGAA-------AGCTTTTCCCT | 7 | 34 | 6.7 |  |  |  |
|  |  | TTGATGCTACAATGAAGGA----GCAAAGCTTTTCCCT | 4 | 30 | 5.9 |  |  |  |
|  |  | TTGATGCTACAATGAAGGAA--------GCTTTTCCCT | 8 | 30 | 5.9 |  |  |  |
|  | 3 | TTGATGCTACAATGAAGGAA--------GCTTTTCCCT | 8 | 73 | 9.0 | 3.2 |  |  |
| | | TTGATGCTACAATGAAGGAACTAGCAAAGCTTT-CCCT | 1 | 50 | 6.2 | | | |
|  |  | TTGATGCTACAATGAAGGA----GCGAAGCTTTTCCCT | 4 | 44 | 5.4 |  |  |  |
|  |  | -------------------------------------- | 65 | 41 | 5.0 |  |  |  |
|  | 4 | TTGATGCTACAATGAAGGAA--------GCTTTTCCCT | 8 | 396 | 8.2 | 19.0 |  |  |
|  |  | TTGATGCTACAATGAAG------------CTTTTCCCT | 12 | 288 | 5.9 |  |  |  |
|  |  | TTGATGCTACAATGAAGGAA-------AGCTTTTCCCT | 7 | 261 | 5.4 |  |  |  |
|  |  | TTGATGCTACAATGAAGGA----GCGAAGCTTTTCCTT | 4 | 257 | 5.3 |  |  |  |
| NbCoeTnpBc-reRNA4$^{PDS}$ | 1 | TTGATGCTACAATGAAGGA----GCGAAGCTTTTCCCT | 4 | 99 | 4.9 | 52.6 | 55 | 8 |
|  |  | TTGATGCTACAATGATG------------CTTTTCCCT | 12 | 69 | 3.4 |  |  |  |
|  |  | TTGATGCTACAATGA----------------------- | 33 | 56 | 2.7 |  |  |  |
|  |  | TTGATGCTACAATGAAA--------------------- | 29 | 50 | 2.4 |  |  |  |
|  | 2 | TTGATGCTACAATGAAGGA----GCGAAGCTTTTCCCT | 4 | 199 | 4.8 | 65.0 |  |  |
|  |  | TTGATGCTACAATGAAG------------CTTTTCCCT | 12 | 182 | 4.4 |  |  |  |
|  |  | TTGATGCTACAATGAA-----------AGCTTTTCCCT | 11 | 136 | 3.3 |  |  |  |
|  |  | TTGATGCTACAATGAAGGAC-------AGCTTTTCCCT | 7 | 107 | 2.6 |  |  |  |
|  | 3 | TTGATGCTACAATGAAGGA----GCGAAGCTTTTCCCT | 4 | 302 | 7.9 | 57.7 |  |  |
|  |  | TTGATGCTACAATTTCG------------CTTTTCCCT | 12 | 166 | 4.3 |  |  |  |
|  |  | TTGATGCTACAATGAAGGAA-------AACTTTTCCCT | 7 | 135 | 3.5 |  |  |  |
|  |  | TTGATGCTACAATGAAGGAA--------GCTTTTCCCT | 8 | 122 | 3.2 |  |  |  |
|  | 4 | TTGATGCTACAATGAAGGA----GCGAAGCTTTTCCCT | 4 | 71 | 5.7 | 46.2 |  |  |
|  |  | TTGATGCTACAATGAAGGAG-------AGCTTTTCCCT | 7 | 69 | 5.5 |  |  |  |
|  |  | TTGATGCTACAATGAAGGAA--------GCTTTTCCCT | 8 | 58 | 4.7 |  |  |  |
|  |  | -------------------------------------- | 67 | 48 | 3.8 |  |  |  |
| NbCoeTnpBc-$^{HH}$reRNA4$^{PDS}$ | 1 | TTGATGCTACAATGAAGGAA-------AGCTTTTCCCT | 7 | 72 | 6.9 | 57.1 | 56 | 6 |
|  |  | TTGATGCTACAATGAAGGA----GCGAAGCTTTTCCCT | 4 | 62 | 5.9 |  |  |  |
|  |  | TTGATGCTACAATGAAG------------CTTTTCCCT | 12 | 61 | 5.8 |  |  |  |
|  |  | TTGATGCTACAATGAAGGAC--------GCTTTTCCCT | 8 | 45 | 4.3 |  |  |  |
|  | 2 | TTGATGCTACAATGAAG------------CTTTTCCCT | 12 | 124 | 6.4 | 57.1 |  |  |
|  |  | TTGATGCTACAATGAAGGA----GCGAAGCTTTTCCCT | 4 | 110 | 5.7 |  |  |  |
|  |  | TTGATGCTACAATGAA-----------AGCTTTTCCCT | 11 | 62 | 3.2 |  |  |  |
|  |  | TTGATGCTACAATGAAGGAA-------AGCTTTTCCCT | 7 | 47 | 2.4 |  |  |  |
|  | 3 | TTGATGCTACAATGAAGGA----GCGAAGCTTTTCCCT | 4 | 88 | 6.6 | 47.7 |  |  |
|  |  | TTGATGCTACAATGAAG------------CTTTTCCCT | 12 | 69 | 5.1 |  |  |  |
|  |  | -------------------------------------- | 51 | 63 | 4.7 |  |  |  |
|  |  | TTGATGGTACAATGAAAGAA--------GCTTTTCCCT | 8 | 59 | 4.4 |  |  |  |
|  | 4 | TTGATGCTACAATGAAGGA----GCGAAGCTTTTCCCT | 4 | 149 | 6.8 | 62.4 |  |  |
|  |  | TTGATGCTACAATGAAG------------CTTTTCCCT | 12 | 147 | 6.7 |  |  |  |
|  |  | TTGATGCTACAATGAAGGA---------GCTTTCCCCT | 9 | 80 | 3.7 |  |  |  |
|  |  | TTGATGCTACAATGAAGGAA-------AGCTTTACCCT | 7 | 76 | 3.5 |  |  |  |

| | | | | | | | | |
|---|---|---|---|---|---|---|---|---|
| NbCoeTnpBe-reRNA4$^{PDS}$ | 1 | TTGATGCTACAATGAAGCA----GCAAAGCTTTTCCCT | 4 | 42 | 9.4 | 17.6 | 29 | 18 |
|  |  | TTGATGCTACAATGAAGGAACTA----AGCTTTTCCCT | 4 | 35 | 7.8 |  |  |  |
| | | TTGATGCTACAATGAAGGAACTAGC-AAGCTTTTCCCT | 1 | 28 | 6.3 | | | |
|  |  | TTGATGCTACAATGAAGGAA--------GCTTTTCCCT | 8 | 25 | 5.6 |  |  |  |
|  | 2 | TTGATGAATGAAGGA----GCAAAGCTTTTCCCTTTCG | 4 | 98 | 9.6 | 16.1 |  |  |
|  |  | TTGATGAATGAAGGAA--------GCTTTTCCCTTTCG | 8 | 73 | 7.1 |  |  |  |
|  |  | TTGATGAATGAAGGAACTA----AGCTTTTCCCTGATA | 4 | 68 | 6.7 |  |  |  |
| | | TTGATGAATGAAGGAACTAGC-AAGCTTTTCCCTTTCG | 1 | 58 | 5.7 | | | |
|  | 3 | TTGATGCTACAATGAAGGA----GCGAAGCTTTTCCCT | 4 | 152 | 8.3 | 25.5 |  |  |
|  |  | TTGATGCTACAATGAAGGAA-------AGCTTTTCCCT | 7 | 145 | 7.9 |  |  |  |
|  |  | -------------------------------------- | 47 | 100 | 5.4 |  |  |  |
|  |  | TTGATGCTACAATGAAGGAA--------GCTTTTCCCT | 8 | 95 | 5.2 |  |  |  |
|  | 4 | TTGATGCTACAATGAAG------------CTTTTCCCT | 12 | 203 | 8.1 | 54.8 |  |  |
|  |  | TTGATGCTACAATGAAGGAA--------GCTTTTCCCT | 8 | 163 | 6.5 |  |  |  |
|  |  | TTGATGCTACAATGAAGGA----GCAAAGCTTTTCCCT | 4 | 151 | 6.0 |  |  |  |
|  |  | TTGATGCTACGATGAAGGAA-------AGCTTTTCCCT | 7 | 144 | 5.7 |  |  |  |
| NbCoeTnpBe-$^{HH}$reRNA4$^{PDS}$ | 1 | TTGATGCTACAATGAAGGAA--------GCTTTTCCCT | 8 | 90 | 6.9 | 32.7 | 20 | 8 |
|  |  | TTGATGCTACAATGAAGGA----GCAGAGCTTTTCCCT | 4 | 88 | 6.8 |  |  |  |
|  |  | TTGATGCTACAATGAAGGAA-------AGCTTTTCCCT | 7 | 77 | 5.9 |  |  |  |
|  |  | TTGATGCTACAATGAAG------------CTTTTCCCT | 12 | 61 | 4.7 |  |  |  |
|  | 2 | TTGATGCTACAATGAAGGA----GCAAAGCTTTTCCCT | 4 | 131 | 11.6 | 16.7 |  |  |
|  |  | TTGATGCTACAATGAAGGAACAA----AGCTTTTCCCT | 4 | 117 | 10.3 |  |  |  |
|  |  | TTGATGCTACAATGAAGGAA-------AGCTTTTCCCT | 7 | 56 | 4.9 |  |  |  |
| | | TTGATGCTACAATGAAGGAA---GCTAAGCCTTTCCCT | 3 | 53 | 4.7 | | | |
|  | 3 | TTGATGCTACAATGAAGGA----GCGAAGCTTTTCCCT | 4 | 161 | 11.8 | 16.0 |  |  |
|  |  | TTGATGCTACAATGAAGGAACAA----AGCTTGTCCCT | 4 | 109 | 8.0 |  |  |  |
|  |  | TTGATGCTACAATGAAGGAA-------AGCTTGTCCCT | 7 | 89 | 6.5 |  |  |  |
|  |  | TTGATGCTACAATGAAAGAA--------GCTTTTCCCT | 8 | 77 | 5.6 |  |  |  |
|  | 4 | TTGATGCTACAATGAAGGA----GCGAAGCTTTTCCCT | 4 | 125 | 12.2 | 14.8 |  |  |
|  |  | TTGATGCTACAATGAAGGAACAA----AGCTTTTCCCT | 4 | 92 | 9.0 |  |  |  |
|  |  | TTGATGCTACAATGAAGGAA-------AGCTTTTCCCT | 7 | 92 | 9.0 |  |  |  |
|  |  | TTGATGCTGCAATGAAGGAA--------GCTTTTCCCT | 8 | 51 | 5.0 |  |  |  |

*Bold red letters indicate TAM, and underlined letters denote reRNA target sequences

**Supplementary Table 10.** Efficiency of somatic editing at the *PDS* locus in infiltrated leaves targeted by *N. tabacum* codon optimized (NtCo) eTnpBc variant with different reRNAs

| Construct | Plant # | Most common indels | n_deleted | Reads | % Edited reads | Overall editing frequency | Average editing frequency | Standard deviation |
|---|---|---|---|---|---|---|---|---|
| | | TTGATTTTCCTGAAGCTCTTCCTGCGCCATTAAATGGT* | Wild-type | | | | | |
| NtCoeTnpBc-reRNA2$^{PDS}$ | 1 | TTGATTTTCCTGAAGCTCTTC-TGCGCCATTAAATGGT | 1 | 12 | 80.0 | 0.2 | 0 | 0 |
| | | TTGATTTTCCTGAAGCTCTTCCTGCGCCATTAA-TGGT | 1 | 2 | 13.3 | | | |
|  | 2 | TTGATTTTCCTGAAGCT------GCGCCATTAAATGGT | 6 | 8 | 22.9 | 0.2 |  |  |
| | | TTGATTTTCCTGTAGCTCTTCCTGCGCCATTAAATGGT | 1 | 5 | 14.3 | | | |
| | | TTGATTTTCCTGAAGCTCTTCCTGCGCCATTAA-TGGT | 1 | 4 | 11.4 | | | |
| | | TTGATTTTCCTGAAGCTCTTC-TGCGCCATTGAATGGT | 1 | 4 | 11.4 | | | |
| | 3 | TTGATTTTCCTGAAGCTCTTC-TGCGCCATTAAATGGT | 1 | 4 | 36.4 | 0.1 | | |
|  |  | TTGATTTTCCTGAAGCTCT------------AAATGGT | 12 | 3 | 27.3 |  |  |  |
|  |  | TTGATTTTCCTGAAGCT------GCGCCATTAAATGGT | 6 | 1 | 9.1 |  |  |  |
| | | TTGATTTTCCTGAAGCTCTTCCTGCGCCATTAAATG-T | 1 | 1 | 9.1 | | | |
|  | 4 | TTGATTTTCCTGAAGCTCTTCC------ATTAAATGGT | 6 | 13 | 24.5 | 0.4 |  |  |
|  |  | TTGATTTTCCTGAAGCTCTTCCT-------TAAATGGT | 7 | 9 | 17.0 |  |  |  |
| | | TTGATTTTCCTGAAGCTCTTCCTGCGCCATTAA-TGGT | 1 | 9 | 17.0 | | | |
| | | TTGATTTTCCTGAAGCTCTTC--GCGCCATTAAATGGT | 2 | 6 | 11.3 | | | |
| NtCoeTnpBc-$^{HH}$reRNA2$^{PDS}$ | 1 | TTGATTTTCCTGAAGCTCTTC----GCCATTAAATGGT | 4 | 9 | 26.5 | 0.4 | 0 | 0 |
| | | TTGATTTTCCTGAAGCTCT-CCTGCGCCATTAAATGGT | 1 | 7 | 20.6 | | | |
|  |  | TTGATTTTCCTGAAGCTCT------------AAATGGT | 12 | 6 | 17.6 |  |  |  |
| | | TTGATTTTCCTGAAGCTCTTCCTGCGCCATTAA-TGGT | 1 | 5 | 14.7 | | | |
| | 2 | TTGATTTTCCTGAAGCTCTTC-TGCGCCATTAAATGGT | 1 | 18 | 40.0 | 0.4 | | |
| | | TTGATTTTCCTGAAGCTCTTCCTGCGCCATTAA-TGGT | 1 | 10 | 22.2 | | | |
| | | TTGATTTTCCTGGAGCTCTTCC---GCCATTAAATGGT | 3 | 3 | 6.7 | | | |
|  |  | TTGATTTTCCTGAAGCTCTTCC------ATTAAATGGT | 6 | 2 | 4.4 |  |  |  |
|  | 3 | TTGATTTTCCTGAAGCTCTTCC------ATTAAATGGT | 6 | 8 | 33.3 | 0.5 |  |  |
| | | TTGATTTTCCTGAAGCTCTTC--GCGCCATTAAATGGT | 2 | 7 | 29.2 | | | |
| | | TTGATTTTCCTGAAGCTCTTC-TGCGCCATTAAATGGT | 1 | 5 | 20.8 | | | |
| | | TTGATTTTCCTGAAGCTCTTCCTGCGCCATTAA-TGGT | 1 | 3 | 12.5 | | | |
| | 4 | TTGATTTTCCTGAAGCTCTT---GCGCCATTAAATGGT | 3 | 9 | 20.9 | 0.6 | | |
|  |  | TTGATTTTCCTGAAGCTCTTC----GCCATTAAATGGT | 4 | 6 | 14.0 |  |  |  |
| | | TTGATTTTCCTGAAGCTCTTC-TGCGCCATTAAATGGT | 1 | 5 | 11.6 | | | |
|  |  | TTGATTTTCCTGAAGCTC-------------------- | 21 | 4 | 9.3 |  |  |  |
| | | TTGATTGCTTTGAACAGATTTCTTCAGGTTAGAATCCC* | Wild-type | | | | | |
| NtCoeTnpBc-reRNA3$^{PDS}$ | 1 | TTGATTGCTTTGAACAGATTTC---AGGTTAGAATCCT | 3 | 182 | 34.1 | 1.8 | 3 | 1 |
|  |  | TTGATTGCTTTGAACAGATT-----AGGTTAGAATCCT | 5 | 30 | 5.6 |  |  |  |
|  |  | TTGATTGCTTTGAACAGATTT---------AGGATCCT | 9 | 27 | 5.1 |  |  |  |
|  |  | TTGATTGCTTTGAACAGATAT-----------AATCCC | 11 | 16 | 3.0 |  |  |  |
| | 2 | TTGATTGCTTTGAACAGATTTC---AGGTTAGAATCCT | 3 | 54 | 15.0 | 2.5 | | |
|  |  | TTGATTGCTTTGAACAGAT------AGGTTAGAATCCT | 6 | 26 | 7.2 |  |  |  |
|  |  | TTGATTGCTTTGAACAGATT----CAGGTTAGAATCCC | 4 | 19 | 5.3 |  |  |  |
|  |  | TTGATTGCTTTGAATAGAT-----CAGGTTAGAATCCC | 5 | 18 | 5.0 |  |  |  |
| | 3 | TTGATTGCTTTGAACAGATTTC---AGGTTAGAATCCC | 3 | 63 | 16.6 | 3.3 | | |
|  |  | TTGATTGCTTTGAACAGA--------GGTTAGAATCCT | 8 | 22 | 5.8 |  |  |  |
|  |  | TTGATTGCTTTGAACAGATT-----AGGTTAGAATCTT | 5 | 20 | 5.3 |  |  |  |
|  |  | TTGATTGCTTTGAACAGAAGA----AGTTAAGAATCCT | 4 | 19 | 5.0 |  |  |  |
| | 4 | TTGATTGCTTTGAACAGATTTC---AGGTTAGAATCCC | 3 | 63 | 16.4 | 2.8 | | |
|  |  | TTGATTGCTTTGAACAG----------GTTAGAATCCC | 10 | 24 | 6.3 |  |  |  |
|  |  | TTGATTGCTTTGAACAGATC----------AGAATCCT | 10 | 22 | 5.7 |  |  |  |
|  |  | TTGATTGCTTTGAACAGATGT----------------- | 28 | 21 | 5.5 |  |  |  |
| NtCoeTnpBc-$^{HH}$reRNA3$^{PDS}$ | 1 | TTGATTGCTTTGAACAGATTTC---AGGTTAGAATCCT | 3 | 167 | 24.3 | 1.7 | 3 | 1 |
|  |  | TTGATTGCTTTGAACAGATT-----AGGTCAGAATCCC | 5 | 37 | 5.4 |  |  |  |
|  |  | TTGATTGCTTTGAACAGAT-----CAGGTTAGAATCCT | 5 | 35 | 5.1 |  |  |  |
|  |  | TTGATTGCTTTGAACAGGTAT-----------AATCCC | 11 | 24 | 3.5 |  |  |  |
| | 2 | TTGATTGCTTTGAACAGATTTC---AGGTTAGAATCCT | 3 | 69 | 15.0 | 2.9 | | |
|  |  | TTGATTGCTGTGAACAGAT-----CAGGTTAGAATCCT | 5 | 30 | 6.5 |  |  |  |
|  |  | TTGATTGCTTTGAACAG----------GTTAGAATCCC | 10 | 28 | 6.1 |  |  |  |
| | | TTGATTGCTTTGAACAGATTTCT---GGTTAGAATCCC | 3 | 24 | 5.2 | | | |
| | 3 | TTGATTGCTTTGAACAGATTTC---AGGTTAGAATCCC | 3 | 398 | 17.8 | 3.3 | | |
|  |  | TTGATTGCTTTGAACAGG--------GGTTAGAATCCC | 8 | 106 | 4.7 |  |  |  |
|  |  | TTGATTGCTTCGAACAGATT-----AGGTTAGAATCCT | 5 | 103 | 4.6 |  |  |  |
|  |  | TTGATTGCTTTGAACAGATT----------AAAATCCC | 10 | 79 | 3.5 |  |  |  |
| | 4 | TTGATTGCTTTGAACAGATTTC---AGGTTAGAATCCC | 3 | 48 | 22.5 | 2.9 | | |
|  |  | TTGATTGCTTTGAACAGATTT---------AGAATCCT | 9 | 16 | 7.5 |  |  |  |
|  |  | TTGATTGCTTTGAACAGATTT----AGGTTAGAATCCT | 4 | 14 | 6.6 |  |  |  |
|  |  | TTGATTGCTTTGAACAGA--------GGTTAGAATCCC | 8 | 13 | 6.1 |  |  |  |

| | | | | | | | |
|---|---|---|---|---|---|---|---|
| | | TTGATGCTACAATGAAGGAACTAGCGAAGCTTTTCCCT* | Wild-type | | | | |
| NtCoeTnpBc-reRNA4$^{PDS}$ | 1 | TTGATGCTACAATGAAGGA----GCGAAGCTTTTCCCT | 4 | 175 | 12.7 | 9.4 | 7 | 2 |
|  |  | TTGATGCTACAATGAAGGAAC----GAAGCTTTTCCCT | 4 | 93 | 6.7 |  |  |  |
|  |  | TTGATGCTACAATGAAGGAA-------AGCTTTTCCCT | 7 | 65 | 4.7 |  |  |  |
|  |  | TTGATGCTACAATGAA-----------AGCTTTTCCCT | 11 | 58 | 4.2 |  |  |  |
|  | 2 | TTGATGCTACAATGAAGCAA-------AGCTTTTCCCT | 7 | 204 | 8.2 | 5.2 |  |  |
|  |  | TTGATGCTACAATGAAGGA----GCGAAGCTTTTCCCT | 4 | 193 | 7.8 |  |  |  |
| | | TTGATGCTACAATGAATGAACTAGAGAATC-TTTCCCT | 1 | 90 | 3.6 | | | |
|  |  | TTGATGCTACAATGAAT------------CTTTTCCCT | 12 | 68 | 2.7 |  |  |  |
|  | 3 | TTGATGCTACAATGAAGGA----GCGAAGCTTTTCCCT | 4 | 154 | 9.5 | 5.3 |  |  |
| | | TTGATGCTACAATGAATGAACTAGCGAAGC-TTTCCCT | 1 | 129 | 7.9 | | | |
|  |  | TTGATGCTACAATGAAG------------CTTTTCCCT | 12 | 128 | 7.9 |  |  |  |
|  |  | TTGATGCTACAATGAAGGAA-------AGCTTTTCCCT | 7 | 85 | 5.2 |  |  |  |
|  | 4 | TTGATGCTACAATGAAGGA----GCGAAGCTTTTCCCT | 4 | 105 | 9.7 | 8.9 |  |  |
|  |  | TTGATGCTACAATGAAGGAA-------AGCTTTTCCCT | 7 | 71 | 6.5 |  |  |  |
| | | TTGATGCTACAATGAAGGAACTAGCGATGC-TTTCCCT | 1 | 55 | 5.1 | | | |
|  |  | TTGATGCTACAATGAAG------------CTTTTCCCT | 12 | 50 | 4.6 |  |  |  |
| NtCoeTnpBc-$^{HH}$reRNA4$^{PDS}$ | 1 | TTGATGCTACAATGAAGGA----GCGAAGCTTTTCCCT | 4 | 149 | 8.6 | 19.1 | 15 | 3 |
|  |  | TTGATGCTACAATGAAGGAA--------GCTTTTCCCT | 8 | 127 | 7.3 |  |  |  |
|  |  | TTGATGCTACAATGAAGGAA-------AGCTTTTCCCT | 7 | 96 | 5.5 |  |  |  |
|  |  | TTGATGCTACAATGAAG------------CTTTTCCCT | 12 | 87 | 5.0 |  |  |  |
|  | 2 | TTGATGCTACAATGAAGGA----GCAAAGCTTTTCCCT | 4 | 123 | 9.7 | 15.3 |  |  |
|  |  | TTGATGCTACAATGACG------------CTTTTCCCT | 12 | 51 | 4.0 |  |  |  |
|  |  | ---------------------------AGCTTTTCCCT | 27 | 43 | 3.4 |  |  |  |
|  |  | TTGATGCTACAATGAAGATA-------AGCTTTTCCCT | 7 | 42 | 3.3 |  |  |  |
|  | 3 | TTGATGCTACAATGAAGGA----GCGAAGCTTTTCCCT | 4 | 133 | 12.1 | 14.5 |  |  |
|  |  | TTGATGCTACAATGAAGGAA-------AGCTTTTCCCT | 7 | 74 | 6.7 |  |  |  |
|  |  | TTGATGCTACAATGAAG------------CTTTTCCCT | 12 | 52 | 4.7 |  |  |  |
|  |  | TTGATGCTACAATGAAGGAA--------GCTTTTCCCT | 8 | 42 | 3.8 |  |  |  |
|  | 4 | TTGATGCTACAATGAAGGA----GCGAAGCTTTTCCCT | 4 | 145 | 14.7 | 12.3 |  |  |
|  |  | TTGATGCTACAATGAAGGAAC----GAAGCTTTTCCCT | 4 | 44 | 4.5 |  |  |  |
|  |  | TTGATGCTACAATGAAGGAACAA----AGCTTTTCCCT | 4 | 38 | 3.8 |  |  |  |
|  |  | TTGATGCTACAATGAAGGAA--------GCTTGTCCCT | 8 | 34 | 3.4 |  |  |  |

*Bold red letters indicate TAM, and underlined letters denote reRNA target sequences

**Supplementary Table 11.** Efficiency of somatic editing at the *ChlH* locus in infiltrated leaves targeted by the eTnpBc variant with reRNA[ChlH]

| Plant # | Tissue sample # | Most common indels | n_deleted | Reads | % Edited reads | Overall editing frequency | Average editing frequency | Standard deviation |
|---|---|---|---|---|---|---|---|---|
| | | TTGTCAACTGCTCTGGGGTGTTCAGAGATCTCTTC**ATCAA*** | Wild-type | | | | | |
| 1 | 1 | TTGTCAACTGCTC------------AGATCTCTTCATCAA | 12 | 1354 | 12.0 | 96.9 | 90 | 12 |
|  |  | TTGTCAACTGCTC--------------ATCTCTTCATCAA | 14 | 533 | 4.7 |  |  |  |
|  |  | ----------------------CAGAGATCTCTTCATCAA | 54 | 465 | 4.1 |  |  |  |
|  |  | ------------------------------TCTTCATCAA | 61 | 270 | 2.4 |  |  |  |
|  | 2 | TTGTCAACTGC--------------AGATCTCTTCATCAA | 14 | 482 | 4.0 | 94.6 |  |  |
|  |  | TTGTCAACTGCTCT-----------AGATCTCTTCATCAA | 11 | 465 | 3.9 |  |  |  |
|  |  | TTATCAACTGCTC------------AGATCTCTTCATCAA | 12 | 454 | 3.8 |  |  |  |
|  |  | TTGTCAACTGTTC-------------GATCTCTTCATCAA | 13 | 417 | 3.5 |  |  |  |
| 2 | 1 | TTGTCAACTGCTC------------AGATCTCTTCATCAA | 12 | 207 | 5.8 | 95.0 |  |  |
|  |  | ---------------------TCAGAGATCTCTTCATCAA | 38 | 121 | 3.4 |  |  |  |
|  |  | ----------------------CAGAGATCTCTTCATCAA | 35 | 110 | 3.1 |  |  |  |
|  |  | TTGTCAACTGCGC----------AGAGATCTCTTCATCAA | 10 | 104 | 2.9 |  |  |  |
| | 2 | TTGTCAACTGCTCTGGG-TGTTCAGAGATCTCTTCATCAA | 1 | 172 | 3.3 | 57.9 | | |
|  |  | TTGTCAACTGC-------------GAGATCTCTTCATCAA | 13 | 170 | 3.3 |  |  |  |
|  |  | ------------------------------TCTTCATCAA | 51 | 149 | 2.9 |  |  |  |
|  |  | TTGTCAACTGCTC----------AGAGATCTCTTCATCAA | 10 | 141 | 2.7 |  |  |  |
| 3 | 1 | TTGTCAACTGCTCTG---------------TCTTCATCAA | 36 | 335 | 8.5 | 96.7 |  |  |
|  |  | TTGTCAACTGCTC--------------ATCTCTTCATCAA | 14 | 193 | 4.9 |  |  |  |
|  |  | TTGTCAACTGCTC------------AGATCTCTTCATCAA | 12 | 147 | 3.7 |  |  |  |
|  |  | ----------------------------TCTCTTCATCAA | 29 | 105 | 2.7 |  |  |  |
|  | 2 | TTGTCAACTGC-------------GAGATCTCTTCATCAA | 13 | 1095 | 6.8 | 89.8 |  |  |
|  |  | ------------------------------TCTTCATCAA | 54 | 600 | 3.7 |  |  |  |
|  |  | TTGTCAACCGCT---------------ATCTCTTCATCAA | 15 | 598 | 3.7 |  |  |  |
|  |  | ---------------------TCAGAGATCTCTTCATCAA | 46 | 365 | 2.3 |  |  |  |
| 4 | 1 | -------------------------AGATCTCTTCATCAA | 27 | 1171 | 13.5 | 97.0 |  |  |
|  |  | TTGTGAAG--------------CAGTGATCTCTTCATCAA | 14 | 324 | 3.7 |  |  |  |
|  |  | TTGTCAACTGC------------------CTCTTCATCAA | 18 | 292 | 3.4 |  |  |  |
|  |  | ----------------------CAGAGATCTCTTCATCAA | 59 | 250 | 2.9 |  |  |  |
|  | 2 | ----------------------CAGAGATCTCTTCATCAA | 62 | 480 | 3.9 | 97.0 |  |  |
|  |  | ---------------------TCAAAGATCTCTTCATCAA | 34 | 338 | 2.7 |  |  |  |
|  |  | ------------------------------TCTTCATCAA | 47 | 334 | 2.7 |  |  |  |
|  |  | TTGTCAACTGC----------------ATCTCTTCATCAA | 16 | 306 | 2.5 |  |  |  |
| 5 | 1 | TTGTCAACTGC--------------AGATCTCTTCATCAA | 14 | 17914 | 38.6 | 79.6 |  |  |
|  |  | TTGTCAACTGCTCT-------TCAGAGATCTCTTCATCAA | 7 | 16451 | 35.5 |  |  |  |
|  |  | TTGTCAACTGCTCTG-----TTCAGAGATCTCTTCATCAA | 5 | 10916 | 23.5 |  |  |  |
|  |  | ------------------------GAGATCTCTTCATCAA | 37 | 395 | 0.9 |  |  |  |
|  | 2 | TTGTCAACTGC--------------AGATCTCTTCATCAA | 14 | 19712 | 35.2 | 79.5 |  |  |
|  |  | TTGTCAACTGCTCT-------TCAGAGATCTCTTCATCAA | 7 | 17674 | 31.5 |  |  |  |
|  |  | TTGTCAACTGCTCTG-----TTCAGAGATCTCTTCATCAA | 5 | 16004 | 28.6 |  |  |  |
|  |  | TTGTCAACTG--------------------TCTTCATCAA | 66 | 856 | 1.5 |  |  |  |
| 6 | 1 | TTGTCAACTGCTC-------------GATCTCTTCATCAA | 13 | 4438 | 9.7 | 97.2 |  |  |
|  |  | -------------------------AGATCTCTTCATCAA | 27 | 4381 | 9.6 |  |  |  |
|  |  | TT---------------------------CTCTTCATCAA | 27 | 3354 | 7.3 |  |  |  |
|  |  | ------------------------------TCTTCATCAA | 65 | 3041 | 6.6 |  |  |  |
|  | 2 | TTGTCAACTG-----------------ATCTCTTCATCAA | 17 | 6950 | 13.9 | 97.1 |  |  |
|  |  | TTGTCAACTGC-------------GAGATCTCTTCATCAA | 13 | 5003 | 10.0 |  |  |  |
|  |  | TTCTCAACTGCTCT-----------AGATCTCTTCATCAA | 11 | 4385 | 8.8 |  |  |  |
|  |  | -------------------------AGATCTCTTCATCAA | 34 | 2979 | 6.0 |  |  |  |

*Bold red letters indicate TAM, and underlined letters denote reRNA target sequences

**Supplementary Table 12.** Efficiency of somatic editing at the *PDS* locus in infiltrated leaves targeted by ISYmu1 with reRNA4$^{PDS}$

| Construct | Plant # | Most common indels | n_deleted | Reads | % Edited reads | Overall editing frequency | Average editing frequency | Standard deviation |
|---|---|---|---|---|---|---|---|---|
| | | **TTGAT**GCTACAATGAAGGAACTAGCGAAGCTTTTCCCT* | Wild-type | | | | | |
| NtCoISYmu1-reRNA4$^{PDS}$ | 1 | TTGATGCTACAATGAAGGAA-------AGCTTTTCCCT | 7 | 696 | 7.9 | 32.6 | 33 | 2 |
|  |  | TTGATGCTACAAT-------------AAGCTTTTCCCT | 13 | 470 | 5.3 |  |  |  |
|  |  | TTGATGCTACAATGAAG------------CTTTTCCCT | 12 | 444 | 5.0 |  |  |  |
|  |  | TTGATGCTACAATGAAGGAACAA----AGCTTTTCCCT | 4 | 437 | 5.0 |  |  |  |
|  | 2 | TTGATGCTACAATGAAGGAA-------AGCTTTTCCCT | 7 | 470 | 5.4 | 34.8 |  |  |
|  |  | TTGATGCTACAATGAAG------------CTTTTCCCT | 12 | 456 | 5.2 |  |  |  |
|  |  | TTGATGCTACAATGAAGGA----GTGAAGCCTTTCCCT | 4 | 454 | 5.2 |  |  |  |
|  |  | TTGATGCTACAATGAAGGAA--------GCTTTTCCCT | 8 | 398 | 4.5 |  |  |  |
|  | 3 | TTGATGCTACAATGAAGGAA-------AGCTTTTCCCT | 7 | 635 | 7.8 | 31.7 |  |  |
|  |  | TTGATGCTACAATGAAG------------CTTTTCCCT | 12 | 618 | 7.6 |  |  |  |
|  |  | TTGATGCTACAATGAAGGAACAA----AGCTTTTCCCT | 4 | 455 | 5.6 |  |  |  |
|  |  | TTGATGCTACAATGAAGGAA------AAGCTTTTCCCT | 6 | 371 | 4.6 |  |  |  |
|  | 4 | TTGATGCTACAATGAAGGAA--------GCTTTTCCCT | 8 | 452 | 5.9 | 31.4 |  |  |
|  |  | TTGATGCTACAATGAAGGAA-------AGCTTTTCCCT | 7 | 400 | 5.2 |  |  |  |
|  |  | TTGATGCTACAATGAAGGA----GCGAAGCTTTTCCCT | 4 | 386 | 5.0 |  |  |  |
|  |  | TTGATGCTACAAT-------------AAGCTATTCCCT | 13 | 372 | 4.9 |  |  |  |
| HsCoISYmu1-reRNA4$^{PDS}$ | 1 | TTGATGCTACAATGAAGGAACAA----AGCTTTTCCCT | 4 | 697 | 7.8 | 26.7 | 24 | 3 |
|  |  | TTGATGCTACAATGAAGGA----GCGAAGCTTTTCCCT | 4 | 634 | 7.1 |  |  |  |
|  |  | TTGATGCTACAATGAAGGAA--------GCTTTTCCCT | 8 | 554 | 6.2 |  |  |  |
|  |  | TTGATGCTACAAT-------------AAGCTTTTCCCT | 13 | 399 | 4.4 |  |  |  |
|  | 2 | TTGATGCTACAATGAAGGA----GCAAAGCTTTTCCCT | 4 | 647 | 13.9 | 19.7 |  |  |
|  |  | TTGATGCTACAATGAAGGAACTA----AGCTTTTCCCT | 4 | 356 | 7.6 |  |  |  |
|  |  | TTGATGCTACAAA-------------AAGCTTTTCCCT | 13 | 339 | 7.3 |  |  |  |
|  |  | TTGATGCTACAATGAA-----------AGCTTTTCCCT | 11 | 279 | 6.0 |  |  |  |
|  | 3 | TTGATGCTACAATGAAGGAACAA----AGCTTTTCCCT | 4 | 472 | 7.6 | 23.1 |  |  |
|  |  | TTGATGCTACAATGAAGGAA--------GCTTTTCCCT | 8 | 429 | 6.9 |  |  |  |
|  |  | TTGATGCTACAATGAATCAA-------AGCTTTTCCCT | 7 | 408 | 6.6 |  |  |  |
|  |  | TTGATGCTACAATGAAG------------CTTTTCACT | 12 | 394 | 6.3 |  |  |  |
|  | 4 | TTGATGCTACAATGAAGGAACAA----AGCTTTTCCCT | 4 | 215 | 6.9 | 26.4 |  |  |
|  |  | TTGATGCTACAATGAAGGA----GCGAAGCTTTTCCCT | 4 | 177 | 5.7 |  |  |  |
|  |  | TTGATGCTACAATGAAGGAA-------AGCTTTTCCCT | 7 | 174 | 5.6 |  |  |  |
|  |  | TTGATGCTACAAT-------------AAGCTTTTCCCT | 13 | 167 | 5.4 |  |  |  |

*Bold red letters indicate TAM, and underlined letters denote reRNA target sequences

**Supplementary Table 13.** Percentage of progeny seedlings from parental plants infected with TRV2 expressing TnpB and TnpB variants with reRNA4$^{PDS}$ showing the white photobleaching phenotype

| Construct | Progeny seeeds from | # of green seedlings | # of white seedlings | Total # of seedlings | % white seedlings | Average % of white seedlings | Standard Deviation |
|---|---|---|---|---|---|---|---|
| TnpB-reRNA4$^{PDS}$ | Pods from the top 1/3rd of the plant | 155 | 0 | 155 | 0 | 0 | 0 |
| | | 144 | 0 | 144 | 0 | | |
| | | 137 | 0 | 137 | 0 | | |
| | | 85 | 0 | 85 | 0 | | |
| | | 65 | 0 | 65 | 0 | | |
| TnpB-$^{HH}$reRNA4$^{PDS}$ | Pods from the top 1/3rd of the plant | 96 | 0 | 96 | 0 | 0 | 0 |
| | | 310 | 0 | 310 | 0 | | |
| | | 95 | 0 | 95 | 0 | | |
| | | 53 | 0 | 53 | 0 | | |
| | | 90 | 0 | 90 | 0 | | |
| eTnpBc-reRNA4$^{PDS}$ | Pods with phtobleaching phenotype | 14 | 57 | 71 | 80 | 89 | 9 |
| | | 0 | 188 | 188 | 100 | | |
| | | 2 | 28 | 30 | 93 | | |
| | | 14 | 56 | 70 | 80 | | |
| | | 2 | 27 | 29 | 93 | | |
| | Pods from the top 1/3rd of the plant | 12 | 15 | 27 | 56 | 52 | 14 |
| | | 20 | 30 | 50 | 60 | | |
| | | 12 | 27 | 39 | 69 | | |
| | | 15 | 10 | 25 | 40 | | |
| | | 26 | 14 | 40 | 35 | | |
| eTnpBc-$^{HH}$reRNA4$^{PDS}$ | Pods with phtobleaching phenotype | 3 | 57 | 60 | 95 | 89 | 8 |
| | | 0 | 77 | 77 | 100 | | |
| | | 33 | 138 | 171 | 81 | | |
| | | 4 | 21 | 25 | 84 | | |
| | | 20 | 100 | 120 | 83 | | |
| | Pods from the top 1/3rd of the plant | 12 | 16 | 28 | 57 | 53 | 20 |
| | | 14 | 55 | 69 | 80 | | |
| | | 28 | 14 | 42 | 33 | | |
| | | 18 | 9 | 27 | 33 | | |
| | | 20 | 32 | 52 | 62 | | |
| eTnpBe-reRNA4$^{PDS}$ | Pods from the top 1/3rd of the plant | 55 | 0 | 55 | 0 | 0 | 0 |
| | | 76 | 0 | 76 | 0 | | |
| | | 64 | 0 | 64 | 0 | | |
| | | 49 | 0 | 49 | 0 | | |
| | | 85 | 0 | 85 | 0 | | |
| eTnpBe-$^{HH}$reRNA4$^{PDS}$ | Pods from the top 1/3rd of the plant | 212 | 0 | 212 | 0 | 0 | 0 |
| | | 99 | 0 | 99 | 0 | | |
| | | 136 | 0 | 136 | 0 | | |
| | | 89 | 0 | 89 | 0 | | |
| | | 72 | 0 | 72 | 0 | | |

**Supplementary Table 14.** Indel types and frequencies in *PDS* of progenies from parental plants infected with TRV2 expressing eTnpBc variant and reRNA4$^{PDS}$

| Construct | Parent # | Plant # | Most common indels | n_deleted | Reads | % Edited reads | Overall editing frequency | Average editing frequency | Standard deviation |
|---|---|---|---|---|---|---|---|---|---|
| | | | TTGATGCTACAATGAAGGAACTAGCGAAGCTTTTCCCT* | Wild-type | | | | | |
| eTnpBc-reRNA4$^{PDS}$ | 1 | 1 | TTGATGCTACAATGAAGGAAG--------------CCT | 14 | 1719 | 25.5 | 97.2 | 96 | 2 |
|  |  |  | TTGATGCTACAATGAAGGA----GCGAAGCTCTTCCCT | 4 | 1624 | 24.1 |  |  |  |
|  |  |  | TTGATGCTACTATGAAGGAACAA----AGCTTTTCCCT | 4 | 1582 | 23.5 |  |  |  |
|  |  |  | TTGATGCTACAATGAAG------------CTTTTCCCT | 12 | 1563 | 23.2 |  |  |  |
|  |  | 2 | TTGATGCTACAATGAAGGA----GCGAAGCTTTTCCCT | 4 | 2560 | 46.7 | 96.1 |  |  |
|  |  |  | TTGATGCTACAATGAAG------------CTTTTCCCT | 12 | 1443 | 26.4 |  |  |  |
|  |  |  | TTGATGCTACAATGAAGGATCAA----AGCTTTTCCCT | 4 | 1223 | 22.3 |  |  |  |
|  |  |  | TTGATGCTACAATGAAGGAAG--------------CCT | 14 | 26 | 0.5 |  |  |  |
|  |  | 3 | TTGATGCTACAATGAAGGA----GCGAAGCTTTTCCCT | 4 | 1988 | 45.6 | 92.7 |  |  |
|  |  |  | TTGATGCTACAATGAAGGAACAA----AGCTTTTCCCT | 4 | 1052 | 24.1 |  |  |  |
|  |  |  | TTGATGCTACAATGAAG------------CTTTTCACT | 12 | 1010 | 23.2 |  |  |  |
|  |  |  | TTGATGCTACAATGAAGGAAG--------------CCT | 14 | 25 | 0.6 |  |  |  |
|  |  | 4 | TTGATGCTACAATGAAGGAAG--------------CCT | 14 | 7254 | 52.0 | 98.2 |  |  |
|  |  |  | TTGATGCTACAATGAAGGAACAA----AGCTTTTCCCT | 4 | 6228 | 44.6 |  |  |  |
|  |  |  | TTGATGCTACAATGAAGGA----GCGAAGCTTTTCCCT | 4 | 97 | 0.7 |  |  |  |
|  |  |  | TTGATGCTACGATGAAG------------CTTTTCCCT | 12 | 69 | 0.5 |  |  |  |
|  |  | 5 | TTGATGCTACAATGAAG------------CTTTTCCCT | 12 | 4413 | 24.9 | 96.5 |  |  |
|  |  |  | TTGATGCTACAATGAAGGAAG--------------CCT | 14 | 4376 | 24.7 |  |  |  |
|  |  |  | TTGATGCTACAATGAAGGAACAA----AGCTTTTCCCT | 4 | 4004 | 22.6 |  |  |  |
|  |  |  | TTGATGCTACAATGAAGGA----GCGAAGCTTTTCCCT | 4 | 3957 | 22.3 |  |  |  |
|  |  | 6 | TTGATGCTACAATGAAGGAAG--------------CCT | 14 | 6478 | 26.1 | 95.4 |  |  |
|  |  |  | TTGATGCTACAATGAAG------------CTTTTCCCT | 12 | 5975 | 24.0 |  |  |  |
|  |  |  | TTGATGCTACAATGAAGGA----GCGAAGCTTTTCCCT | 4 | 5830 | 23.4 |  |  |  |
|  |  |  | TTGATGCTACAATGAAGGAACAA----AGCTTTTCCCT | 4 | 5569 | 22.4 |  |  |  |
|  |  | 7 | TTGATGCTACAATGAAGGAAG--------------CCT | 14 | 3154 | 54.5 | 97.8 |  |  |
|  |  |  | TTGATGCTACAATGAAGGAACAA----AGCTTTTCCCT | 4 | 2363 | 40.9 |  |  |  |
|  |  |  | TTGATGCTACAATGAAT------------CTTTTCCCT | 12 | 39 | 0.7 |  |  |  |
|  |  |  | TTGATGCTACAATGAAGGA----GCGAAGCTTTTCCCT | 4 | 38 | 0.7 |  |  |  |
|  |  | 8 | TTGATGCTACAATGAAGGAACAA----AGCTTTTCCCT | 4 | 13257 | 46.0 | 97.6 |  |  |
|  |  |  | TTGATGCTACACTGGAGGAAG--------------CCT | 14 | 7699 | 26.7 |  |  |  |
|  |  |  | TTGATGCTACAATGAAGGA----GCGAAGCTTTTCCCT | 4 | 6969 | 24.2 |  |  |  |
|  |  |  | TTGATGCTACAATGAAG------------CTTTTCCCT | 12 | 182 | 0.6 |  |  |  |
|  | 2 | 1 | -------------GCGAAGCTTTTCCCT | 39 | 6038 | 25.2 | 94.6 | 90 | 9 |
|  |  |  | TTGATGCTACAATGAAGGA----GCAAAGCTTTTCCCT | 4 | 4932 | 20.6 |  |  |  |
|  |  |  | TTGATGCTACAATGAAGGAA-------AGCTTTTCCCT | 7 | 4801 | 20.1 |  |  |  |
|  |  |  | TTGATGCTACAATGAAGGAACTA----AGCTTTTCCCT | 4 | 4713 | 19.7 |  |  |  |
|  |  | 2 | TTGATGCTACAATGAAGGAA-------AGCTTTTCCCT | 7 | 7304 | 36.0 | 70.9 |  |  |
|  |  |  | TTGATGCTACGAT------------------------- | 60 | 6111 | 30.1 |  |  |  |
|  |  |  | TTGATGCTACAATGTAGGAAC-----AAGCTTTTCCCT | 5 | 3495 | 17.2 |  |  |  |
|  |  |  | ---------------------------AGCTTTTCCCT | 44 | 272 | 1.3 |  |  |  |
|  |  | 3 | -----------------------GCGAAGCTTTTCCCT | 39 | 6107 | 26.5 | 90.7 |  |  |
|  |  |  | ----------------------------GCTTTTCCCT | 28 | 5752 | 25.0 |  |  |  |
|  |  |  | TTGATGCTACAATGAAGGAACTA----AGCTTTTCCCT | 4 | 4925 | 21.4 |  |  |  |
|  |  |  | TTGATGCTACAATGAAGGAA-------AGCTTTTCCCT | 7 | 4576 | 19.9 |  |  |  |
|  |  | 4 | ---------------------------AGCTTTTCCCT | 18 | 5854 | 25.2 | 96.3 |  |  |
|  |  |  | TTGATGCTACAATGAAGGA----GCAAAGCTTTTCCCT | 4 | 5466 | 23.6 |  |  |  |
|  |  |  | TTGATGCTACAATGAAGGAA--------GCTTTTCCCT | 8 | 5359 | 23.1 |  |  |  |
|  |  |  | TTGATGCTACAATGAAGGAA-------AGCTTTTCCCT | 7 | 5348 | 23.0 |  |  |  |
|  |  | 5 | TTGATGCTACAATGAAGGAACAA----AGCTTTTCCCT | 4 | 3151 | 12.5 | 91.6 |  |  |
|  |  |  | TTGATGCTACAATGAAGGAAG--------------CCT | 14 | 2811 | 11.2 |  |  |  |
|  |  |  | TTGATGCTACAATGAAG------------CTTTTTCCT | 12 | 1592 | 6.3 |  |  |  |
|  |  |  | TTGATGCTACAATGA-------------GCTTTTCCCT | 13 | 1284 | 5.1 |  |  |  |
|  |  | 6 | TTGATGCTACAATGAAGGA----GCGAAGCTTTTCCCT | 4 | 5013 | 19.9 | 93.8 |  |  |
|  |  |  | TTGATGCTACAATGAAGGAAC-------GCTTTTCCCT | 7 | 4290 | 17.0 |  |  |  |
|  |  |  | TTGATGCTACAATGAAGGAAC------AGCTTTTCCCT | 6 | 3809 | 15.1 |  |  |  |
|  |  |  | TTGATGCTACAATGA----------GAAGCTTTTCCCT | 10 | 3611 | 14.3 |  |  |  |
| eTnpBc-$^{HH}$reRNA4$^{PDS}$ | 1 | 1 | ---------------------------AGCTTTTCCCT | 86 | 10107 | 51.2 | 92.7 | 92 | 6 |
|  |  |  | TTGATGCTACAATGAAGGAA--------GCT-----CT | 13 | 4404 | 22.3 |  |  |  |
| | | | TTGATGCTACAATGAAGGAACTAGC-AAGCTTTTCCCT | 1 | 3965 | 20.1 | | | |
|  |  |  | TTGATGCTACAATGAAGGAA--------GCCTTTCCCT | 8 | 82 | 0.4 |  |  |  |
|  |  | 2 | ---------------------------AGCTTTTCCCT | 86 | 5567 | 28.8 | 86.1 |  |  |
|  |  |  | TTGATGCTACAATGAAGGAA--------GCT-----CT | 13 | 4533 | 23.5 |  |  |  |
| | | | TTGATGCTACAATGAAGGAACTAGC-AAGCTTTTCCCT | 1 | 4336 | 22.5 | | | |
|  |  |  | TTGATGCTACAATGAAGGAAC----GAAGCTTTTCCCT | 4 | 3948 | 20.4 |  |  |  |
|  |  | 3 | ---------------------------AGCTTTTCCCT | 86 | 7165 | 38.4 | 83.3 |  |  |
|  |  |  | TTGATGCTACAATGAAGGAA--------GCT-----CT | 13 | 6406 | 34.4 |  |  |  |
|  |  |  | TTGATGCTACAATGAAG------------CTTTTCCCT | 12 | 417 | 2.2 |  |  |  |
|  |  |  | TTGATGCTACAATGAAGGA----GCGAAGCTTTTCCCT | 4 | 375 | 2.0 |  |  |  |

| | | | | | | | | | |
|---|---|---|---|---|---|---|---|---|---|
|  |  | 4 | ---------------------------AGCTTTTCCCT | 86 | 6682 | 29.9 | 98.5 |  |  |
|  |  |  | TTGATGCTACAATGAAGGAA--------GCT-----CT | 13 | 5317 | 23.8 |  |  |  |
|  |  |  | TTGATGCTACAATGAAGGAAC----GAAGCTTTTCCCT | 4 | 5096 | 22.8 |  |  |  |
| | | | TTGATGCTACAATGAAGGAACTAGC-AAGCTTTTCCCT | 1 | 4951 | 22.1 | | | |
|  |  | 5 | ---------------------------AGCTTTTCCCT | 86 | 8569 | 51.3 | 84.7 |  |  |
| | | | TTGATGCTACAATGAAGGAACTAGC-AAGCTTTTCCCT | 1 | 7241 | 43.3 | | | |
|  |  |  | TTGATGCTACAATGAAGGAAC----GAAGCTTTTCCCT | 4 | 82 | 0.5 |  |  |  |
|  |  |  | TTGATGCTACAATGAAGGAA--------GCT-----CT | 13 | 60 | 0.4 |  |  |  |
| | | 6 | TTGATGCTACAATGAAGGAACTAGC-AAGCTTTTCCCT | 1 | 9444 | 45.4 | 97.5 | | |
|  |  |  | ---------------------------AGCTTTTCCCT | 86 | 6124 | 29.4 |  |  |  |
|  |  |  | TTGATGCTACAATGAAGGAAC----GAAGCTTTTCCCT | 4 | 4617 | 22.2 |  |  |  |
|  |  |  | TTGATGCTACAATGAAGGAA--------GCT-----CT | 13 | 59 | 0.3 |  |  |  |
|  |  | 7 | ---------------------------AGCTTTTCCCT | 86 | 6261 | 30.5 | 97.3 |  |  |
|  |  |  | TTGATGCTACAATGAAGGAA--------GCT-----CT | 13 | 4989 | 24.3 |  |  |  |
| | | | TTGATGCTACAATGAAGGAACTAGC-AAGCTTTTCCCT | 1 | 4455 | 21.7 | | | |
|  |  |  | TTGATGCTACAATGAAGGAAC----GAAGCTTTTCCCT | 4 | 4258 | 20.7 |  |  |  |
|  |  | 8 | TTGATGCTACAATGAAGGAAC----GAAGCTTTTCCCT | 4 | 8904 | 47.4 | 93.0 |  |  |
|  |  |  | TTGATGCTACAATGAAGGAA--------GCT-----CT | 13 | 4698 | 25.0 |  |  |  |
| | | | TTGATGCTACAATGAAGGAACTAGC-AAGCTTTTCCCT | 1 | 4509 | 24.0 | | | |
|  |  |  | TTGATGCTACAATGAAGGA----GCAAAGCTTTTCCCT | 4 | 79 | 0.4 |  |  |  |
|  | 2 | 1 | ---------------------------AGCTTTTCCCT | 86 | 4748 | 29.3 | 96.3 | 88 | 8 |
|  |  |  | TTGATGCTACAATGAAGGAA--------GCT-----CT | 13 | 4059 | 25.1 |  |  |  |
| | | | TTGATGCTACAATGAAGGAACTAGC-AAGCTTTTCCCT | 1 | 3641 | 22.5 | | | |
|  |  |  | TTGATGCTACAATGAAGGAAC----GAAGCTTTTCCCT | 4 | 3409 | 21.1 |  |  |  |
|  |  | 2 | ---------------------------AGCTTTTCCCT | 86 | 4134 | 27.2 | 96.8 |  |  |
|  |  |  | TTGATGCTACAATGAAGGAA--------GCT-----CT | 13 | 3723 | 24.5 |  |  |  |
|  |  |  | TTGATGCTACAATGAAGGAAC----GAAGCTTTTCCCT | 4 | 3577 | 23.6 |  |  |  |
| | | | TTGATGCTACAATGAAGGAACTAGC-AAGCTTTTCCCT | 1 | 3408 | 22.4 | | | |
|  |  | 3 | TTGATGCTACAATGAAGGAA--------GCT-----CT | 13 | 6490 | 45.2 | 88.8 |  |  |
|  |  |  | ---------------------------AGCTTTTCCCT | 86 | 4232 | 29.4 |  |  |  |
|  |  |  | TTGATGCTACAATGAAGGATC----GAAGCTTTTCCCT | 4 | 3146 | 21.9 |  |  |  |
| | | | TTGATGCTACAATGAAGGAACTAGA-AAGCTTTTCCCT | 1 | 99 | 0.7 | | | |
| | | 4 | TTGATGCTACAATGAAGGAACTAGC-AAGCTTTTCCCT | 1 | 6844 | 46.6 | 73.9 | | |
|  |  |  | TTGATGCTACAATGAAGGAAC----GAAGCTTTTCCCT | 4 | 6834 | 46.5 |  |  |  |
|  |  |  | TTGATGCTACAATGAAGGAA--------GCT-----CT | 13 | 121 | 0.8 |  |  |  |
|  |  |  | ---------------------------AGCTTTTCCCT | 42 | 101 | 0.7 |  |  |  |
|  |  | 5 | TTGATGCTACAATGAAGGAAC----GAAGCTTTTCCCT | 4 | 12698 | 47.5 | 80.2 |  |  |
| | | | TTGATGCTACAATGAAGGAACTAGC-AAGCTTTTCCCT | 1 | 12045 | 45.1 | | | |
|  |  |  | TTGATGCTACAATGAAGGAA--------GCT-----CT | 13 | 264 | 1.0 |  |  |  |
|  |  |  | -----------------AGCTTTTCCCTGATGAAATTT | 42 | 187 | 0.7 |  |  |  |
|  |  | 6 | ---------------------------AGCTTTTCCCT | 86 | 4008 | 26.4 | 86.6 |  |  |
|  |  |  | TTGATGCTACAATGAAGGAA--------GCT-----CT | 13 | 3792 | 25.0 |  |  |  |
|  |  |  | TTGATGCTACAATGAAGGAAC----GAAGCTTTTCCCT | 4 | 3423 | 22.6 |  |  |  |
| | | | TTGATGCTACAATGAAGGAACTAGC-AAGCTTTTCCCT | 1 | 3076 | 20.3 | | | |
|  |  | 7 | TTGATGCTACAATGAAGGAA--------GCT-----CT | 13 | 7441 | 48.1 | 92.3 |  |  |
|  |  |  | TTGATGCTACAATGAAGGAAC----GAAGCTTTTCCCT | 4 | 6632 | 42.9 |  |  |  |
| | | | TTGATGCTACAATGAAGGATCTAGC-AAGCTTTTCCCT | 1 | 139 | 0.9 | | | |
|  |  |  | ---------------------------AGCTTTTCCCT | 26 | 131 | 0.8 |  |  |  |
| | | 8 | TTGATGCTACAATGAAGGAACTAGC-AAGCTTTTCCCT | 1 | 6537 | 43.4 | 92.0 | | |
|  |  |  | ---------------------------AGCTTTTCCCT | 42 | 4790 | 31.8 |  |  |  |
|  |  |  | TTGATGCTACAATGAAGGAAC----GAAGCTTTTCCCT | 4 | 3043 | 20.2 |  |  |  |
|  |  |  | TTGATGCTACAATGAAGGAA--------GCT-----CT | 13 | 71 | 0.5 |  |  |  |

*Bold red letters indicate TAM, and underlined letters denote reRNA target sequences

**Supplementary Table 15.** Indel types and frequencies in two *PDS* genes of progenies from parental plants infected with TRV2 expressing eTnpBc variant and $^{HH}$reRNA4$^{PDS}$

| Parent # | Gene ID | Plant # | Most common indels | n_deleted | Reads | % Edited reads | Overall editing frequency | Average editing frequency | Standard deviation |
|---|---|---|---|---|---|---|---|---|---|
| 1 | *PDS1* -Nbe05g35010 | | **TTGAT**GCTACAATGAAGGAACTAGCGAAGCTTTTCCCT* | Wild-type | | | | | |
|  |  | 1 | ---------------------------AGCTTTTCCCT | 86 | 10091 | 86.9 | 92.6 | 92 | 6 |
|  |  | 2 | ---------------------------AGCTTTTCCCT | 86 | 5563 | 48.9 | 86.6 |  |  |
|  |  |  | TTGATGCTACAATGAAGGAAC----GAAGCTTTTCCCT | 4 | 3915 | 34.4 |  |  |  |
|  |  | 3 | ---------------------------AGCTTTTCCCT | 86 | 7151 | 64.4 | 81.7 |  |  |
|  |  |  | TTGATGCTACAATGAAGGA----GCGAAGCTTTTCCCT | 4 | 259 | 2.3 |  |  |  |
|  |  |  | TTGATGCTACAACGAAGGAA-------AGCTTTTCCCT | 7 | 159 | 1.4 |  |  |  |
|  |  | 4 | ---------------------------AGCTTTTCCCT | 86 | 6681 | 54.9 | 98.4 |  |  |
|  |  |  | TTGATGCTACAATGAAGGAAC----GAAGCTTTTCCCT | 4 | 5057 | 41.5 |  |  |  |
|  |  | 5 | ---------------------------AGCTTTTCCCT | 86 | 8568 | 81.2 | 86.5 |  |  |
| | | | TTGATGCTACAATGAAGGAACTAGC-AAGCTTTTCCCT | 1 | 83 | 0.8 | | | |
|  |  | 6 | ---------------------------AGCTTTTCCCT | 86 | 6105 | 54.3 | 97.6 |  |  |
|  |  |  | TTGATGCTACAATGAAGGAAC----GAAGCTTTTCCCT | 4 | 4571 | 40.6 |  |  |  |
|  |  | 7 | ---------------------------AGCTTTTCCCT | 86 | 6258 | 56.1 | 97.1 |  |  |
|  |  |  | TTGATGCTACAATGAAGGAAC----GAAGCTTTTCCCT | 4 | 4230 | 37.9 |  |  |  |
|  |  | 8 | TTGATGCTACAATGAAGGAAC----GAAGCTTTTCCCT | 4 | 8809 | 88.5 | 92.6 |  |  |
| | | | TTGATGCTACAATGAAGGAACTAGC-AAGCTTTTCCCT | 1 | 82 | 0.8 | | | |
|  |  |  | TTGATGCTACAATGAAGGA----GCGAAGCTTTTCCCT | 1 | 57 | 0.6 |  |  |  |
| | *PDS2* -Nbe06g25970 | | **TTGAT**GCTACAATGAAGGAACTAGCAAAGCTTTTCCCT* | Wild-type | | | | | |
|  |  | 1 | TTGATGCTACAATGAAGGAA--------GCT-----CT | 13 | 4380 | 45.1 | 92.8 | 92 | 6 |
| | | | TTGATGCTACAATGAAGGAACTAGC-AAGCTTTTCCCT | 1 | 3918 | 40.4 | | | |
|  |  | 2 | TTGATGCTACAATGAAGGAA--------GCT-----CT | 13 | 4513 | 40.9 | 85.6 |  |  |
| | | | TTGATGCTACAATGAAGGAACTAGC-AAGCTTTTCCCT | 1 | 4268 | 38.7 | | | |
|  |  | 3 | TTGATGCTACAATGAAGGAA--------GCT-----CT | 13 | 6387 | 56.6 | 85.0 |  |  |
|  |  |  | TTGATGCTACAATGAAG------------CTTTTCCCT | 12 | 378 | 3.3 |  |  |  |
|  |  |  | TTGATGCTACAATGA-------------GCTTTTCCCT | 13 | 323 | 2.9 |  |  |  |
|  |  | 4 | TTGATGCTACAATGAAGGAA--------GCT-----CT | 13 | 5291 | 50.1 | 98.5 |  |  |
| | | | TTGATGCTACAATGAAGGAACTAGC-AAGCTTTTCCCT | 1 | 4868 | 46.1 | | | |
| | | 5 | TTGATGCTACAATGAAGGAACTAGC-AAGCTTTTCCCT | 1 | 7158 | 77.9 | 82.5 | | |
|  |  |  | TTGATGCTACAATGAAGGAA--------GCT-----CT | 13 | 58 | 0.6 |  |  |  |
|  |  |  | TTGATGCTACAATGA-------------GCTTTTCCCT | 13 | 52 | 0.6 |  |  |  |
| | | 6 | TTGATGCTACAATGAAGGAACTAGC-AAGCTTTTCCCT | 1 | 9290 | 92.1 | 97.4 | | |
|  |  |  | TTGATGCTACAATGAAGGAA--------GCT-----CT | 13 | 59 | 0.6 |  |  |  |
|  |  | 7 | TTGATGCTACAATGAAGGAA--------GCT-----CT | 13 | 4963 | 49.8 | 97.6 |  |  |
| | | | TTGATGCTACAATGAAGGAACTAGC-AAGCTTTTCCCT | 1 | 4374 | 43.9 | | | |
|  |  | 8 | TTGATGCTACAATGAAGGAA--------GCT-----CT | 13 | 4687 | 45.7 | 93.4 |  |  |
| | | | TTGATGCTACAATGAAGGAACTAGC-AAGCTTTTCCCT | 1 | 4427 | 43.2 | | | |
|  |  |  | TTGATGCTGCAATGAAGGAAC----GAAGCTTTTCCCT | 4 | 95 | 0.9 |  |  |  |
| 2 | *PDS1* -Nbe05g35010 | | **TTGAT**GCTACAATGAAGGAACTAGCGAAGCTTTTCCCT* | Wild-type | | | | | |
|  |  | 1 | ---------------------------AGCTTTTCCCT | 86 | 4748 | 54.7 | 96.3 | 88 | 8 |
|  |  |  | TTGATGCTACAATGAAGGAAC----GAAGCTTTTCCCT | 4 | 3370 | 38.8 |  |  |  |
| | | | TTGATGCTACAATGAAGGAACTAGC-AAGCTTTTCCCT | 1 | 95 | 1.1 | | | |
|  |  | 2 | ---------------------------AGCTTTTCCCT | 86 | 4134 | 50.9 | 97.0 |  |  |
|  |  |  | TTGATGCTACAATGAAGGAAC----GAAGCTTTTCCCT | 4 | 3537 | 43.6 |  |  |  |
|  |  | 3 | ---------------------------AGCTTTTCCCT | 86 | 4232 | 50.1 | 89.2 |  |  |
|  |  |  | TTGATGCTACAATGAAGGAAC----GAAGCTTTTCCCT | 4 | 3122 | 37.0 |  |  |  |
|  |  | 4 | TTGATGCTACAATGAAGGAAC----GAAGCTTTTCCCT | 4 | 6728 | 68.2 | 73.2 |  |  |
| | | | TTGATGCTACAATGAAGGAACTAGC-AAGCTTTTCCCT | 1 | 136 | 1.4 | | | |
|  |  |  | ---------------------------AGCTTTTCCCT | 86 | 101 | 1.0 |  |  |  |
|  |  | 5 | TTGATGCTACAATGAAGGAAC----GAAGCTTTTCCCT | 4 | 12557 | 75.2 | 80.5 |  |  |
|  |  |  | ---------------------------AGCTTTTCCCT | 44 | 180 | 1.1 |  |  |  |
| | | | TTGATGCTACAATGAAGGAACTAGC-AAGCTTTTCCCT | 1 | 179 | 1.1 | | | |
|  |  | 6 | ---------------------------AGCTTTTCCCT | 86 | 4008 | 44.7 | 86.7 |  |  |
|  |  |  | TTGATGCTACAATGAAGGAAC----GAAGCTTTTCCCT | 4 | 3387 | 37.8 |  |  |  |
|  |  | 7 | TTGATGCTACAATGAAGGAAC----GAAGCTTTTCCCT | 4 | 6594 | 83.5 | 91.1 |  |  |
|  |  |  | ---------------------------AGCTTTTCCCT | 86 | 122 | 1.5 |  |  |  |
|  |  | 8 | ---------------------------AGCTTTTCCCT | 86 | 4787 | 54.4 | 92.6 |  |  |
|  |  |  | TTGATGCTACAATGAAGGAAC----GAAGCTTTTCCCT | 4 | 3004 | 34.1 |  |  |  |
| | | | TTGATGCTACAATGAAGGAACTAGC-AAGCTTTTCCCT | 1 | 153 | 1.7 | | | |
| | *PDS2* -Nbe06g25970 | | **TTGAT**GCTACAATGAAGGAACTAGCAAAGCTTTTCCCT* | Wild-type | | | | | |
|  |  | 1 | TTGATGCTACAATGAAGGAA--------GCT-----CT | 13 | 4036 | 49.6 | 96.3 | 88 | 8 |
| | | | TTGATGCTACAATGAAGGAACTAGC-AAGCTTTTCCCT | 1 | 3546 | 43.6 | | | |
|  |  | 2 | TTGATGCTACAATGAAGGAA--------GCT-----CT | 13 | 3712 | 49.0 | 96.6 |  |  |
| | | | TTGATGCTACAATGAAGGAACTAGC-AAGCTTTTCCCT | 1 | 3344 | 44.1 | | | |
|  |  | 3 | TTGATGCTACAATGAAGGAA--------GCT-----CT | 13 | 6458 | 83.5 | 88.4 |  |  |
| | | | TTGATGCTACAATGAAGGAACTAGC-AAGCTTTTCCCT | 1 | 85 | 1.1 | | | |
| | | 4 | TTGATGCTACAATGAAGGAACTAGC-AAGCTTTTCCCT | 1 | 6708 | 67.0 | 74.5 | | |
|  |  |  | TTGATGCTACAATGAAGGAA--------GCT-----CT | 13 | 116 | 1.2 |  |  |  |
|  |  |  | TTGATGCTACAATGAAGGAAC----GAAGCTTTTCCCT | 4 | 107 | 1.1 |  |  |  |

| | | | | | | | |
|---|---|---|---|---|---|---|---|
| | | 5 | TTGATGCTACAATGAAGGAACTAGC-AAGCTTTTCCCT | 1 | 11868 | 71.5 | 79.9 |
|  |  |  | TTGATGCTACAATGAAGGAA--------GCT-----CT | 13 | 262 | 1.6 |  |
|  |  |  | TTGATGCTACAATGAAGGAAC----GAAGCTTTTCCCT | 4 | 141 | 0.8 |  |
|  |  | 6 | TTGATGCTACAATGAAGGAA--------GCT-----CT | 13 | 3774 | 44.2 | 86.6 |
| | | | TTGATGCTACAATGAAGGAACTAGC-AAGCTTTTCCCT | 1 | 3028 | 35.5 | |
|  |  | 7 | TTGATGCTACAATGAAGGAA--------GCT-----CT | 13 | 7404 | 83.5 | 93.5 |
| | | | TTGATGCTACAATGAAGGAACTAGC-AAGCTTTTCCCT | 1 | 134 | 1.5 | |
| | | 8 | TTGATGCTACAATGAAGGAACTAGC-AAGCTTTTCCCT | 1 | 6384 | 84.6 | 91.3 |
|  |  |  | TTGATGCTACAATGAAGGAA--------GCT-----CT | 13 | 69 | 0.9 |  |

*Bold red letters indicate TAM, and underlined letters denote reRNA target sequences

**Supplementary Table 16.** Indel types and frequencies in *PDS* of progenies from parental plants infected with TRV2 expressing TnpB and reRNA4$^{PDS}$

| Construct | Parent # | Plant # | 4 most common indels | n_deleted | Reads | % Edited reads | Overall editing frequency | Average editing frequency | Standard deviation |
|---|---|---|---|---|---|---|---|---|---|
| | | | **TTGAT**GCTACAATGAAGGAACTAGCGAAGCTTTTCCCT* | Wild-type | | | | | |
| TnpB-reRNA4$^{PDS}$ | 1 | 1 | TTGATGCTACAATGAAGGAACTAGCGAAGC-TTTCCCT | 1 | 59 | 11.6 | 1.5 | 2 | 1 |
|  |  |  | TTGATGCTACAATGAAGGAACTA----AGCTTTTCCCT | 4 | 48 | 9.4 |  |  |  |
|  |  |  | TTGATGCTACAATG-------TAGCAAAGCTTTTCCCT | 7 | 47 | 9.2 |  |  |  |
|  |  |  | TTGATGCTACAATGAAG------------CTTTTCCCT | 12 | 46 | 9.0 |  |  |  |
|  |  | 2 | TTGATGCTACAATGAAG------------CTTTTCCCT | 12 | 130 | 13.3 | 3.6 |  |  |
|  |  |  | TTGATGCTACAATGA-------------GCTTTTCCCT | 13 | 74 | 7.6 |  |  |  |
|  |  |  | TTGATGCTACAATGAAGGAA--------GCTTTTCCCT | 8 | 62 | 6.4 |  |  |  |
|  |  |  | TTGATGCTACAATGAAGGA----GCAAAGCTTTTCCCT | 4 | 61 | 6.3 |  |  |  |
| | | 3 | TTGATGCTACAATGAAGGAACTAGCAAAGC-TTTCCAT | 1 | 23 | 13.9 | 0.7 | | |
|  |  |  | TTGATGCTACAATGAAGGAA-------AGCTTTTCCCT | 7 | 22 | 13.3 |  |  |  |
|  |  |  | TTGATGCTACAATGAAG------------CCTTTCCCT | 12 | 21 | 12.7 |  |  |  |
|  |  |  | TTGATGCTACAATGAAGGAACTAGC-----TTTTCCCT | 5 | 14 | 8.5 |  |  |  |
|  |  | 4 | TTGATGCTACAATGAAG------------CTTTTCCCT | 12 | 55 | 19.5 | 1.7 |  |  |
| | | | TTGATGCTACAATGAAGGAACTAGCAAAGC-TTTCCCT | 1 | 26 | 9.2 | | | |
|  |  |  | TTGATGCTACAATGAAGGAA--------GCTTTTCCCT | 8 | 24 | 8.5 |  |  |  |
|  |  |  | TTGATGCTACAATGAAGGAAT----GAAGCTTTTCCCT | 4 | 21 | 7.4 |  |  |  |
|  |  | 5 | TTGATGCTACAATGAAGGAA--------GCTTTTCCCT | 8 | 19 | 10.4 | 1.5 |  |  |
|  |  |  | TTGATGCTACAATGAAGGA------------------- | 30 | 17 | 9.3 |  |  |  |
|  |  |  | TTGATGCTACAATGAAG------------CTTTTCCCT | 12 | 14 | 7.7 |  |  |  |
| | | | TTGATGCTACAATGAAGGAA---GCAACGCTTTTCCCT | 3 | 13 | 7.1 | | | |
|  |  | 6 | TTGATGCTACAATGAAGGAA--------GCTTTTCCCT | 8 | 37 | 22.7 | 1.0 |  |  |
|  |  |  | TTGATGCTACAATGAAGTAA-------AGCTTTTCCCT | 7 | 35 | 21.5 |  |  |  |
|  |  |  | TTGATGCTACAATGAAGGAAC----------------- | 31 | 17 | 10.4 |  |  |  |
|  |  |  | TTGATGCTACAATGAAGG-----------CTTTTCCCT | 11 | 9 | 5.5 |  |  |  |
| | | 7 | TTGATGCTACAATGAAGGAACTAGCAAAGC-TTTCCCT | 1 | 27 | 32.5 | 0.4 | | |
|  |  |  | TTGATGCTACAAT------------------------- | 60 | 12 | 14.5 |  |  |  |
|  |  |  | TTGATGCTACAATGAAG------------CTTTTCCCT | 12 | 10 | 12.0 |  |  |  |
|  |  |  | TTGATGCTACAAT--------TAGCAAAGCTTTTCCCT | 8 | 7 | 8.4 |  |  |  |
|  |  | 8 | TTGATGCTACAATGAAGGAA--------GCTTTTCCCT | 8 | 95 | 18.5 | 2.2 |  |  |
|  |  |  | TTGATGCTACAATGAA-----------AGCTTTTCCCT | 11 | 56 | 10.9 |  |  |  |
|  |  |  | TTGATGCTACAATGAAGGA----GCGAAGCTTTTCCCT | 4 | 50 | 9.7 |  |  |  |
|  |  |  | TTGATGCTAC------------AGC-AATCTTTTACCT | 18 | 35 | 6.8 |  |  |  |
|  | 2 | 1 | TTGATGCTACAATGAAG------------CTTTTCCCT | 12 | 332 | 12.3 | 31.6 | 33 | 2 |
|  |  |  | TTGATGCTACAATGAAGGAA--------GCTTTTCCCT | 8 | 246 | 9.1 |  |  |  |
|  |  |  | TTGATGCTACAATGAA-----------AGCTTTTCCCT | 11 | 174 | 6.4 |  |  |  |
|  |  |  | TTGATGCTACAATGAAGGAAT----GAAGCTTTTCCCT | 4 | 172 | 6.4 |  |  |  |
|  |  | 2 | TTGATGCTACAATGAAG------------CTTTTCCCT | 12 | 262 | 12.7 | 35.2 |  |  |
|  |  |  | TTGATGCTACAATGAAGGAA--------GCTTTTCCCT | 8 | 192 | 9.3 |  |  |  |
|  |  |  | TTGATGCTACAATGA-------------GCTTTTGCCT | 13 | 126 | 6.1 |  |  |  |
|  |  |  | TTGATGCTACAATGAAGGAC-------AGCTTTTCCCT | 7 | 118 | 5.7 |  |  |  |
|  |  | 3 | TTGATGCTACAATGAAG------------CTTTTCCCT | 12 | 344 | 15.5 | 28.9 |  |  |
|  |  |  | TTGATGCTACAATGAAGGAA--------GCTTTTCCCT | 8 | 189 | 8.5 |  |  |  |
|  |  |  | TTGATGCTACAATGA-------------GCTTTTCCCT | 13 | 185 | 8.3 |  |  |  |
|  |  |  | TTGATGCTACAATGAAGTAA-------AGCTTTTCCCT | 7 | 149 | 6.7 |  |  |  |
|  |  | 4 | TTGATGCTACAATGAAG------------CTTTTCCCT | 12 | 429 | 14.4 | 30.1 |  |  |
|  |  |  | TTGATGCTACAATGAAGGAA--------GCTTTTCCCT | 8 | 287 | 9.6 |  |  |  |
|  |  |  | TTGATGCTACAAGGA-------------GCTTTTCCCT | 13 | 198 | 6.6 |  |  |  |
|  |  |  | TTGATGCTACAATGAA-----------AACTTTTCCCT | 11 | 174 | 5.8 |  |  |  |
|  |  | 5 | TTGATGCTACAATGAAG------------CTTTTCCCT | 12 | 428 | 15.3 | 35.3 |  |  |
|  |  |  | TTGATGCTACAATGAAGGAA--------GCTTTTCCCT | 8 | 277 | 9.9 |  |  |  |
|  |  |  | TTGATGCTACAATGA-------------GCTTTTCCCT | 13 | 218 | 7.8 |  |  |  |
|  |  |  | TTGATGCTACAATGAAGGAAC----GAAGCTTTTCCCT | 4 | 178 | 6.4 |  |  |  |
|  |  | 6 | TTGATGCTACAATGAAG------------CTTTTCCCT | 12 | 461 | 15.0 | 33.7 |  |  |
|  |  |  | TTGATGCTACAATGAAGGAA--------GCCTTTCCCT | 8 | 299 | 9.7 |  |  |  |
|  |  |  | TTGATGCTACAATGA-------------GCTTTTCCCT | 13 | 208 | 6.8 |  |  |  |
|  |  |  | TTGATGCTACAATGAGGGAAC----GAAGCTTTTCCCT | 4 | 196 | 6.4 |  |  |  |
|  |  | 7 | TTGATGCTACAATGAAG------------CTTTTCCCT | 12 | 326 | 11.7 | 32.0 |  |  |
|  |  |  | TTGATGCTACACTGAAGGAA--------GCTTTTCCCT | 8 | 287 | 10.3 |  |  |  |
|  |  |  | TTGATGCTACAATGA-------------GCTTTTCCCT | 13 | 181 | 6.5 |  |  |  |
|  |  |  | TTGATGCTACAATGAA-----------AGCTTTTCCCT | 11 | 166 | 5.9 |  |  |  |

| | | | | | | | | |
|---|---|---|---|---|---|---|---|---|
|  |  | 8 | TTGATGCTACAATGAAG------------CTTTTCCCT | 12 | 398 | 13.1 | 34.4 |  |  |
|  |  |  | TTGATGCTACACTGAAGGAA--------GCTTTTCCCT | 8 | 313 | 10.3 |  |  |  |
|  |  |  | TTGATGCTACAATGA-------------GCTTTTCCCT | 13 | 216 | 7.1 |  |  |  |
|  |  |  | TTGATGCTACAATGAAGGAC-------AGCTTTTGCCT | 7 | 180 | 5.9 |  |  |  |
| TnpB-[HH]reRNA4[PDS] | 1 | 1 | TTGATGCTACAATGAAG------------CTTTTCCCT | 12 | 505 | 14.3 | 32.0 | 34 | 1 |
|  |  |  | TTGATGCTACAATGAAGGAA--------GCTTTTTCCT | 8 | 330 | 9.4 |  |  |  |
|  |  |  | TTGATGCTACAATGA-------------GCTTTTCCCT | 13 | 224 | 6.4 |  |  |  |
|  |  |  | TTGATGCTACAATGAAGGAAC----GAAGCTTTTCCCT | 4 | 195 | 5.5 |  |  |  |
|  |  | 2 | TTGATGCTACAATGAAG------------CTTTTCCCT | 12 | 368 | 13.3 | 33.6 |  |  |
|  |  |  | TTGATGCTACAATGAAGGAA--------GCTTTTCCCT | 8 | 264 | 9.6 |  |  |  |
|  |  |  | TTGATGCTACAATGA-------------GCTTTTCCCT | 13 | 174 | 6.3 |  |  |  |
|  |  |  | TTGATGCTACAATGAAGGAA-------AGCTTTTCCCT | 7 | 173 | 6.3 |  |  |  |
|  |  | 3 | TTGATGCTACAATGAAG------------CTTTTCCCT | 12 | 433 | 14.2 | 33.7 |  |  |
|  |  |  | TTGATGCTACAATGAAGGAA--------GCTTTTCCCT | 8 | 337 | 11.0 |  |  |  |
|  |  |  | TTGATGCTACAATGAAGGAAC----GAAGCTTTTCCCT | 4 | 224 | 7.3 |  |  |  |
|  |  |  | TTGATGCTACAATGA-------------GCTTTTCCCT | 13 | 161 | 5.3 |  |  |  |
|  |  | 4 | TTGATGCTACAATGAAG------------CTTTTCCCT | 12 | 261 | 11.8 | 31.5 |  |  |
|  |  |  | TTGATGCTACAATGAAGGAA--------GCTTTTCCCT | 8 | 200 | 9.1 |  |  |  |
|  |  |  | TTGATGCTACAATGA-------------GCTTTTCCCT | 13 | 176 | 8.0 |  |  |  |
|  |  |  | TTGATGCTACAATGAAGGATC----GAAGCTTTTCCCT | 4 | 149 | 6.8 |  |  |  |
|  |  | 5 | TTGATGCTACAATGAAG------------CTTTTCCCT | 12 | 374 | 14.8 | 34.4 |  |  |
|  |  |  | TTGATGCTACAATGAAGGAA--------GCTTTTCCCT | 8 | 231 | 9.1 |  |  |  |
|  |  |  | TTGATGCTACAATGA-------------GCTTTTCCCT | 13 | 172 | 6.8 |  |  |  |
|  |  |  | TTGATGCTACAATGAAGGAAC----GAAGCTTGTTCCC | 4 | 152 | 6.0 |  |  |  |
|  |  | 6 | TTGATGCTACAATGAAG------------CTTTTCCCT | 12 | 196 | 13.7 | 35.8 |  |  |
|  |  |  | TTGATGCTACAATGAAGGAA--------GCTTTTCCCC | 8 | 128 | 9.0 |  |  |  |
|  |  |  | TTGATGCTACAATGA-------------GCTTTTCCCT | 13 | 107 | 7.5 |  |  |  |
|  |  |  | TTGATGCTACAATGAA-----------AGCTTTTCCCT | 11 | 83 | 5.8 |  |  |  |
|  |  | 7 | TTGATGCTACAATGAAG------------CTTTTCCCT | 12 | 166 | 11.1 | 33.3 |  |  |
|  |  |  | TTGATGCTACAATGAAGGAA--------GCTTTTCCCT | 8 | 149 | 9.9 |  |  |  |
|  |  |  | TTGATGCTACAATGA-------------GCTTTTCCCT | 13 | 124 | 8.3 |  |  |  |
|  |  |  | TTGATGCTACAATGAAGTAA-------AGCTTTTCCCT | 7 | 83 | 5.5 |  |  |  |
|  |  | 8 | TTGATGCTACAATGAAG------------CTTTTCCCT | 12 | 339 | 14.8 | 34.6 |  |  |
|  |  |  | TTGATGCTACAATGAAGGAA--------GCTTTTCCCT | 8 | 182 | 8.0 |  |  |  |
|  |  |  | TTGATGCTACAATGAAGCAA-------AGCTTTTCCCT | 7 | 158 | 6.9 |  |  |  |
|  |  |  | TTGATGCTACAATGA-------------GCTTTTCCCT | 13 | 148 | 6.5 |  |  |  |
|  | 2 | 1 | TTGATGCTACAATGAAG------------CTTTTCCCT | 12 | 269 | 12.2 | 30.6 | 32 | 3 |
|  |  |  | TTGATGCTACAATGAAGGAA--------GCCTTTCCCT | 8 | 208 | 9.4 |  |  |  |
|  |  |  | TTGATGCTACAATGA-------------GCTTTTCCCT | 13 | 171 | 7.7 |  |  |  |
|  |  |  | TTGATGCTACAATGAA-----------AGCTTTTCCCT | 11 | 158 | 7.1 |  |  |  |
|  |  | 2 | TTGATGCTACAATGAAG------------CTTTTCCCT | 12 | 253 | 14.4 | 27.0 |  |  |
|  |  |  | TTGATGCTACAATGAAGGAA--------GCTTTTCCCT | 8 | 181 | 10.3 |  |  |  |
|  |  |  | TTGATGCTACGATGA-------------GCTTTTCCCT | 13 | 129 | 7.3 |  |  |  |
|  |  |  | TTGATGCTACAATGAAGCAA-------AGCTTTTCCCT | 7 | 117 | 6.7 |  |  |  |
|  |  | 3 | TTGATGCTACAATGAAG------------CTTTTCCCT | 12 | 381 | 15.3 | 34.2 |  |  |
|  |  |  | TTGATGCTACAATGAAGGAA--------GCTTTTCCCT | 8 | 238 | 9.5 |  |  |  |
|  |  |  | TTGATGCTACAATGA-------------GCTTTTCCCT | 13 | 185 | 7.4 |  |  |  |
|  |  |  | TTGATGCTACAATGAAGGAC-------AGCTTTTCCCT | 7 | 126 | 5.1 |  |  |  |
|  |  | 4 | TTGATGCTACAATGAAG------------CTTTTCCCT | 12 | 366 | 18.0 | 28.8 |  |  |
|  |  |  | TTGATGCTACAATGAAGGAA--------GCTTTTCCCT | 8 | 181 | 8.9 |  |  |  |
|  |  |  | TTGATGCTACAATGA-------------GCTTTTCCCT | 13 | 140 | 6.9 |  |  |  |
|  |  |  | TTGATGCTACAATGAA-----------AGCTTTTCCCT | 11 | 121 | 5.9 |  |  |  |
|  |  | 5 | TTGATGCTACAATGAAG------------CTTTTCCCT | 12 | 265 | 12.7 | 32.4 |  |  |
|  |  |  | TTGATGCTACAATGAAGGAA--------GCTTTTCCCT | 8 | 193 | 9.3 |  |  |  |
|  |  |  | TTGATGCTACAATGA-------------GCTTTTCCCT | 13 | 167 | 8.0 |  |  |  |
|  |  |  | TTGATGCTACAATGAAGGAA-------AGCTTTTCGCT | 7 | 121 | 5.8 |  |  |  |
|  |  | 6 | TTGATGCTACAATGAAG------------CTTTTCCCT | 12 | 413 | 13.5 | 34.8 |  |  |
|  |  |  | TTGATGCTACAATGAAGGAA--------GCTTTTCCCT | 8 | 287 | 9.4 |  |  |  |
|  |  |  | TTGATGCTACAATGA-------------GCTTTTCCCT | 13 | 218 | 7.1 |  |  |  |
|  |  |  | TTGATGCTACAATGAAGGAC-------AGCTTGTCCCT | 7 | 177 | 5.8 |  |  |  |
|  |  | 7 | TTGATGCTACAATGAAG------------CTTTTCCCT | 12 | 324 | 14.0 | 33.2 |  |  |
|  |  |  | TTGATGCTACAATGAAGGAA--------GCTTTTCCCT | 8 | 237 | 10.2 |  |  |  |
|  |  |  | TTGATGCTACAATGAAGGAA-------AGCTTTTCCCT | 7 | 146 | 6.3 |  |  |  |
|  |  |  | TTGATGCTACAATGAAGGAAC----GAAGCTTTTCCAT | 4 | 139 | 6.0 |  |  |  |
|  |  | 8 | TTGATGCTACAATGAAG------------CTTTTCCCT | 12 | 358 | 15.2 | 33.2 |  |  |
|  |  |  | TTGATGCTACAATGAAGGAA--------GCTTTTCCCT | 8 | 230 | 9.8 |  |  |  |
|  |  |  | TTGATGCTACAATGA-------------GCTTTTCCCT | 13 | 167 | 7.1 |  |  |  |
|  |  |  | TTGATGCTACAATGAAGGAC-------AGCTTTTCCCT | 7 | 138 | 5.9 |  |  |  |

*Bold red letters indicate TAM, and underlined letters denote reRNA target sequences

**Supplementary Table 17.** Indel types and frequencies in *PDS* of progenies from parental plants infected with TRV2 expressing eTnpBe variant and reRNA4$^{PDS}$

| Construct | Parent # | Plant # | Most common indels | n_deleted | Reads | % Edited reads | Overall editing frequency | Average editing frequency | Standard deviation |
|---|---|---|---|---|---|---|---|---|---|
| | | | **TTGAT**GCTACAATGAAGGAACTAGCGAAGCTTTTCCCT* | Wild-type | | | | | |
| eTnpBe-reRNA4$^{PDS}$ | 1 | 1 | TTGATGCTACAATGAAG------------CTTTTCCCT | 12 | 308 | 7.5 | 9.1 | 16 | 7 |
|  |  |  | TTGATGCTACAATGAAGGAA-------AGCTTTTCCCT | 7 | 260 | 6.3 |  |  |  |
|  |  |  | TTGATGCTACAATGAAGGAA--------GCTTTTCCCT | 8 | 244 | 5.9 |  |  |  |
|  |  |  | -------------------------------------- | 56 | 237 | 5.7 |  |  |  |
|  |  | 2 | TTGATGCTACAATGAAGGA----GCGAAGCTTTTCCCT | 4 | 1202 | 9.2 | 21.4 |  |  |
|  |  |  | TTGATGCTACATTGAAG------------CTTTTCCCT | 12 | 1094 | 8.3 |  |  |  |
|  |  |  | TTGATGCTACAATGAAGGAA--------GCTTTTCCCT | 8 | 1054 | 8.0 |  |  |  |
|  |  |  | TTGATGCTACAATGAAGGAA-------AGCTTTTCCCT | 7 | 717 | 5.5 |  |  |  |
|  |  | 3 | TTGATGCTACAATGAAG------------CTTTTCCCT | 12 | 454 | 9.0 | 10.8 |  |  |
|  |  |  | TTGATGCTACAATGAAGGA----GCGAAGCTTTTCCCT | 4 | 356 | 7.0 |  |  |  |
|  |  |  | TTGATGCTACAATGAAGGAA--------GCTTTTCCCT | 8 | 335 | 6.6 |  |  |  |
|  |  |  | TTGATGCTACAATGAAGGAA-------AGCTTTTCCCT | 7 | 261 | 5.1 |  |  |  |
|  |  | 4 | TTGATGCTACAATGAAG------------CTTTTCCCT | 12 | 1027 | 12.5 | 26.5 |  |  |
|  |  |  | TTGATGCTACAATGAAGGA----GCGAAGCTTTTCCCT | 4 | 680 | 8.3 |  |  |  |
|  |  |  | TTGATGCTACAATGAAGGAAC----GAAGCTTTTCCCT | 4 | 479 | 5.8 |  |  |  |
|  |  |  | TTGATGCTACAAAGAAGGAA--------GCTTTTCCCT | 8 | 436 | 5.3 |  |  |  |
|  |  | 5 | TTGATGCTACAATGAAG------------CTTTTCCCT | 12 | 965 | 9.2 | 21.6 |  |  |
|  |  |  | TTGATGCTACAATGAAGGAA-------AGCTCTTCCCT | 7 | 800 | 7.6 |  |  |  |
|  |  |  | TTGATGCTACAATGAAGGAA--------GCTTTTCCCT | 8 | 705 | 6.7 |  |  |  |
|  |  |  | TTGATGCTACAATGAAGGAAC----GAAGCTTTTCCCT | 4 | 564 | 5.4 |  |  |  |
|  |  | 6 | TTGATGCTACAATGAAG------------CTTTTCCCT | 12 | 1397 | 12.8 | 20.7 |  |  |
|  |  |  | TTGATGCTACAATGAAGGAAC----GAAGCTTTTCCCT | 4 | 940 | 8.6 |  |  |  |
|  |  |  | TTGATGCTACAATGAAGGAA-------AGCTTTTCCCT | 7 | 853 | 7.8 |  |  |  |
|  |  |  | TTGATGCTACAATGAAGGA----GCAAAGCTCTTCCCT | 4 | 756 | 6.9 |  |  |  |
|  |  | 7 | TTGATGCTACAATGAAG------------CTTTTCCCT | 12 | 740 | 10.2 | 8.4 |  |  |
|  |  |  | TTGATGCTACAATGAAGGAA-------AGCTTTTCCCT | 7 | 496 | 6.8 |  |  |  |
|  |  |  | TTGATGCTACAATGA-------------GCTTTTCCCT | 13 | 487 | 6.7 |  |  |  |
|  |  |  | TTGATGCTACAATGAATGA----GCGAAGCTTTTCCCT | 4 | 438 | 6.0 |  |  |  |
|  |  | 8 | TTGATGCTACAATGAAGGAA-------AGCTTTTCCCT | 7 | 306 | 7.7 | 8.6 |  |  |
|  |  |  | TTGATGCTACAATGAAG------------CTTTTCCCT | 12 | 301 | 7.6 |  |  |  |
|  |  |  | TTGATGCTACACTGAAGGA----GCGAAGCTTTTCCCT | 4 | 298 | 7.5 |  |  |  |
|  |  |  | TTGATGCTACAATGAAGGAA--------GCTTTTCCCT | 8 | 184 | 4.6 |  |  |  |
|  | 2 | 1 | TTGATGCTACAATGAAG------------CTTTTCCCT | 12 | 320 | 13.1 | 30.1 | 24 | 14 |
|  |  |  | TTGATGCTACAATGA-------------GCTTTTCCCT | 13 | 240 | 9.8 |  |  |  |
|  |  |  | TTGATGCTACAATGAAGGAA--------GCTTTTCCCT | 8 | 197 | 8.0 |  |  |  |
|  |  |  | TTGATGCTACAATGAAGGAAC----GAAGCTTTTCCCT | 4 | 165 | 6.7 |  |  |  |
|  |  | 2 | TTGATGCTACAATGAAG------------CTTTTCCCT | 12 | 445 | 14.4 | 28.4 |  |  |
|  |  |  | TTGATGCTACAATGAAGGAA--------GCTTTTCCCT | 8 | 313 | 10.1 |  |  |  |
|  |  |  | TTGATGCTACAATGA-------------GCTTTTCCCT | 13 | 247 | 8.0 |  |  |  |
|  |  |  | TTGATGCTACAATGAATGAAC----GAAGCTTTTCCCT | 4 | 182 | 5.9 |  |  |  |
|  |  | 3 | TTGATGCTACAATGAAG------------CTTTTCCCT | 12 | 495 | 13.5 | 33.1 |  |  |
|  |  |  | TTGATGCTACAATGGAGGAA--------GCTTTTCCCT | 8 | 313 | 8.6 |  |  |  |
|  |  |  | TTGATGCTACAATGA-------------GCTTTTCTCT | 13 | 309 | 8.5 |  |  |  |
|  |  |  | TTGATGCTACAATGAAGGAAC----GAAACTTTTCCCT | 4 | 246 | 6.7 |  |  |  |
|  |  | 4 | TTGATGCTACAATGAAG------------CTTTTCCCT | 12 | 416 | 15.2 | 33.3 |  |  |
|  |  |  | TTGATGCTACAATGAAGGAA--------GCTTTTCCCT | 8 | 204 | 7.5 |  |  |  |
|  |  |  | TTGATGCTACAATGA-------------GCTTTTCCCT | 13 | 187 | 6.9 |  |  |  |
|  |  |  | TTGATGCTACAATGAA-----------AGCTTTTCCCT | 11 | 172 | 6.3 |  |  |  |
|  |  | 5 | TTGATGCTACAATGAAG------------CTTTTCCCT | 12 | 370 | 12.3 | 34.3 |  |  |
|  |  |  | TTGATGCTACAATGAAGGAA--------GCTTTTCCCT | 8 | 281 | 9.3 |  |  |  |
|  |  |  | TTGATGCTACAATGAAGGAAC----GAAACTTTTCCCT | 4 | 247 | 8.2 |  |  |  |
|  |  |  | TTGATGCTACAATGA-------------GCTTTTCCCT | 13 | 209 | 6.9 |  |  |  |
|  |  | 6 | TTGATGCTACAATGAAG------------CTTTTCCCT | 12 | 376 | 13.4 | 33.3 |  |  |
|  |  |  | TTGATGCTACAATGAAGGAA--------GCTTTTCCCT | 8 | 295 | 10.5 |  |  |  |
|  |  |  | TTGATGCTACAATGA-------------GCTTTTCCCT | 13 | 172 | 6.1 |  |  |  |
|  |  |  | TTGATGCTACAATGAA-----------AGCTTTTCCCT | 11 | 156 | 5.6 |  |  |  |
|  |  | 7 | TTGATGCTACAATGAAGGAA--------GCTTTTCCCT | 8 | 40 | 14.3 | 2.3 |  |  |
|  |  |  | TTGATGCTACAATGAAG------------CTTTTCCCT | 12 | 37 | 13.2 |  |  |  |
|  |  |  | TTGATGCTACAATGAAGGA----GCAAAGC-TTTCTCT | 5 | 18 | 6.4 |  |  |  |
|  |  |  | TTGATGCTACAATGAAGGA----------CTTTTCCCT | 10 | 17 | 6.1 |  |  |  |

| | | | | | | | | | |
|---|---|---|---|---|---|---|---|---|---|
|  |  | 8 | TTGATGCTACAATGAAG------------CTTTTCCCT | 12 | 20 | 11.6 | 1.1 |  |  |
| | | | TTGATGCTACAATGAAGG---TAGCGAAGCTTTTCCCT | 3 | 16 | 9.2 | | | |
|  |  |  | TTGATGCTACAATGA-------------GCTTTTCCCT | 13 | 14 | 8.1 |  |  |  |
|  |  |  | TTGATGCTACAATGAAGGAG--------GCTTTTCCCT | 8 | 13 | 7.5 |  |  |  |
| eTnpBe-$^{HH}$reRNA4$^{PDS}$ | 1 | 1 | TTGATGCTACAATGAAG------------CTTTTCCCT | 12 | 676 | 12.4 | 28.5 | 23 | 8 |
|  |  |  | TTGATGCTACAATGAAGGAA--------GCTTTTCCCT | 8 | 508 | 9.4 |  |  |  |
|  |  |  | TTGATGCTACAATGAAGGA----GCAAAGCTTTTCCCT | 4 | 382 | 7.0 |  |  |  |
|  |  |  | TTGATGCTACAATGAAGTAA-------AGCTTTTCCCT | 7 | 303 | 5.6 |  |  |  |
|  |  | 2 | TTGATGCTACAATGAAG------------CTTTTCCCT | 12 | 520 | 17.3 | 21.1 |  |  |
|  |  |  | TTGATGCTACAATGAAGGAA--------GCTTTTCCCT | 8 | 200 | 6.7 |  |  |  |
|  |  |  | TTGATGCTACAATGAAGGA----GCAAAGCCTTTCCCT | 4 | 151 | 5.0 |  |  |  |
|  |  |  | -------------------------------------- | 65 | 134 | 4.5 |  |  |  |
|  |  | 3 | TTGATGCTACAATGAAG------------CTTTTCCCT | 12 | 911 | 16.2 | 36.9 |  |  |
|  |  |  | TTGATGCTACAATGAAGGAA--------GCTTTTCCCT | 8 | 489 | 8.7 |  |  |  |
|  |  |  | TTGATGCTACAATGAAGGA----GCAAAGCTTTTCCCT | 4 | 410 | 7.3 |  |  |  |
|  |  |  | TTGATGCTACAATGAAGGAAC----GAAGCTTTTCCCT | 4 | 326 | 5.8 |  |  |  |
|  |  | 4 | TTGATGCTACAATGAAG------------CTTTTCCCT | 12 | 484 | 11.6 | 27.5 |  |  |
|  |  |  | TTGATGCTACAATGAAGGTA--------GCTTTTCCCT | 8 | 366 | 8.7 |  |  |  |
|  |  |  | TTGATGCTACAATGAAGGAA-------AGCTTTTCCCT | 7 | 284 | 6.8 |  |  |  |
|  |  |  | TTGATGCTACAATGA-------------GCTTTTCCCT | 13 | 214 | 5.1 |  |  |  |
|  |  | 5 | TTGATGCTACAATGAAG------------CTTTTCCCT | 12 | 447 | 12.3 | 22.7 |  |  |
|  |  |  | TTGATGCTACAATGAAGGAA--------GCTTTTCCCT | 8 | 250 | 6.9 |  |  |  |
|  |  |  | TTGATGCTACAATGAAGGA----GTGAAGCTTTTCCCT | 4 | 246 | 6.8 |  |  |  |
|  |  |  | TTGATGCTACAATGA-------------GCTTTTCCCT | 13 | 154 | 4.2 |  |  |  |
|  |  | 6 | TTGATGCTACAATGAAG------------CTTTTCCCT | 12 | 743 | 13.6 | 24.6 |  |  |
|  |  |  | TTGATGCTACAATGAAGGA----GCGAAGCTTTTCCCT | 4 | 408 | 7.5 |  |  |  |
|  |  |  | TTGATGCTACAATGAAGGAAC----GAAGCTTTTCCCT | 4 | 404 | 7.4 |  |  |  |
|  |  |  | TTGATGCTACAATGAAGGAC-------AGCTTTTCCCT | 7 | 368 | 6.7 |  |  |  |
|  |  | 7 | TTGATGCTACAATGAAG------------CTTTTCCCT | 12 | 128 | 9.1 | 8.4 |  |  |
|  |  |  | TTGATGCTACAATGAAGGAA--------GCTTTTCCCT | 8 | 80 | 5.7 |  |  |  |
|  |  |  | TTGATGCTACAATGAAGGAAC----GAAGTTTTTCCCT | 4 | 68 | 4.8 |  |  |  |
|  |  |  | TTGATGCTACAATGAAGGAC-------AGCTTTTCCCT | 7 | 61 | 4.3 |  |  |  |
|  |  | 8 | TTGATGCTACAATGAAG------------CTTTTCCCT | 12 | 316 | 11.3 | 16.4 |  |  |
|  |  |  | TTGATGCTACAATGAAGGAA-------AGCCTTTCCCT | 7 | 185 | 6.6 |  |  |  |
|  |  |  | TTGATGCTACAATGAAGGAA--------GCTTTTCCCT | 8 | 166 | 5.9 |  |  |  |
|  |  |  | TTGATGCTACAATGAAGGA----GCAAAGCTTTTCCCT | 4 | 157 | 5.6 |  |  |  |
|  | 2 | 1 | TTGATGCTACAATGAAG------------CTTTTCCCT | 12 | 974 | 10.9 | 18.9 | 20 | 8 |
|  |  |  | TTGATGCTACAATGAAGGAAC----GAAGCTTTTCCCT | 4 | 648 | 7.3 |  |  |  |
|  |  |  | TTGATGCTACAATGAAGGAA--------GCTTTTCCTT | 8 | 608 | 6.8 |  |  |  |
|  |  |  | TTGATGCTACAATGA-------------GCTTTTCCCT | 13 | 515 | 5.8 |  |  |  |
|  |  | 2 | TTGATGCTACAATGAAG------------CTTTTCCCT | 12 | 657 | 11.7 | 14.2 |  |  |
|  |  |  | TTGATGCTACAATGATGGAC-------AGCTTTTCCCT | 7 | 393 | 7.0 |  |  |  |
|  |  |  | TTGATGCTACAATGG-------------GCTTTTCCCT | 13 | 344 | 6.1 |  |  |  |
|  |  |  | TTGATGCTACAATGAAGGAA--------GCTTTTCCCT | 8 | 342 | 6.1 |  |  |  |
|  |  | 3 | TTGATGCTACAATGAAG------------CTTTTCCCT | 12 | 3014 | 17.8 | 25.5 |  |  |
|  |  |  | TTGATGCTACAATGA-------------GCTTTTCCCT | 13 | 1979 | 11.7 |  |  |  |
|  |  |  | TTGATGCTACAATGAAGGAAC----GAAGCTTTTCCCT | 4 | 1959 | 11.5 |  |  |  |
|  |  |  | TTGATGCTACAATGAAGGAA--------GCTTTTCCCT | 8 | 1472 | 8.7 |  |  |  |
|  |  | 4 | TTGATGCTACAATGAAG------------CTTTTCCCT | 12 | 831 | 13.8 | 19.6 |  |  |
|  |  |  | TTGATGCTACAATTAAGGAAC----GAAGCTTTTCCCT | 4 | 476 | 7.9 |  |  |  |
|  |  |  | TTGATGCTACAATGA-------------GCTTTTCCCT | 13 | 475 | 7.9 |  |  |  |
|  |  |  | TTGATGCTACAATGAAGGAA--------GCTTTTTCCT | 8 | 399 | 6.6 |  |  |  |
|  |  | 5 | TTGATGCTACAATGAAG------------CTTTTCCCT | 12 | 2190 | 13.2 | 31.7 |  |  |
|  |  |  | TTGATGCTACAATGA-------------GCTTTTCCCT | 13 | 1335 | 8.0 |  |  |  |
|  |  |  | TTGATGCTACAATGAAGGAAC----GAAGCTTTTCCCT | 4 | 1128 | 6.8 |  |  |  |
|  |  |  | TTGATGCTACAATGAAGGAA-------AGCTTTTACCT | 7 | 1078 | 6.5 |  |  |  |
|  |  | 6 | TTGATGCTACAATGAAG------------CTTTTCCCT | 12 | 553 | 14.0 | 11.4 |  |  |
|  |  |  | TTGATGCTACAATGAAGCAA-------AACTTTTCCCT | 7 | 330 | 8.4 |  |  |  |
|  |  |  | TTGATGCTACAATGAAGGAAC----GAAGCTTTTCCCT | 4 | 255 | 6.5 |  |  |  |
|  |  |  | TTGATGCTACAATGAAGGAA--------GCTTTTCCCT | 8 | 225 | 5.7 |  |  |  |
|  |  | 7 | TTGATGCTACAATGAAG------------CTTTTCCCT | 12 | 351 | 8.3 | 11.5 |  |  |
|  |  |  | TTGATGCTACAATGAACGAA-------AGCTTTTCCCT | 7 | 244 | 5.7 |  |  |  |
|  |  |  | TTGATGCTACAATGAAGGAA--------GCTTTTCCCT | 8 | 244 | 5.7 |  |  |  |
|  |  |  | -------------------------------------- | 67 | 218 | 5.1 |  |  |  |
|  |  | 8 | TTGATGCTACAATGAAG------------CTTTTCCCT | 12 | 1085 | 12.3 | 29.6 |  |  |
|  |  |  | TTGATGCTACAATGA-------------GCTTTTCCCT | 13 | 575 | 6.5 |  |  |  |
|  |  |  | TTGATGCTACAATGAAGGAAC----GAATCTTTTCCCT | 4 | 555 | 6.3 |  |  |  |
|  |  |  | TTGATGCTACAATGAAGGAA-------AGCTTTTCCCT | 7 | 516 | 5.9 |  |  |  |

*Bold red letters indicate TAM, and underlined letters denote reRNA target sequences

**Supplementary Table 18.** Percentage of progeny seedlings from parental plants infected with TRV2 with $eTnpBc_{2xSV40nls}$ exhibiting a white photobleaching phenotype

| Construct | Progeny seeeds from | # of green seedlings | # of white seedlings | Total # of seedlings | % white seedlings | Average % of white seedlings | Standard Deviation |
|---|---|---|---|---|---|---|---|
| $eTnpBc_{2xSV40nls}$-$reRNA4^{PDS}$ | Pods from the top 1/3rd of the plant | 23 | 83 | 106 | 78 | 55 | 16 |
| | | 80 | 78 | 158 | 49 | | |
| | | 79 | 66 | 145 | 46 | | |
| | | 42 | 38 | 80 | 48 | | |
| $eTnpBc_{2xSV40nls}$-$^{HH}reRNA4^{PDS}$ | Pods from the top 1/3rd of the plant | 17 | 65 | 82 | 79 | 68 | 15 |
| | | 18 | 55 | 73 | 75 | | |
| | | 29 | 30 | 59 | 51 | | |
| | | 14 | 70 | 84 | 83 | | |
| | | 29 | 33 | 62 | 53 | | |

**Supplementary Table 19.** Indel types and frequencies in *ChlH* of progenies from parental plants infected with TRV2 expressing eTnpBc variant and reRNA$^{ChlH}$

| Construct | Parent # | Plant # | Most common indels | n_deleted | Reads | % Edited reads | Overall editing frequency | Average editing frequency | Standard deviation |
|---|---|---|---|---|---|---|---|---|---|
| | | | GTCAACTGCTCTGGGGTGTTCAGAGATCTCTTC**ATCAA*** | Wild-type | | | | | |
| eTnpBc-reRNA$^{ChlH}$ | 1 | 1 | GTCAACTG---------GTTCAGAGATCTCTTCATCAA | 9 | 842 | 22.9 | 99.6 | 100 | 0 |
|  |  |  | GTCAACT-------------CAGAGATCTCTTCATCAA | 13 | 823 | 22.4 |  |  |  |
|  |  |  | GTCAACTGCTCTGG-----TCAGAGATCTCTTCATCAA | 5 | 697 | 19.0 |  |  |  |
|  |  |  | GTCAACTGCTCT-----GTTCAGAGATCTCTTCATCAA | 5 | 654 | 17.8 |  |  |  |
|  |  | 2 | GTCAACT-------------CAGAGATCTCTTCATCAA | 13 | 1116 | 23.5 | 99.8 |  |  |
|  |  |  | GTCAACTG---------GTTCAGAGATCTCTTCATCAA | 9 | 939 | 19.8 |  |  |  |
|  |  |  | GTCAACTGCTCT-----GTTCAGAGATCTCTTCATCAA | 5 | 930 | 19.6 |  |  |  |
|  |  |  | GTCAACTGCTCTGG-----TCAGAGATCTCTTCATCAA | 5 | 864 | 18.2 |  |  |  |
|  |  | 3 | GTCAACT-------------CAGAGATCTCTTCATCAA | 13 | 1872 | 47.1 | 99.8 |  |  |
|  |  |  | GTCAACTGCTCTGG-----TCAGAGATCTCTTCATCAA | 5 | 748 | 18.8 |  |  |  |
|  |  |  | GTCAACTG---------GTTCAGAGATCTCTTCATCAA | 9 | 743 | 18.7 |  |  |  |
|  |  | 4 | GTCAACT-------------CAGAGATCTCTTCATCAA | 13 | 1426 | 45.2 | 99.2 |  |  |
|  |  |  | GTCAACTG---------GTTCAGAGATCTCTTCATCAA | 9 | 621 | 19.7 |  |  |  |
|  |  |  | GTCAACTGCTCTGG-----TCAGAGATCTCTTCATCAA | 5 | 555 | 17.6 |  |  |  |
|  |  | 5 | GTCAACT-------------CAGAGATCTCTTCATCAA | 13 | 1164 | 45.2 | 99.3 |  |  |
|  |  |  | GTCAACTG---------GTTCAGAGATCTCTTCATCAA | 9 | 482 | 18.7 |  |  |  |
|  |  |  | GTCAACTGCTCTGG-----TCAGAGATCTCTTCATCAA | 5 | 430 | 16.7 |  |  |  |
|  |  |  | GTCAACTGCTCT-----GTTCAGAGATCTCTTCATCAA | 5 | 70 | 2.7 |  |  |  |
|  |  | 6 | GTCAACTG---------GTTCAGAGATCTCTTCATCAA | 9 | 1507 | 38.2 | 99.6 |  |  |
|  |  |  | GTCAACT-------------CAGAGATCTCTTCATCAA | 13 | 922 | 23.4 |  |  |  |
|  |  |  | GTCAACTGCTCT-----GTTCAGAGATCTCTTCATCAA | 5 | 817 | 20.7 |  |  |  |
|  |  | 7 | GTCAACT-------------CAGAGATCTCTTCATCAA | 13 | 1301 | 45.0 | 99.6 |  |  |
|  |  |  | GTCAACTG---------GTTCAGAGATCTCTTCATCAA | 9 | 579 | 20.0 |  |  |  |
|  |  |  | GTCAACTGCTCTGG-----TCAGAGATCTCTTCATCAA | 5 | 541 | 18.7 |  |  |  |
|  |  | 8 | GTCAACTGCTCT-----GTTCAGAGATCTCTTCATCAA | 5 | 1636 | 41.4 | 99.6 |  |  |
|  |  |  | GTCAACTG---------GTTCAGAGATCTCTTCATCAA | 9 | 1565 | 39.6 |  |  |  |
|  |  |  | GTCAACTGCTCT-----GTTCAGAGATCTCTTCATCAA | 5 | 62 | 1.6 |  |  |  |
|  | 2 | 1 | GTCAACTG---------GTTCAGAGATCTCTTCATCAA | 9 | 434 | 36.2 | 98.5 | 99 | 0 |
|  |  |  | GTCAACT-------------CAGAGATCTCTTCATCAA | 13 | 317 | 26.5 |  |  |  |
|  |  |  | GTCAACTGCTCT-----GTTCAGAGATCTCTTCATCAA | 5 | 229 | 19.1 |  |  |  |
|  |  |  | GTCAACTGCTCTGG-----TCAGAGATCTCTTCATCAA | 5 | 33 | 2.8 |  |  |  |
|  |  | 2 | GTCAACTG---------GTTCAGAGATCTCTTCATCAA | 9 | 390 | 29.8 | 98.5 |  |  |
|  |  |  | GTCAACT-------------CAGAGATCTCTTCATCAA | 13 | 313 | 23.9 |  |  |  |
|  |  |  | GTCAACTGCTCT-----GTTCAGAGATCTCTTCATCAA | 5 | 275 | 21.0 |  |  |  |
|  |  |  | GTCAACTGCTCT--------CAGAGATCTCTTCATCAA | 8 | 35 | 2.7 |  |  |  |
|  |  |  | GTCAACTGCTCTGG-----TCAGAGATCTCTTCATCAA | 5 | 27 | 2.1 |  |  |  |
|  |  | 3 | GTCAACT-------------CAGAGATCTCTTCATCAA | 13 | 249 | 25.6 | 99.0 |  |  |
|  |  |  | GTCAACTG---------GTTCAGAGATCTCTTCATCAA | 9 | 199 | 20.5 |  |  |  |
|  |  |  | GTCAACTGCTCT-----GTTCAGAGATCTCTTCATCAA | 5 | 198 | 20.4 |  |  |  |
|  |  |  | GTCAACTGCTCTGG-----TCAGAGATCTCTTCATCAA | 5 | 156 | 16.0 |  |  |  |
|  |  | 4 | GTCAACTGCTCTGG-----TCAGAGATCTCTTCATCAA | 5 | 475 | 34.5 | 97.9 |  |  |
|  |  |  | GTCAACT-------------CAGAGATCTCTTCATCAA | 13 | 304 | 22.1 |  |  |  |
|  |  |  | GTCAACTGCTCT-----GTTCAGAGATCTCTTCATCAA | 5 | 288 | 20.9 |  |  |  |
|  |  |  | GTCAACTG---------GTTCAGAGATCTCTTCATCAA | 9 | 45 | 3.3 |  |  |  |
|  |  | 5 | GTCAACTGCTCT-----GTTCAGAGATCTCTTCATCAA | 5 | 746 | 44.3 | 99.1 |  |  |
|  |  |  | GTCAACTG---------GTTCAGAGATCTCTTCATCAA | 9 | 307 | 18.2 |  |  |  |
|  |  |  | GTCAACTGCTCTGG-----TCAGAGATCTCTTCATCAA | 5 | 257 | 15.3 |  |  |  |
|  |  |  | GTCAACT-------------CAGAGATCTCTTCATCAA | 13 | 46 | 2.7 |  |  |  |
|  |  | 6 | GTCAACTGCTCT-----GTTCAGAGATCTCTTCATCAA | 5 | 653 | 42.8 | 99.1 |  |  |
|  |  |  | GTCAACTG---------GTTCAGAGATCTCTTCATCAA | 9 | 323 | 21.2 |  |  |  |
|  |  |  | GTCAACTGCTCTGG-----TCAGAGATCTCTTCATCAA | 5 | 197 | 12.9 |  |  |  |
|  |  |  | GTCAACTGCTCT-----GTTCAGAGATCTCTTCATCAA | 5 | 32 | 2.1 |  |  |  |
|  |  | 7 | GTCAACT-------------CAGAGATCTCTTCATCAA | 13 | 331 | 25.7 | 98.9 |  |  |
|  |  |  | GTCAACTGCTCT-----GTTCAGAGATCTCTTCATCAA | 5 | 221 | 17.1 |  |  |  |
|  |  |  | GTCAACTG---------GTTCAGAGATCTCTTCATCAA | 9 | 188 | 14.6 |  |  |  |
|  |  |  | GTCAACTGCTCTGG-----TCAGAGATCTCTTCATCAA | 5 | 183 | 14.2 |  |  |  |
|  |  | 8 | GTCAACT-------------CAGAGATCTCTTCATCAA | 13 | 301 | 22.7 | 99.3 |  |  |
|  |  |  | GTCAACTGCTCT-----GTTCAGAGATCTCTTCATCAA | 5 | 282 | 21.3 |  |  |  |
|  |  |  | GTCAACTG---------GTTCAGAGATCTCTTCATCAA | 9 | 252 | 19.0 |  |  |  |
|  |  |  | GTCAACTGCTCTGG-----TCAGAGATCTCTTCATCAA | 5 | 220 | 16.6 |  |  |  |

*Bold red letters indicate TAM, and underlined letters denote reRNA target sequences

**Supplementary Table 20.** Indel types and frequencies in two *ChlH* genes of progenies from parental plants infected with TRV2 expressing eTnpBc variant and reRNA4$^{ChlH}$

| Parent # | Gene ID | Plant # | Most common indels | n_deleted | Reads | % Edited reads | Overall editing frequency | Average editing frequency | Standard deviation |
|---|---|---|---|---|---|---|---|---|---|
| 1 | *ChlH1* -Nbe07g00760 | | GTCAACTGCTCTGGGGTGTTCAGAGATCTCTTCATCAA* | Wild-type | | | | | |
|  |  | 1 | GTCAACTG---------GTTCAGAGATCTCTTCATCAA | 9 | 937 | 51.1 | 99.4 | 99 | 0 |
|  |  |  | GTCAACTGCTCTGG-----TCAGAGATCTCTTCATCAA | 5 | 804 | 43.9 |  |  |  |
|  |  |  | GTCAACT-------------CAGAGATCTCTTCATCAA | 13 | 34 | 1.9 |  |  |  |
|  |  |  | GTCAACTGCTCT-----GATCAGAGATCTCTTCATCAA | 5 | 32 | 1.7 |  |  |  |
|  |  | 2 | GTCAACTG---------GTTCAGAGATCTCTTCATCAA | 9 | 1085 | 49.1 | 99.6 |  |  |
|  |  |  | GTCAACTGCTCTGG-----TCAGAGATCTCTTCATCAA | 5 | 992 | 44.9 |  |  |  |
|  |  |  | GTCAACTGCTCT-----GTTCAGAGATCTCTTCATCAA | 5 | 56 | 2.5 |  |  |  |
|  |  |  | GTCAACT-------------CAGAGATCTCTTCATCAA | 13 | 44 | 2.0 |  |  |  |
|  |  | 3 | GTCAACTGCTCTGG-----TCAGAGATCTCTTCATCAA | 5 | 846 | 47.4 | 100.0 |  |  |
|  |  |  | GTCAACTG---------GTTCAGAGATCTCTTCATCAA | 9 | 845 | 47.4 |  |  |  |
|  |  |  | GTCAACT-------------CAGAGATCTCTTCATCAA | 13 | 53 | 3.0 |  |  |  |
|  |  | 4 | GTCAACTG---------GTTCAGAGATCTCTTCATCAA | 9 | 710 | 49.7 | 98.8 |  |  |
|  |  |  | GTCAACTGCTCTGG-----TCAGAGATCTCTTCATCAA | 5 | 641 | 44.9 |  |  |  |
|  |  |  | GTCAACT-------------CAGAGATCTCTTCATCAA | 13 | 37 | 2.6 |  |  |  |
|  |  | 5 | GTCAACTG---------GTTCAGAGATCTCTTCATCAA | 9 | 562 | 50.5 | 98.7 |  |  |
|  |  |  | GTCAACTGCTCTGG-----TCAGAGATCTCTTCATCAA | 5 | 499 | 44.9 |  |  |  |
|  |  |  | GTCAACT-------------CAGAGATCTCTTCATCAA | 13 | 21 | 1.9 |  |  |  |
|  |  | 6 | GTCAACTG---------GTTCAGAGATCTCTTCATCAA | 9 | 1738 | 94.1 | 99.1 |  |  |
|  |  |  | GTCAACT-------------CAGAGATCTCTTCATCAA | 13 | 37 | 2.0 |  |  |  |
|  |  |  | GTCAACTGCCCT-----GTTCAGAGATCTCTTCATCAA | 5 | 19 | 1.0 |  |  |  |
|  |  | 7 | GTCAACTG---------GTTCAGAGATCTCTTCATCAA | 9 | 653 | 49.2 | 99.1 |  |  |
|  |  |  | GTCAACTGCTCTGG-----TCAGAGATCTCTTCATCAA | 5 | 617 | 46.5 |  |  |  |
|  |  |  | GTCAACT-------------CAGAGATCTCTTCATCAA | 13 | 26 | 2.0 |  |  |  |
|  |  | 8 | GTCAACTG---------GTTCAGAGATCTCTTCATCAA | 9 | 1805 | 94.9 | 99.6 |  |  |
|  |  |  | GTCAACTGCTCT-----GTTCAGAGATCTCTTCATCAA | 5 | 56 | 2.9 |  |  |  |
| | *ChlH2* -Nbe08g21240 | | GTCAACTGCTCTGGGGTGTTCAGAGATCTCTTCATCAA* | Wild-type | | | | | |
|  |  | 1 | GTCAACT-------------CAGAGATCTCTTCATCAA | 13 | 944 | 51.3 | 99.8 | 100 | 0 |
|  |  |  | GTCAACTGCTCT-----GTTCAGAGATCTCTTCATCAA | 5 | 776 | 42.2 |  |  |  |
|  |  |  | GTCAACTGCTCTGG-----TCAGAGATCTCTTCATCAA | 5 | 33 | 1.8 |  |  |  |
|  |  |  | GTCAACTG---------GTTCAGAGATCTCTTCATCAA | 9 | 27 | 1.5 |  |  |  |
|  |  |  | GTCAACTGC--------------AGATCTCTTCATCAA | 14 | 24 | 1.3 |  |  |  |
|  |  | 2 | GTCAACT-------------CAGAGATCTCTTCATCAA | 13 | 1281 | 50.5 | 100.0 |  |  |
|  |  |  | GTCAACTGCTCT-----GTTCAGAGATCTCTTCATCAA | 5 | 1132 | 44.6 |  |  |  |
|  |  |  | GTCGACTGCTCTGG-----TCAGAGATCTCTTCATCAA | 5 | 38 | 1.5 |  |  |  |
|  |  | 3 | GTCAACT-------------CAGAGATCTCTTCATCAA | 13 | 2095 | 95.7 | 100.0 |  |  |
|  |  |  | GTCAACTGCTCTGG-----TCAGAGATCTCTTCATCAA | 5 | 25 | 1.1 |  |  |  |
|  |  | 4 | GTCAACT-------------CAGAGATCTCTTCATCAA | 13 | 1619 | 93.7 | 99.6 |  |  |
|  |  |  | GTCAACTGCTCT-----GTTCAGAGATCTCTTCATCAA | 5 | 21 | 1.2 |  |  |  |
|  |  |  | GTCAACTGC--------------AGATCTCTTCATCAA | 14 | 21 | 1.2 |  |  |  |
|  |  | 5 | GTCAACT-------------CAGAGATCTCTTCATCAA | 13 | 1311 | 89.6 | 99.9 |  |  |
|  |  |  | GTCAACTGCTCT-----GTTCAGAGATCTCTTCATCAA | 5 | 99 | 6.8 |  |  |  |
|  |  | 6 | GTCAACT-------------CAGAGATCTCTTCATCAA | 13 | 1040 | 49.5 | 100.0 |  |  |
|  |  |  | GTCAACTGCTCT-----GTTCAGAGATCTCTTCATCAA | 5 | 949 | 45.2 |  |  |  |
|  |  |  | GTCAACTG---------GTTCAGAGATCTCTTCATCAA | 9 | 48 | 2.3 |  |  |  |
|  |  |  | GTCAACTGCTCT--------CAGAGATCTCTTCATCAA | 8 | 27 | 1.3 |  |  |  |
|  |  | 7 | GTCAACT-------------CAGAGATCTCTTCATCAA | 13 | 1464 | 93.8 | 99.9 |  |  |
|  |  |  | GTCAACTGCTCT-----GTTCAGAGATCTCTTCATCAA | 5 | 22 | 1.4 |  |  |  |
|  |  | 8 | GTCAACTGCTCT-----GTTCAGAGATCTCTTCATCAA | 5 | 1895 | 92.6 | 99.6 |  |  |
|  |  |  | GTCAACT-------------CAGAGATCTCTTCATCAA | 13 | 54 | 2.6 |  |  |  |
|  |  |  | GTCAACTG---------GTTCAGAGATCTCTTCATCAA | 9 | 48 | 2.3 |  |  |  |
| 2 | *ChlH1* -Nbe07g00760 | | GTCAACTGCTCTGGGGTGTTCAGAGATCTCTTCATCAA* | Wild-type | | | | | |
|  |  | 1 | GTCAACTG---------GTTCAGAGATCTCTTCATCAA | 9 | 485 | 87.4 | 97.8 | 98 | 1 |
|  |  |  | GTCAACTGCTCTGG-----TCAGAGATCTCTTCATCAA | 5 | 41 | 7.4 |  |  |  |
|  |  |  | GTCAACT-------------CAGAGATCCCTTCATCAA | 13 | 8 | 1.4 |  |  |  |
|  |  | 2 | GTCAACTG---------GTTCAGAGATCTCTTCATCAA | 9 | 435 | 78.7 | 96.8 |  |  |
|  |  |  | GTCAACTGCTCTGG-----TCAGAGATCTCTTCATCAA | 5 | 40 | 7.2 |  |  |  |
|  |  |  | GTCAACT-------------CAGAGATCTCTTCATCAA | 13 | 28 | 5.1 |  |  |  |
|  |  | 3 | GTCAACTG---------GTTCAGAGATCTCTTCATCAA | 9 | 215 | 50.9 | 97.9 |  |  |
|  |  |  | GTCAACTGCTCTGG-----TCAGAGATCTCTTCATCAA | 5 | 179 | 42.4 |  |  |  |
|  |  |  | GTCAACTGCTCT-----GATCAGAGATCTCTTCATCAA | 5 | 8 | 1.9 |  |  |  |
|  |  | 4 | GTCAACTGCTCTGG-----TCAGAGATCTCTTCATCAA | 5 | 540 | 83.9 | 97.5 |  |  |
|  |  |  | GTCAACTG---------GTTCAGAGATCTCTTCATCAA | 9 | 52 | 8.1 |  |  |  |
|  |  |  | GTCAACTGCTCT-----GTTCAGAGATCTCTTCATCAA | 5 | 17 | 2.6 |  |  |  |
|  |  | 5 | GTCAACTG---------GTTCAGAGATCTCTTCATCAA | 9 | 352 | 50.4 | 98.3 |  |  |
|  |  |  | GTCAACTGCTCTGG-----TCAGAGATCTCTTCATCAA | 5 | 298 | 42.6 |  |  |  |
|  |  |  | GTCAACTGCTCT-----GTTCAGAGATCTCTTCATCAA | 5 | 28 | 4.0 |  |  |  |
|  |  | 6 | GTCAACTG---------GTTCAGAGATCTCTTCATCAA | 9 | 364 | 54.6 | 98.7 |  |  |
|  |  |  | GTCAACTGCTCTGG-----TCAGAGATCTCTTCATCAA | 5 | 243 | 36.4 |  |  |  |
|  |  |  | GTCAACTGCTCT-----GTTCAGAGATCTCTTCATCAA | 5 | 36 | 5.4 |  |  |  |

| | | | | | | | | | |
|---|---|---|---|---|---|---|---|---|---|
|  |  | 7 | GTCAACTG---------GTTCAGAGATCTCTTCATCAA | 9 | 247 | 45.7 | 98.2 |  |  |
|  |  |  | GTCAACTGCTCTGG-----TCAGAGATCTCTTCATCAA | 5 | 241 | 44.6 |  |  |  |
|  |  |  | GTCAACT-------------CAGAGATCTCTTCATCAA | 13 | 20 | 3.7 |  |  |  |
|  |  | 8 | GTCAACTG---------GTTCAGAGATCTCTTCATCAA | 9 | 292 | 48.1 | 99.0 |  |  |
|  |  |  | GTCAACTGCTCTGG-----TCAGAGATCTCTTCATCAA | 5 | 262 | 43.2 |  |  |  |
|  |  |  | GTCAACTGCTCT-----GTTCAGAGATCTCTTCATCAA | 5 | 16 | 2.6 |  |  |  |
| | *ChlH2* -Nbe08g21240 | | GTCAACTGCTCTGGGGTGTTCAGAGATCTCTTC**ATCAA*** | Wild-type | | | | | |
|  |  | 1 | GTCAACT-------------CAGAGATCTCTTCATCAA | 13 | 352 | 54.7 | 99.1 | 99 | 1 |
|  |  |  | GTCAACTGCTCT-----GTTCAGAGATCTCTTCATCAA | 5 | 258 | 40.1 |  |  |  |
|  |  |  | GTCAACTG---------GTTCAGAGATCTCTTCATCAA | 9 | 7 | 1.1 |  |  |  |
|  |  | 2 | GTCAACT-------------CAGAGATCTCTTCATCAA | 13 | 359 | 47.5 | 99.7 |  |  |
|  |  |  | GTCAACTGCTCT-----GTTCAGAGATCTCTTCATCAA | 5 | 317 | 42.0 |  |  |  |
|  |  |  | GTCAACTGCTCT--------CAGAGATCTCTTCATCAA | 8 | 41 | 5.4 |  |  |  |
|  |  | 3 | GTCAACT-------------CAGAGATCTCTTCATCAA | 13 | 283 | 51.5 | 99.8 |  |  |
|  |  |  | GTCAACTGCTCT-----GTTCAGAGATCTCTTCATCAA | 5 | 229 | 41.6 |  |  |  |
|  |  |  | GTCAACTGCTCTGG-----TCAGAGATCTATTCATCAA | 5 | 14 | 2.5 |  |  |  |
|  |  | 4 | GTCAACT-------------CAGAGATCTCTTCATCAA | 13 | 341 | 46.6 | 98.2 |  |  |
|  |  |  | GTCAACTGCTCT-----GTTCAGAGATCTCTTCATCAA | 5 | 335 | 45.8 |  |  |  |
|  |  |  | GTCAACTGCTCTGG-----TCAGAGATCTCTTCATCAA | 5 | 20 | 2.7 |  |  |  |
|  |  |  | GTCAACTGCTCT--------CAGAGATCTCTTCATCAA | 8 | 15 | 2.0 |  |  |  |
|  |  | 5 | GTCAACTGCTCT-----GTTCAGAGATCTCTTCATCAA | 5 | 879 | 89.3 | 99.6 |  |  |
|  |  |  | GTCAACT-------------CAGAGATCTCTTCATCAA | 13 | 61 | 6.2 |  |  |  |
|  |  |  | GTCAACTGCTCTGG-----TCAGAGATCTCTTCATCAA | 5 | 16 | 1.6 |  |  |  |
|  |  | 6 | GTCAACTGCTCT-----GTTCAGAGATCTCTTCATCAA | 5 | 753 | 87.9 | 99.4 |  |  |
|  |  |  | GTCAACT-------------CAGAGATCTCTTCATCAA | 13 | 36 | 4.2 |  |  |  |
|  |  |  | GTCAACTGCTCT--------CAGAGATCTCTTCATCAA | 8 | 22 | 2.6 |  |  |  |
|  |  | 7 | GTCAACT-------------CAGAGATCTCTTCATCAA | 13 | 397 | 53.0 | 99.5 |  |  |
|  |  |  | GTCAACTGCTCT-----GTTCAGAGATCTCTTCATCAA | 5 | 290 | 38.7 |  |  |  |
|  |  |  | GTCAACTGC--------------AGATCTCTTCATCAA | 14 | 15 | 2.0 |  |  |  |
|  |  | 8 | GTCAACTGCTCT-----GTTCAGAGATCTCTTCATCAA | 5 | 340 | 47.3 | 99.6 |  |  |
|  |  |  | GTCAACT-------------CAGAGATCTCTTCATCAA | 13 | 334 | 46.5 |  |  |  |
|  |  |  | GTCAACTGCTCT--------CAGAGATCTCTTCATCAA | 8 | 16 | 2.2 |  |  |  |

*Bold red letters indicate TAM, and underlined letters denote reRNA target sequences

**Supplementary Table 21.** No editing at the off-target sites of reRNA4[PDS]

| Off Taget # | | Sequence sourrounding taget site* | Indel % |
|---|---|---|---|
| PDS4.1off | Target | CTTTCACAGCAGACAACTTT**GACATTTCCTTTATTGAAGCATCAA**AATGA | |
| | Wild-type | CTTTCACAGCAGACAACTTTGACATTTCCTTTATTGAAGCATCAAAATGA | |
| | Plant #1 | CTTTCACAGCAGACAACTTTGACATTTCCTTTATTGAAGCATCAAAATGA | 0 |
| | Plant #2 | CTTTCACAGCAGACAACTTTGACATTTCCTTTATTGAAGCATCAAAATGA | 0 |
| | Plant #3 | CTTTCACAGCAGACAACTTTGACATTTCCTTTATTGAAGCATCAAAATGA | 0 |
| | Plant #4 | CTTTCACAGCAGACAACTTTGACATTTCCTTTATTGAAGCATCAAAATGA | 0 |
| | Plant #5 | CTTTCACAGCAGACAACTTTGACATTTCCTTTATTGAAGCATCAAAATGA | 0 |
| PDS4.2off | Target | TGAGT**TTGATGCAACTGTAAAGGCACTAGC**GCCAATTGAGGCCGAGTTCG | |
| | Wild-type | TGAGTTTGATGCAACTGTAAAGGCACTAGCGCCAATTGAGGCCGAGTTCG | |
| | Plant #1 | TGAGTTTGATGCAACTGTAAAGGCACTAGCGCCAATTGAGGCCGAGTTCG | 0 |
| | Plant #2 | TGAGTTTGATGCAACTGTAAAGGCACTAGCGCCAATTGAGGCCGAGTTCG | 0 |
| | Plant #3 | TGAGTTTGATGCAACTGTAAAGGCACTAGCGCCAATTGAGGCCGAGTTCG | 0 |
| | Plant #4 | TGAGTTTGATGCAACTGTAAAGGCACTAGCGCCAATTGAGGCCGAGTTCG | 0 |
| | Plant #5 | TGAGTTTGATGCAACTGTAAAGGCACTAGCGCCAATTGAGGCCGAGTTCG | 0 |
| PDS4.3off | Target | TCCCC**TTGATGCTACACTGATGAAACTTCC**AAATTGTGTCCAATCAATTA | |
| | Wild-type | TCCCCTTGATGCTACACTGATGAAACTTCCAAATTGTGTCCAATCAATTA | |
| | Plant #1 | TCCCCTTGATGCTACACTGATGAAACTTCCAAATTGTGTCCAATCAATTA | 0 |
| | Plant #2 | TCCCCTTGATGCTACACTGATGAAACTTCCAAATTGTGTCCAATCAATTA | 0 |
| | Plant #3 | TCCCCTTGATGCTACACTGATGAAACTTCCAAATTGTGTCCAATCAATTA | 0 |
| | Plant #4 | TCCCCTTGATGCTACACTGATGAAACTTCCAAATTGTGTCCAATCAATTA | 0 |
| | Plant #5 | TCCCCTTGATGCTACACTGATGAAACTTCCAAATTGTGTCCAATCAATTA | 0 |
| PDS4.4off | Target | ATCAG**TTGATGCTAGTATGAGAGAACCAGC**AGAGAGGGGGCGAGCAATCT | |
| | Wild-type** | ATCAGTTGATGCTAGTcTGAGAGAACCAGCAGAGAGGGGGCcAaCAATCT | |
| | Plant #1** | ATCAGTTGATGCTAGTcTGAGAGAACCAGCAGAGAGGGGGCcAaCAATCT | 0 |
| | Plant #2** | ATCAGTTGATGCTAGTcTGAGAGAACCAGCAGAGAGGGGGCcAaCAATCT | 0 |
| | Plant #3** | ATCAGTTGATGCTAGTcTGAGAGAACCAGCAGAGAGGGGGCcAaCAATCT | 0 |
| | Plant #4** | ATCAGTTGATGCTAGTcTGAGAGAACCAGCAGAGAGGGGGCcAaCAATCT | 0 |
| | Plant #5** | ATCAGTTGATGCTAGTcTGAGAGAACCAGCAGAGAGGGGGCcAaCAATCT | 0 |
| PDS4.5off | Target | TGCTT**TTGATGCTGCAATTAAAGAAGTAAC**TGTATATTATTGTTATTCCC | |
| | Wild-type | TGCTTTTGATGCTGCAATTAAAGAAGTAACTGTATATTATTGTTATTCCC | |
| | Plant #1 | TGCTTTTGATGCTGCAATTAAAGAAGTAACTGTATATTATTGTTATTCCC | 0 |
| | Plant #2 | TGCTTTTGATGCTGCAATTAAAGAAGTAACTGTATATTATTGTTATTCCC | 0 |
| | Plant #3 | TGCTTTTGATGCTGCAATTAAAGAAGTAACTGTATATTATTGTTATTCCC | 0 |
| | Plant #4 | TGCTTTTGATGCTGCAATTAAAGAAGTAACTGTATATTATTGTTATTCCC | 0 |
| | Plant #5 | TGCTTTTGATGCTGCAATTAAAGAAGTAACTGTATATTATTGTTATTCCC | 0 |
| PDS4.6off | Target | TCTATACATCAAGCACCAGA**GATAATTCCTACAGAGTAGCATCAA**TTGCT | |
| | Wild-type | TCTATACATCAAGCACCAGAGATAATTCCTACAGAGTAGCATCAATTGCT | |
| | Plant #1 | TCTATACATCAAGCACCAGAGATAATTCCTACAGAGTAGCATCAATTGCT | 0 |
| | Plant #2 | TCTATACATCAAGCACCAGAGATAATTCCTACAGAGTAGCATCAATTGCT | 0 |
| | Plant #3 | TCTATACATCAAGCACCAGAGATAATTCCTACAGAGTAGCATCAATTGCT | 0 |
| | Plant #4 | TCTATACATCAAGCACCAGAGATAATTCCTACAGAGTAGCATCAATTGCT | 0 |
| | Plant #5 | TCTATACATCAAGCACCAGAGATAATTCCTACAGAGTAGCATCAATTGCT | 0 |
| PDS4.7off | Target | TGGTGAATGTTACGAACCGA**GTTAGTTTCGTCATTGCAACATCAA**CCGTT | |
| | Wild-type | TGGTGAATGTTACGAACCGAGTTAGTTTCGTCATTGCAACATCAACCGTT | |
| | Plant #1 | TGGTGAATGTTACGAACCGAGTTAGTTTCGTCATTGCAACATCAACCGTT | 0 |
| | Plant #2 | TGGTGAATGTTACGAACCGAGTTAGTTTCGTCATTGCAACATCAACCGTT | 0 |
| | Plant #3 | TGGTGAATGTTACGAACCGAGTTAGTTTCGTCATTGCAACATCAACCGTT | 0 |
| | Plant #4 | TGGTGAATGTTACGAACCGAGTTAGTTTCGTCATTGCAACATCAACCGTT | 0 |
| | Plant #5 | TGGTGAATGTTACGAACCGAGTTAGTTTCGTCATTGCAACATCAACCGTT | 0 |

*Bold red letters indicate TAM, and bold underlined letters denote reRNA target sequence
**Lower case letters indiate changes observed compared to the deposited sequence

**Supplementary Table 22.** No editing at the off-target sites of reRNA4$^{ChlH}$

| Off Taget # | | Sequence sourrounding taget site* | Indel % |
|---|---|---|---|
| ChlH.1off | Target | CTAACATCATGTCCGACAAC**GGGTGATCAAAGATCTCAAT****ATCAA**TTGCA | |
| | Wild-type | CTAACATCATGTCCGACAACGGGTGATCAAAGATCTCAATATCAATTGCA | |
| | Plant #1 | CTAACATCATGTCCGACAACGGGTGATCAAAGATCTCAATATCAATTGCA | 0 |
| | Plant #2 | CTAACATCATGTCCGACAACGGGTGATCAAAGATCTCAATATCAATTGCA | 0 |
| | Plant #3 | CTAACATCATGTCCGACAACGGGTGATCAAAGATCTCAATATCAATTGCA | 0 |
| | Plant #4 | CTAACATCATGTCCGACAACGGGTGATCAAAGATCTCAATATCAATTGCA | 0 |
| | Plant #5 | CTAACATCATGTCCGACAACGGGTGATCAAAGATCTCAATATCAATTGCA | 0 |
| ChlH.2off | Target | GTAAT**TTGAT****GAAGACAGATCTGAAAACCA**CACGAAAATTAAAAATGGTC | |
| | Wild-type | GTAATTTGATGAAGACAGATCTGAAAACCACACGAAAATTAAAAATGGTC | |
| | Plant #1 | GTAATTTGATGAAGACAGATCTGAAAACCACACGAAAATTAAAAATGGTC | 0 |
| | Plant #2 | GTAATTTGATGAAGACAGATCTGAAAACCACACGAAAATTAAAAATGGTC | 0 |
| | Plant #3 | GTAATTTGATGAAGACAGATCTGAAAACCACACGAAAATTAAAAATGGTC | 0 |
| | Plant #4 | GTAATTTGATGAAGACAGATCTGAAAACCACACGAAAATTAAAAATGGTC | 0 |
| | Plant #5 | GTAATTTGATGAAGACAGATCTGAAAACCACACGAAAATTAAAAATGGTC | 0 |
| ChlH.3off | Target | TTCGTATTTACCATAAGTTA**GGGTTTTCAGATCTCTTTTA****ATCAA**ACCAA | |
| | Wild-type | TTCGTATTTACCATAAGTTAGGGTTTTCAGATCTCTTTTAATCAAACCAA | |
| | Plant #1 | TTCGTATTTACCATAAGTTAGGGTTTTCAGATCTCTTTTAATCAAACCAA | 0 |
| | Plant #2 | TTCGTATTTACCATAAGTTAGGGTTTTCAGATCTCTTTTAATCAAACCAA | 0 |
| | Plant #3 | TTCGTATTTACCATAAGTTAGGGTTTTCAGATCTCTTTTAATCAAACCAA | 0 |
| | Plant #4 | TTCGTATTTACCATAAGTTAGGGTTTTCAGATCTCTTTTAATCAAACCAA | 0 |
| | Plant #5 | TTCGTATTTACCATAAGTTAGGGTTTTCAGATCTCTTTTAATCAAACCAA | 0 |
| ChlH.4off | Target | ACTTGAATTGTAACTTACCA**AAATATACAGAGATCTCTTC****ATCAA**AAAAG | |
| | Wild-type** | ACTTGAATTGTAACTTACCAAAATATACAGAGATCTCTTCATCAAAAAAG | |
| | Plant #1** | ACTTGAATTGTAACTTACCAAAATATACAGAGATCTCTTCATCAAAAAAG | 0 |
| | Plant #2** | ACTTGAATTGTAACTTACCAAAATATACAGAGATCTCTTCATCAAAAAAG | 0 |
| | Plant #3** | ACTTGAATTGTAACTTACCAAAATATACAGAGATCTCTTCATCAAAAAAG | 0 |
| | Plant #4** | ACTTGAATTGTAACTTACCAAAATATACAGAGATCTCTTCATCAAAAAAG | 0 |
| | Plant #5** | ACTTGAATTGTAACTTACCAAAATATACAGAGATCTCTTCATCAAAAAAG | 0 |
| ChlH.5off | Target | AAGAC**TTGAT****GAAGAGATTTCTGTTTAGCC**AGGCCAGTAGGGTCGCCATC | |
| | Wild-type | AAGACTTGATGAAGAGATTTCTGTTTAGCCAGGCCAGTAGGGTCGCCATC | |
| | Plant #1 | AAGACTTGATGAAGAGATTTCTGTTTAGCCAGGCCAGTAGGGTCGCCATC | 0 |
| | Plant #2 | AAGACTTGATGAAGAGATTTCTGTTTAGCCAGGCCAGTAGGGTCGCCATC | 0 |
| | Plant #3 | AAGACTTGATGAAGAGATTTCTGTTTAGCCAGGCCAGTAGGGTCGCCATC | 0 |
| | Plant #4 | AAGACTTGATGAAGAGATTTCTGTTTAGCCAGGCCAGTAGGGTCGCCATC | 0 |
| | Plant #5 | AAGACTTGATGAAGAGATTTCTGTTTAGCCAGGCCAGTAGGGTCGCCATC | 0 |
| ChlH.6off | Target | TGGTG**TTGAT****GAAGAGTTCGCAAAACTGCCC**TGAGGTGATCGGCATGGTC | |
| | Wild-type | TGGTGTTGATGAAGAGTTCGCAAAACTGCCCTGAGGTGATCGGCATGGTC | |
| | Plant #1 | TGGTGTTGATGAAGAGTTCGCAAAACTGCCCTGAGGTGATCGGCATGGTC | 0 |
| | Plant #2 | TGGTGTTGATGAAGAGTTCGCAAAACTGCCCTGAGGTGATCGGCATGGTC | 0 |
| | Plant #3 | TGGTGTTGATGAAGAGTTCGCAAAACTGCCCTGAGGTGATCGGCATGGTC | 0 |
| | Plant #4 | TGGTGTTGATGAAGAGTTCGCAAAACTGCCCTGAGGTGATCGGCATGGTC | 0 |
| | Plant #5 | TGGTGTTGATGAAGAGTTCGCAAAACTGCCCTGAGGTGATCGGCATGGTC | 0 |

*Bold red letters indicate TAM, and bold underlined letters denote reRNA target sequence

**Supplementary Table. 23.** Targets used in the paper

| | |
|---|---|
| reRNA1$^{PDS}$ | TATCCAAGACCGGAGCTAGA |
| reRNA2$^{PDS}$ | TTTCCTGAAGCTCTTCCTGC |
| reRNA3$^{PDS}$ | TGCTTTGAACAGATTTCTTC |
| reRNA4$^{PDS}$ | GCTACAATGAAGGAACTAGC |
| reRNA$^{ChlH}$ | GAAGAGATCTCTGAACACCC |

**Supplementary Table 24.** Primers used for generating vectors, RT-PCR, and qPCR

| Primer # | Primer sequence (5' to 3') |
|---|---|
| 100131 | TGATAcccgggTGCTTCCGGCGGGGCTCGAACCCG |
| 101572 | atTCCGgaTCCcatgcatggactacgtgggtactttttttgaacacgtGCCAGAACCctgatgtgagacatttgtgttactccatttag |
| 103919 | ATCtctagaccATGATACGCAACAAGGCCTTTGTTG |
| 103920 | cagTCGCGAcgcgTCAcactttcctcttcttcttagg |
| 103921 | ATCtctagacgcgtGGTGGCTGCGGGAATCTCAGACAC |
| 103922 | GTAcccgggAGCTCTGGTCTTGGATTTGAACCT |
| 103923 | GTAcccgggagctcGAAGAGCTTCAGGAAATTGAACCT |
| 103925 | ATCggtaccGGTGGCTGCGGGAATCTCAGACAC |
| 104172 | GTAcccgggagctcAAATCTGTTCAAAGCATTGAACCTCACACGACTAAAGTCG |
| 104173 | GTAcccgggagctcGTTCCTTCATTGTAGCTTGAACCTCACACGACTAAAGTCG |
| 104178 | atcTCTAGACCATGGCTCCTAAGAAGA |
| 104180 | TCCTGCGGTTCacgcgtTTACTTC |
| 104181 | ATCacgcgtGAACCGCAGGAATTGCGG |
| 104200 | atcGGTACCttcttgaatcctgatgag |
| 104205 | atcTCTAGAttcttgaatcctgatgagtccgtg |
| 104230 | GTAgagctcGTCCCATTCGCCATGCCGAAGCAT |
| 104361 | ATCggtaccGAACCGCAGGAATTGCGGGGGTAG |
| 104365 | ATCtctagaATGATACGCtatAAGGCA |
| 104366 | GTAggtaccCTAGTTTGTATAAGCCTG |
| 104367 | ATCtctagaATGATTAGAtatAAGGCT |
| 104368 | GTAggtaccTCAGTTTGTATAAGCCTG |
| 104452 | ATCtctagaGGTGGCTGCGGGAATCTCAGACAC |
| 104534 | atctctagaATGATACGCtatAAGGCATTTGTG |
| 104639 | ATCtctagaATGATACGCaatAAGGCA |
| 104640 | TGAggtaccCTAGTTTGTATAAGCCTG |
| 104641 | TGAggtaccTCAcactttcctcttcttcttagggtcttccacctttcttttctttttagggtcggccctgctCACTACGAGAGTAGCG |
| 104717 | gaaaatcacgcgacactggttgtgagcagggccgacGAAGGTTCAGGGA |
| 104718 | ATCtctagaatgatacgcTATaaggcc |
| 104729 | GTATCACTTACCCGAGTTAACGAGTCTAGAATGATACGCAACAAGGCCTTTGTT |
| 104730 | CACAACCAGTGTCGCGTGATTTTC |
| 104930 | ataTCTAGAatgttacaacacaaggcttacgaa |
| 104931 | ataGGTACCtcatcacactttcctcttcttctt |
| 105425 | GACATCATGACATTATCAATTGAC |
| 105426 | ACCTGTAATGGGCTCTGAAATAA |
| 105432 | CGAAGTTAGAGATTTGTGGAAAGC |
| 105433 | AGACGAGCTGATGAGAAGAATG |
| 105434 | GTGTTCGCCTTGGTAGTAGTAG |
| 105443 | CGCTAGATATGGCAAGGCTAAG |
| 105444 | CAGGTGTTCAGTCCCGATTATT |
| 105448 | GACAAATTCGCCTTGCAGAATAG |
| 105449 | ATTGGTAAATTGGGTACGGTAAGA |

**Supplementary Table 25.** Constructs generated and used in the study

| Construct ID | Description | Included in | Addgene # |
|---|---|---|---|
| 4927 | TRV2-TnpB-reRNA4$^{PDS}$ | Extended Data Fig. 4a | Will deposit |
| 4928 | TRV2-TnpB-$^{HH}$reRNA4$^{PDS}$ | Fig. 1b | Will deposit |
| 4784 | TRV2-eTnpBc-reRNA4$^{PDS}$ | Extended Data Fig. 4a | Will deposit |
| 4787 | TRV2-eTnpBc-$^{HH}$reRNA4$^{PDS}$ | Fig. 1b | Will deposit |
| 4954 | TRV2-eTnpBe-reRNA4$^{PDS}$ | Extended Data Fig. 4a | Will deposit |
| 4955 | TRV2-eTnpBe-$^{HH}$reRNA4$^{PDS}$ | Extended Data Fig. 4a | Will deposit |
| 4654 | TRV2-NbCoTnpB$_{2xSV40nls}$ | Extended Data Fig. 1a | |
| 4655 | TRV2-reRNA1$^{PDS}$ | Extended Data Fig. 1a | |
| 4657 | TRV2-reRNA2$^{PDS}$ | Extended Data Fig. 1a | |
| 4742 | TRV2-reRNA3$^{PDS}$ | Extended Data Fig. 1a | |
| 4743 | TRV2-reRNA4$^{PDS}$ | Extended Data Fig. 1a | |
| 4984 | TRV2-TnpB$_{FDnls}$ | Extended Data Fig. 3a | |
| 4949 | TRV2-eTnpBc$_{FDnls}$ | Extended Data Fig. 3a | |
| 4985 | TRV2-eTnpBe$_{FDnls}$ | Extended Data Fig. 3a | |
| 4950 | TRV2-reRNA4$^{PDS}$ | Extended Data Fig. 3a | |
| 4951 | TRV2-$^{HH}$reRNA4$^{PDS}$ | Extended Data Fig. 3a | |
| 4947 | TRV2-NbCoTnpB-reRNA4$^{PDS}$ | Extended Data Fig. 6a | |
| 4948 | TRV2-NbCoTnpB-$^{HH}$reRNA4$^{PDS}$ | Extended Data Fig. 6a | |
| 4981 | TRV2-NbCoeTnpBc-reRNA4$^{PDS}$ | Extended Data Fig. 6a | |
| 4982 | TRV2-NbCoeTnpBc-$^{HH}$reRNA4$^{PDS}$ | Extended Data Fig. 6a | |
| 4945 | TRV2-NbCoeTnpBe-reRNA4$^{PDS}$ | Extended Data Fig. 6a | |
| 4946 | TRV2-NbCoeTnpBe-$^{HH}$reRNA4$^{PDS}$ | Extended Data Fig. 6a | |
| 4788 | TRV2-NtCoeTnpBc-reRNA2$^{PDS}$ | Extended Data Fig. 7a | |
| 4791 | TRV2-NtCoeTnpBc-$^{HH}$reRNA2$^{PDS}$ | Extended Data Fig. 7a | |
| 4789 | TRV2-NtCoeTnpBc-reRNA3$^{PDS}$ | Extended Data Fig. 7a | |
| 4792 | TRV2-NtCoeTnpBc-$^{HH}$reRNA3$^{PDS}$ | Extended Data Fig. 7a | |
| 4790 | TRV2-NtCoeTnpBc-reRNA4$^{PDS}$ | Extended Data Fig. 7a | |
| 4793 | TRV2-NtCoeTnpBc-$^{HH}$reRNA4$^{PDS}$ | Extended Data Fig. 7a | |
| 4931 | TRV2-TnpB$_{2xSV40nls}$-reRNA4$^{PDS}$ | Extended Data Fig. 5a | Will deposit |
| 4932 | TRV2-TnpB$_{2xSV40nls}$-$^{HH}$reRNA4$^{PDS}$ | Extended Data Fig. 5a | Will deposit |
| 4929 | TRV2-eTnpBc$_{2xSV40nls}$-reRNA4$^{PDS}$ | Extended Data Fig. 5a | Will deposit |
| 4930 | TRV2-eTnpBc$_{2xSV40nls}$-$^{HH}$reRNA4$^{PDS}$ | Extended Data Fig. 5a | Will deposit |
| 4756 | TRV2-NtcoISYmu1-reRNA4$^{PDS}$ | Extended Data Fig. 7a | |
| 5034 | TRV2-HscoISYmu1-reRNA4$^{PDS}$ | Extended Data Fig. 7a | |
| 5035 | TRV2-ISYmu1$^{L167G/V305R}$-reRNA4$^{PDS}$ | Extended Data Fig. 7d | |
| 5036 | TRV2-ISYmu1$^{H4Y/V305R}$-reRNA4$^{PDS}$ | Extended Data Fig. 7e | |
| 5166 | TRV2-AtcoISYmu1-reRNA4$^{PDS}$ | Extended Data Fig. 7c | |

**Supplementary Table 26.** Primers used for amplicon generation

| Purpose | Primer # | Sequence | Barcodes |
|---|---|---|---|
| NGS of amplicons of reRNA1$^{PDS}$ | 105450 | CTTTCCCTACACGACGCTCTTCCGATCTaacgtgatGTTATGTTTTGGTAGTAGCGACTC | AACGTGAT |
| | 105451 | ggagttcagacgtgtgctcttccgatctCGATGTTTaccagcaataacaatctccaatg | AAACATCG |
| | 105452 | CTTTCCCTACACGACGCTCTTCCGATCTatgcctaaGTTATGTTTTGGTAGTAGCGACTC | ATGCCTAA |
| | 105453 | ggagttcagacgtgtgctcttccgatctTGACCACTaccagcaataacaatctccaatg | AGTGGTCA |
| | 105454 | CTTTCCCTACACGACGCTCTTCCGATCTaccactgtGTTATGTTTTGGTAGTAGCGACTC | ACCACTGT |
| | 105455 | ggagttcagacgtgtgctcttccgatctGCCAATGTaccagcaataacaatctccaatg | ACATTGGC |
| | 105456 | CTTTCCCTACACGACGCTCTTCCGATCTcagatctgGTTATGTTTTGGTAGTAGCGACTC | CAGATCTG |
| | 105457 | ggagttcagacgtgtgctcttccgatctACTTGATGaccagcaataacaatctccaatg | CATCAAGT |
| NGS of amplicons of reRNA2$^{PDS}$ | 104626 | ACACTCTTTCCCTACACGACGCTCTTCCGATCTaacgtgatATATGCAGAACCTGTTTGGAGAAC | AACGTGAT |
| | 104627 | gactggagttcagacgtgtgctcttccgatctCGATGTTTagcatttcgttgttctttagtatg | AAACATCG |
| | 104628 | ACACTCTTTCCCTACACGACGCTCTTCCGATCTatgcctaaATATGCAGAACCTGTTTGGAGAAC | ATGCCTAA |
| | 104629 | gactggagttcagacgtgtgctcttccgatctTGACCACTagcatttcgttgttctttagtatg | AGTGGTCA |
| | 104630 | ACACTCTTTCCCTACACGACGCTCTTCCGATCTaccactgtATATGCAGAACCTGTTTGGAGAAC | ACCACTGT |
| | 104631 | gactggagttcagacgtgtgctcttccgatctGCCAATGTagcatttcgttgttctttagtatg | ACATTGGC |
| | 104632 | ACACTCTTTCCCTACACGACGCTCTTCCGATCTcagatctgATATGCAGAACCTGTTTGGAGAAC | CAGATCTG |
| | 104633 | gactggagttcagacgtgtgctcttccgatctACTTGATGagcatttcgttgttctttagtatg | CATCAAGT |
| | 105458 | CTTTCCCTACACGACGCTCTTCCGATCTcgctgatcCAAATATGCAGAACCTGTTTGGA | CGCTGATC |
| | 105459 | ggagttcagacgtgtgctcttccgatctTAGCTTGTctacatgcagaaattctgatcaag | ACAAGCTA |
| | 105460 | CTTTCCCTACACGACGCTCTTCCGATCTctgtagccCAAATATGCAGAACCTGTTTGGA | CTGTAGCC |
| | 105461 | ggagttcagacgtgtgctcttccgatctCTTGTACTctacatgcagaaattctgatcaag | AGTACAAG |
| | 105462 | CTTTCCCTACACGACGCTCTTCCGATCTaacaaccaCAAATATGCAGAACCTGTTTGGA | AACAACCA |
| | 105463 | ggagttcagacgtgtgctcttccgatctTCTCGGTTctacatgcagaaattctgatcaag | AACCGAGA |
| | 105464 | CTTTCCCTACACGACGCTCTTCCGATCTaacgcttaCAAATATGCAGAACCTGTTTGGA | AACGCTTA |
| | 105465 | ggagttcagacgtgtgctcttccgatctTCCGTCTTctacatgcagaaattctgatcaag | AAGACGGA |
| NGS of amplicons of reRNA3$^{PDS}$ | 104594 | ACACTCTTTCCCTACACGACGCTCTTCCGATCTaacgtgatTCAGGGTGTGCCTGATAGGGTGAC | AACGTGAT |
| | 104595 | gactggagttcagacgtgtgctcttccgatctCGATGTTTcatttttgaaccatgtttctcctg | AAACATCG |
| | 104596 | ACACTCTTTCCCTACACGACGCTCTTCCGATCTatgcctaaTCAGGGTGTGCCTGATAGGGTGAC | ATGCCTAA |
| | 104597 | gactggagttcagacgtgtgctcttccgatctTGACCACTcatttttgaaccatgtttctcctg | AGTGGTCA |
| | 104598 | ACACTCTTTCCCTACACGACGCTCTTCCGATCTaccactgtTCAGGGTGTGCCTGATAGGGTGAC | ACCACTGT |
| | 104599 | gactggagttcagacgtgtgctcttccgatctGCCAATGTcatttttgaaccatgtttctcctg | ACATTGGC |
| | 104600 | ACACTCTTTCCCTACACGACGCTCTTCCGATCTcagatctgTCAGGGTGTGCCTGATAGGGTGAC | CAGATCTG |
| | 104601 | gactggagttcagacgtgtgctcttccgatctACTTGATGcatttttgaaccatgtttctcctg | CATCAAGT |
| | 104602 | ACACTCTTTCCCTACACGACGCTCTTCCGATCTcgctgatcTCAGGGTGTGCCTGATAGGGTGAC | CGCTGATC |
| | 104603 | gactggagttcagacgtgtgctcttccgatctTAGCTTGTcatttttgaaccatgtttctcctg | ACAAGCTA |
| | 104604 | ACACTCTTTCCCTACACGACGCTCTTCCGATCTctgtagccTCAGGGTGTGCCTGATAGGGTGAC | CTGTAGCC |
| | 104605 | gactggagttcagacgtgtgctcttccgatctCTTGTACTcatttttgaaccatgtttctcctg | AGTACAAG |
| | 104606 | ACACTCTTTCCCTACACGACGCTCTTCCGATCTaacaaccaTCAGGGTGTGCCTGATAGGGTGAC | AACAACCA |
| | 104607 | gactggagttcagacgtgtgctcttccgatctTCTCGGTTcatttttgaaccatgtttctcctg | AACCGAGA |
| | 104608 | ACACTCTTTCCCTACACGACGCTCTTCCGATCTaacgcttaTCAGGGTGTGCCTGATAGGGTGAC | AACGCTTA |
| | 104609 | gactggagttcagacgtgtgctcttccgatctTCCGTCTTcatttttgaaccatgtttctcctg | AAGACGGA |
| | 105466 | CTTTCCCTACACGACGCTCTTCCGATCTaacgtgatATCTCAGCAGCAGCTATTTGCTTA | AACGTGAT |
| | 105467 | ggagttcagacgtgtgctcttccgatctCGATGTTTcaggtccctaacatgatttacagc | AAACATCG |
| | 105468 | CTTTCCCTACACGACGCTCTTCCGATCTatgcctaaATCTCAGCAGCAGCTATTTGCTTA | ATGCCTAA |
| | 105469 | ggagttcagacgtgtgctcttccgatctTGACCACTcaggtccctaacatgatttacagc | AGTGGTCA |
| | 105470 | CTTTCCCTACACGACGCTCTTCCGATCTaccactgtATCTCAGCAGCAGCTATTTGCTTA | ACCACTGT |
| | 105471 | ggagttcagacgtgtgctcttccgatctGCCAATGTcaggtccctaacatgatttacagc | ACATTGGC |
| | 105472 | CTTTCCCTACACGACGCTCTTCCGATCTcagatctgATCTCAGCAGCAGCTATTTGCTTA | CAGATCTG |
| | 105473 | ggagttcagacgtgtgctcttccgatctACTTGATGcaggtccctaacatgatttacagc | CATCAAGT |
| NGS of amplicons of reRNA4$^{PDS}$ | 104610 | ACACTCTTTCCCTACACGACGCTCTTCCGATCTaacgtgatGTCTCTCTGCAGGAATATTACAAC | AACGTGAT |
| | 104611 | gactggagttcagacgtgtgctcttccgatctCGATGTTTggcacagttttataaacagacctg | AAACATCG |
| | 104612 | ACACTCTTTCCCTACACGACGCTCTTCCGATCTatgcctaaGTCTCTCTGCAGGAATATTACAAC | ATGCCTAA |
| | 104613 | gactggagttcagacgtgtgctcttccgatctTGACCACTggcacagttttataaacagacctg | AGTGGTCA |
| | 104614 | ACACTCTTTCCCTACACGACGCTCTTCCGATCTaccactgtGTCTCTCTGCAGGAATATTACAAC | ACCACTGT |
| | 104615 | gactggagttcagacgtgtgctcttccgatctGCCAATGTggcacagttttataaacagacctg | ACATTGGC |
| | 104616 | ACACTCTTTCCCTACACGACGCTCTTCCGATCTcagatctgGTCTCTCTGCAGGAATATTACAAC | CAGATCTG |
| | 104617 | gactggagttcagacgtgtgctcttccgatctACTTGATGggcacagttttataaacagacctg | CATCAAGT |
| | 104618 | ACACTCTTTCCCTACACGACGCTCTTCCGATCTcgctgatcGTCTCTCTGCAGGAATATTACAAC | CGCTGATC |
| | 104619 | gactggagttcagacgtgtgctcttccgatctTAGCTTGTggcacagttttataaacagacctg | ACAAGCTA |
| | 104620 | ACACTCTTTCCCTACACGACGCTCTTCCGATCTctgtagccGTCTCTCTGCAGGAATATTACAAC | CTGTAGCC |
| | 104621 | gactggagttcagacgtgtgctcttccgatctCTTGTACTggcacagttttataaacagacctg | AGTACAAG |
| | 104622 | ACACTCTTTCCCTACACGACGCTCTTCCGATCTaacaaccaGTCTCTCTGCAGGAATATTACAAC | AACAACCA |
| | 104623 | gactggagttcagacgtgtgctcttccgatctTCTCGGTTggcacagttttataaacagacctg | AACCGAGA |

| | | | |
|---|---|---|---|
| | 104624 | ACACTCTTTCCCTACACGACGCTCTTCCGATCTaacgcttaGTCTCTCTGCAGGAATATTACAAC | AACGCTTA |
| | 104625 | gactggagttcagacgtgtgctcttccgatctTCCGTCTTggcacagttttataaacagacctg | AAGACGGA |
| | 105206 | ACACTCTTTCCCTACACGACGCTCTTCCGATCTaacgtgatGAGTGAGTGGGGAGTAAAATTA | AACGTGAT |
| | 105207 | gactggagttcagacgtgtgctcttccgatctCGATGTTTaaagcagattaatcaggattca | AAACATCG |
| | 105208 | ACACTCTTTCCCTACACGACGCTCTTCCGATCTatgcctaaGAGTGAGTGGGGAGTAAAATTA | ATGCCTAA |
| | 105209 | gactggagttcagacgtgtgctcttccgatctTGACCACTaaagcagattaatcaggattca | AGTGGTCA |
| | 105210 | ACACTCTTTCCCTACACGACGCTCTTCCGATCTaccactgtGAGTGAGTGGGGAGTAAAATTA | ACCACTGT |
| | 105211 | gactggagttcagacgtgtgctcttccgatctGCCAATGTaaagcagattaatcaggattca | ACATTGGC |
| | 105212 | ACACTCTTTCCCTACACGACGCTCTTCCGATCTcagatctgGAGTGAGTGGGGAGTAAAATTA | CAGATCTG |
| | 105213 | gactggagttcagacgtgtgctcttccgatctACTTGATGaaagcagattaatcaggattca | CATCAAGT |
| NGS of amplicons of reRNA$^{ChlH}$ | 105065 | ACACTCTTTCCCTACACGACGCTCTTCCGATCTaacgtgatGGTGATGTGGATGATTGGTGTTAG | AACGTGAT |
| | 105066 | gactggagttcagacgtgtgctcttccgatctCGATGTTTactgctcggtcaaggagattcatc | AAACATCG |
| | 105067 | ACACTCTTTCCCTACACGACGCTCTTCCGATCTatgcctaaGGTGATGTGGATGATTGGTGTTAG | ATGCCTAA |
| | 105068 | gactggagttcagacgtgtgctcttccgatctTGACCACTactgctcggtcaaggagattcatc | AGTGGTCA |
| | 105069 | ACACTCTTTCCCTACACGACGCTCTTCCGATCTaccactgtGGTGATGTGGATGATTGGTGTTAG | ACCACTGT |
| | 105070 | gactggagttcagacgtgtgctcttccgatctGCCAATGTactgctcggtcaaggagattcatc | ACATTGGC |
| | 105071 | ACACTCTTTCCCTACACGACGCTCTTCCGATCTcagatctgGGTGATGTGGATGATTGGTGTTAG | CAGATCTG |
| | 105072 | gactggagttcagacgtgtgctcttccgatctACTTGATGactgctcggtcaaggagattcatc | CATCAAGT |

**Supplementary Table 27.** Primers used to assess off-targets

| Primer # | Primer sequence (5' to 3') | |
|---|---|---|
| 105485 | CTTTCCCTACACGACGCTCTTCCGATCTaacgtgatCAGGCAATGTAGACCTGTCATC | UP to PCR PDS4.1off |
| 105486 | ggagttcagacgtgtgctcttccgatctCGATGTTTgaccagcagttgttggtacata | DS to PCR PDS4.1off |
| 105487 | CTTTCCCTACACGACGCTCTTCCGATCTatgcctaaGAGGATAATTGAGTTCCTGAAGGG | UP to PCR PDS4.2off |
| 105488 | ggagttcagacgtgtgctcttccgatctTGACCACTgagcatctgctttgagggtaaa | DS to PCR PDS4.2off |
| 105489 | CTTTCCCTACACGACGCTCTTCCGATCTaccactgtGACACTTTCCCTTTACCTCTACTT | UP to PCR PDS4.3off |
| 105490 | ggagttcagacgtgtgctcttccgatctGCCAATGTagcaccgcaaatgtaatcttattc | DS to PCR PDS4.3off |
| 105491 | CTTTCCCTACACGACGCTCTTCCGATCTcagatctgGGACCAAACTCTTGTTGGTAGT | UP to PCR PDS4.4off |
| 105492 | gagttcagacgtgtgctcttccgatctACTTGATGcaatttcaaaggttaggcgtgatag | DS to PCR PDS4.4off |
| 105493 | CTTTCCCTACACGACGCTCTTCCGATCTcgctgatcGCAGTTCTATCGGACAAGAGAA | UP to PCR PDS4.5off |
| 105494 | gagttcagacgtgtgctcttccgatctTAGCTTGTatggcatcaaaaagacatatattgt | DS to PCR PDS4.5off |
| 105495 | CTTTCCCTACACGACGCTCTTCCGATCTctgtagccAATAATTTCGCACGCTTGACTC | UP to PCR PDS4.6off |
| 105496 | ggagttcagacgtgtgctcttccgatctCTTGTACTacatgaatataccagtgagctgat | DS to PCR PDS4.6off |
| 105497 | CTTTCCCTACACGACGCTCTTCCGATCTaacaaccaGGAAGTATTTCAAGAGATGCCAAG | UP to PCR PDS4.7off |
| 105498 | ggagttcagacgtgtgctcttccgatctTCTCGGTTtcaagtcgattctagctccattat | DS to PCR PDS4.7off |
| 105503 | TTTCCCTACACGACGCTCTTCCGATCTatgcctaaGCCATAACAAAGAACATAGCCATAG | UP to PCR ChlHt.1off |
| 105504 | gagttcagacgtgtgctcttccgatctTGACCACTtgcaatttagtgattaagggttgtc | DS to PCR ChlHt.1off |
| 105505 | CTTTCCCTACACGACGCTCTTCCGATCTaccactgtCTCTCCACATAGTAAGGCATTAGG | UP to PCR ChlHt.2off |
| 105506 | ggagttcagacgtgtgctcttccgatctGCCAATGTgtttgatctgtatacccggagaag | DS to PCR ChlHt.2off |
| 105507 | CTTTCCCTACACGACGCTCTTCCGATCTcagatctgCAGCATCTTCAACAAACAAATGTC | UP to PCR ChlHt.3off |
| 105508 | gttcagacgtgtgctcttccgatctACTTGATGacaataccacaagatatatagagacac | DS to PCR ChlHt.3off |
| 105509 | CTTTCCCTACACGACGCTCTTCCGATCTcgctgatcCCTGTTTGGTCAGAGGTATATCG | UP to PCR ChlHt.4off |
| 105510 | ggagttcagacgtgtgctcttccgatctTAGCTTGTgctagacaagtaaggtcacatagg | DS to PCR ChlHt.4off |
| 105511 | CTTTCCCTACACGACGCTCTTCCGATCTctgtagccAGGACATCATAAGCAGTTGTCATA | UP to PCR ChlHt.5off |
| 105512 | ggagttcagacgtgtgctcttccgatctCTTGTACTtactgacgtgcgagatcaataaa | DS to PCR ChlHt.5off |
| 105515 | CTTTCCCTACACGACGCTCTTCCGATCTaacgcttaGCCAATCCTTCAATGCTTCTATC | UP to PCR ChlHt.6off |
| 105516 | ggagttcagacgtgtgctcttccgatctTCCGTCTTtcacaacctgctatggttactt | DS to PCR ChlHt.6off |
| 105517 | CAGGCAATGTAGACCTGTCATC | to seq PDS4.1off |
| 105518 | GAGGATAATTGAGTTCCTGAAGGG | to seq PDS4.2off |
| 105519 | GACACTTTCCCTTTACCTCTACTT | to seq PDS4.3off |
| 105520 | GGACCAAACTCTTGTTGGTAGT | to seq PDS4.4off |
| 105521 | GCAGTTCTATCGGACAAGAGAA | to seq PDS4.5off |
| 105522 | AATAATTTCGCACGCTTGACTC | to seq PDS4.5off |
| 105523 | GGAAGTATTTCAAGAGATGCCAAG | to seq PDS4.7off |
| 105524 | GGAAGTATTTCAAGAGATGCCAAG | to seq PDS4.8off |
| 105526 | GCCATAACAAAGAACATAGCCATAG | to seq ChlH.1off |
| 105527 | CTCTCCACATAGTAAGGCATTAGG | to seq ChlH.2off |
| 105528 | CAGCATCTTCAACAAACAAATGTC | to seq ChlH.3off |
| 105529 | CCTGTTTGGTCAGAGGTATATCG | to seq ChlH.4off |
| 105530 | GGACATCATAAGCAGTTGTCATA | to seq ChlH.5off |
| 105532 | GCCAATCCTTCAATGCTTCTATC | to seq ChlH.6off |
